# Beyond Blast: Enabling Microbiologists to Better Extract Literature, Taxonomic Distributions and Gene Neighborhood Information for Protein Families

**DOI:** 10.1101/2023.05.03.539116

**Authors:** Colbie J. Reed, Rémi Denise, Jacob Hourihan, Jill Babor, Marshall Jaroch, Maria Martinelli, Geoffrey Hutinet, Valérie de Crécy-Lagard

## Abstract

Capturing the published corpus of information on all members of a given protein family should be an essential step in any study focusing on specific members of that said family. Using a previously gathered dataset of more than 280 references mentioning a member of the DUF34 (NIF3/Ngg1-interacting Factor 3), we evaluated the efficiency of different databases and search tools, and devised a workflow that experimentalists can use to capture the most published information on members of a protein family in the least amount of time. To complement this workflow, web-based platforms allowing for the exploration of protein family members across sequenced genomes or for the analysis of gene neighborhood information were reviewed for their versatility and ease of use. Recommendations that can be used for experimentalist users, as well as educators, are provided and integrated within a customized, publicly accessible Wiki.

**Data summary:** The authors confirm all supporting data, code, and protocols have been provided within the article or through supplementary data files. Complete set of supplementary data sheets may be accessed via FigShare.

## Introduction

In the last 35 years, the field of microbiology has undergone a total revolution. The completion of the first whole genome sequence of a bacterium, *Haemophilus influenzae* RD40 in 1995 [1], changed the way bench scientists design and/or interpret their experiments: the analysis of sequences (gene, protein, whole genomes) has become an integral part of the whole process [2]. This led to the incredible success of the BLAST suite developed at NCBI by Altschul et al. [3] that allowed any scientist with an internet connection to ask whether his/her favorite gene/protein was similar to an already experimentally characterized one or whether a similar sequence was present in particular organisms. From 1995 to 2005, most microbiologists could get by with NCBI and cloning design platforms as their bioinformatic toolboxes. The arrival of Next Generation Sequencing (NGS) technologies has made the sequencing of microbial genomes a routine procedure. Today, this technological advancement is feeding thousands of microbial genomes and metagenomes into GenBank [4] every week (or even every day) thus transforming many fields of microbiology, from ecology [5] to food microbiology [6], infectious diseases [7] and basic enzymology [8]. This ‘deluge of data’ [9] is making simple BLAST searches useless for most applications as, without specific filters, BLAST will just retrieve hundreds of sequences closely related to the input sequence. In an ideal world, every biologist would be trained in using command line and programming tools that would allow them to cope with this encumbrance of data [10]. This might be the case in a few years’ time, but such a solution has yet to be realized and many researchers are likely to be left behind due to resource, access, and opportunity constraints. Fortunately, a plethora of databases have developed various programs with web-accessible Graphical User Interfaces (GUIs) that allow users with little to no programming experience to take full advantage of the information possible to be derived from the over 250K available complete microbial genome sequences [11].

Integrated microbial genome portals (e.g., MicrobesOnline, JGI-IMG) are the easiest entry points for accessing and analyzing data derived from microbial genomes. Many microbiologists become aware of these resources only when they need to annotate a genome sequenced in their own laboratories, as most offer user-friendly annotation pipelines [12–15]. These microbial genome web-portals are quite versatile and offer various tools that were recently extensively reviewed in a side-by-side comparison [16]. Some databases offer training through introductory workshops, which can be great gateways into the available resources, yet these tend to reach only a small audience and are often restricted to a specific platform. Tutorials are also available but—in our experience teaching the use of web-based tools to undergraduate, graduate, and post-graduate audiences, both, in formal classes and in workshops—we find that these are most useful when used to “refresh” the skills of seasoned users instead of being used to get a novice user started.

We have been using comparative genomic driven approaches using only web-based tools to link genes and functions for over 20 years, leading to the functional characterization of more than 65 gene families (**Table S1**). This work required the use of all the available microbial genome web-portals, learning the strengths and weaknesses of each in the process. Here, we address problems that routinely arise for experimentalists interested in a specific protein family and show how they can be resolved using the web-portals as well as more specialized online tools. We focus on answering three specific questions. First, “what information has already been published for any member of a protein family?” Second, “how can one best analyze and visualize the taxonomic distribution for members of a protein family?” Finally, “how can physical clustering data for genes of a given family be gathered and visualized?” In answering these questions, we intend to showcase the different microbial web-portals, as well as identify and discuss their limitations. Additionally, we present a resource targeted towards novice bioinformatic tool users, the VDC-Lab Wiki, that compiles databases that we routinely use for research and teaching, doing so with an informed curatorial eye guided by 20 years of experience in navigating biological databases.

## Methods

### Protein Family Case Study and Literature Review, Curation

Process of retrieval described in detail in text; resulting accumulation of published keywords, identifiers and accessions is provided (Data S1). A list of tools, databases, and search engines were compiled for use in and as a result of this work. The totality of these resources can be reviewed in the provided supplemental materials (Data S2). Venn diagrams were generated using the online bioinformatics tools of Gent University (https://bioinformatics.psb.ugent.be/webtools/Venn/).

### Data Analysis, Figure Generation

Microsoft Office Excel (Office16) was used for tallying observations, query results, in addition to documenting the curation process and generating figures of curation results. Other figures and diagrams were created using Microsoft Office PowerPoint.

### Wiki website development and publishing, gathering and curating resource information

The websites featured on the wiki originate from a list of websites amassed by Valérie de Crécy-Lagard over time, as well as from discoveries made by laboratory members throughout their research. The included websites were tested by at least one lab member, who then crafted a brief description for it. Subsequently, all the websites were categorized and incorporated into the wiki. The VDC lab wiki can be found at the following address: https://vdclab-wiki.herokuapp.com/

## Results

### Investigating workflows for capturing literature for all members of a protein family

Comprehensively identifying literature pertaining to all members of a given protein family for the purposes of background review or hypothesis generation is often the first step in many biological studies. This task remains rife with challenges in an era defined by massive accumulations of biological data [9,17,18]. Most microbiologists depend on PubMed [19] to find literature, relying on its text-based search tools. Although efforts by the scientific community have been made within the last decade to popularize adherence to uniform data standards that prioritize the findability, <u>a</u>ccessibility interoperability and reusability (or ‘FAIR’) of information [20], these principles have yet to be systematically implemented among databases and publishing journals [21], and, as a result, the state of linking publications to the biomolecular entities (genes or proteins) that they describe remains suboptimal. The only journal to-date that has imposed such a standard is *Biochemistry*; since 2018, it has required authors to complete a form providing UniProt entry information for the proteins described in the paper being submitted [22,23].

To both explore the challenges of finding all relevant literature of a protein family and propose potential solutions, a stepwise demonstration of the capture process was recapitulated using the conserved unknown protein family, DUF34, recently examined in Reed et al. [24]. In this case study, publications were classified as being either “focal” (i.e., any family homolog being mentioned in the title or abstract) or “non-focal” (i.e., any family homolog being mentioned anywhere outside of the abstract or title, including supplementary materials). Additionally, “false positives” (i.e., resulting papers found to lack relevance to any DUF34 family member) were also flagged. The DUF34 family was selected for its high level of conservation across all three domains of life, its described within-family diversity, and its variable, pleiotropic functional associations observed between study organisms. This family was used as a shared target for different methods of literature capture described and compared in the subsequent sections of this work. In addition, a workflow using only web-based bioinformatic tools was developed to guide experimentalists into capturing a high proportion of published data pertinent to a protein family in a timely and efficient fashion (**Fig. 1**). The DUF34 family, again, allows us to show examples of each of these recommended steps and discuss their strengths and weaknesses in the process of reviewing and characterizing a protein family of unknown function.

**Figure 1.**
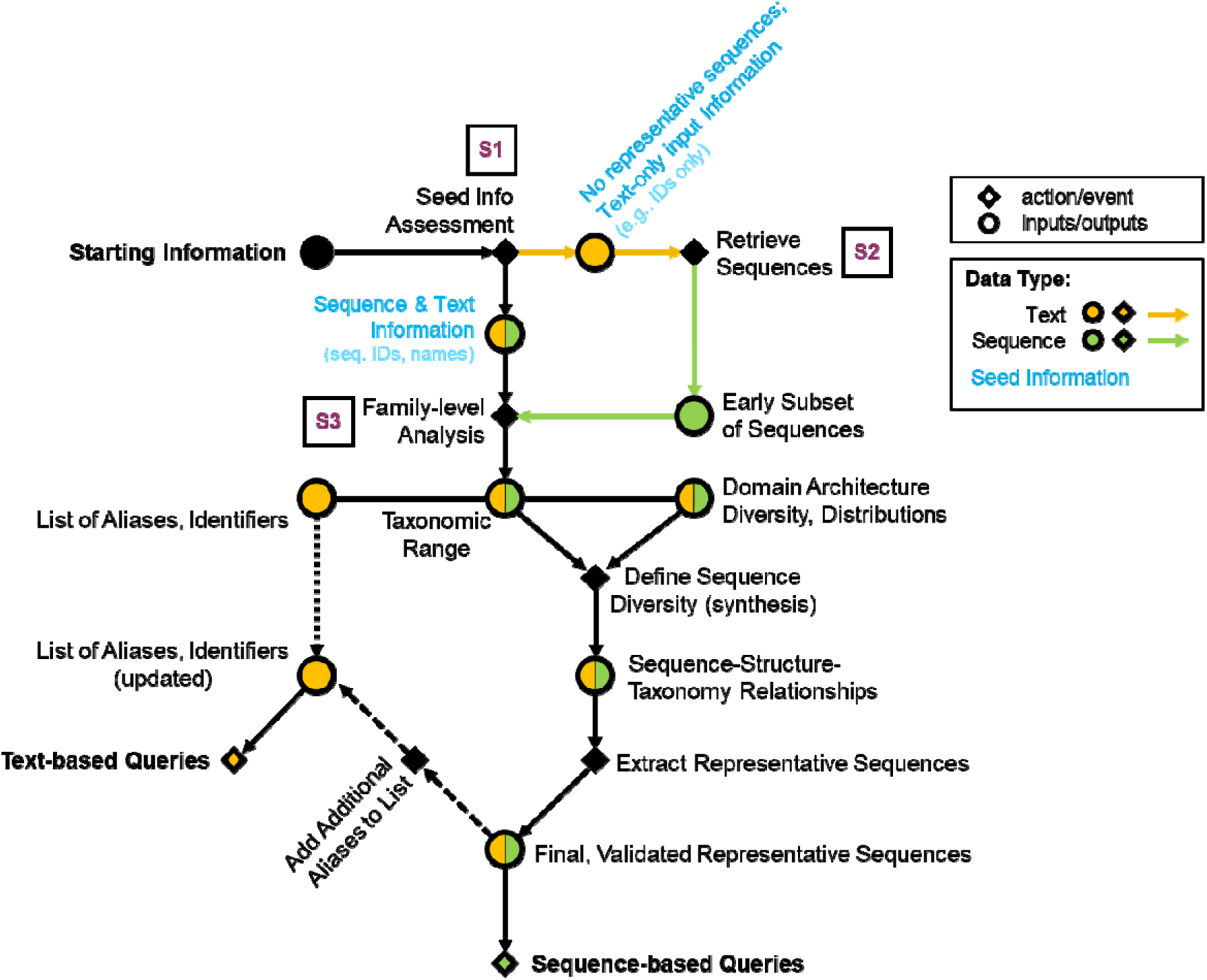
Workflow diagram recommended for capturing published data of a protein family. Supplemental figures were generated for examples (Figures S1-S3). Accompanying supplemental figures are boxed in the diagram and shown in purple text. Nodes of the directed network diagram are distinguished by the shape and color. Circular nodes indicate inputs/outputs (data), while diamond-shaped nodes denote an action or event. The colors of circular nodes convey the type of input/output data: green for sequences and yellow for text. Circular nodes regarding data of both types are split in two (proportions not to scale, do not reflect any guaranteed experience observed with the applied workflow).

### Orthology databases are useful for gathering an overview of protein family domain organization and taxonomic distributions

The first step in any protein family analysis requires the gathering of input data (e.g., a sequence or an identifier) that will be used as seed information for queries (**Fig. 1** and Fig. S1-2) . This process generates two master lists: 1) a list of identifiers, gene/protein names; and 2) a list of representative sequences. Protein family databases such as Pfam [25], InterPro [26], CDD [27], EggNOG [28] are essential tools in generating these two lists.

Orthology databases, broad resources for examining proteins at the family level, are often pre-computed by HMM, bi-directional best hits (BLAST), or motif signatures, and allow for swift analysis of one target family at a time across a predetermined set of genomes (Data S2a). If a seed input sequence is available, it can be used to directly query these databases and extract family names and identifiers, in addition to sequences of other family members. It also provides topical insight into the taxonomic and domain distribution of members of the family, which will guide subsequent queries. For the DUF34 family, this step led to a list of ten most frequently used keywords among UniProtKB entries for this protein family: NGG1 interacting factor 3, NIF3, NIF3L1, GTP Cyclohydrolase 1 type 2, DUF34, YbgI, PF01784, COG0327, YqfO, and COG3323. Without a sequence, known keywords/aliases must be used to acquire sequences from a general protein knowledge database (e.g., UniProtKB [29], NCBI [30], JGI-IMG [31], BV-BRC [32]) (e.g., DUF34 protein family, Data S1) that can then be used to query family databases (Fig. S3b). Together, these processes allow for populating a final list of searchable identifiers/accessions/names (i.e., keywords; **Fig. 1**); this process is completed most comprehensively for the DUF34 family in the context of the later-discussed “QCC (Query, Curate, and Catalog) Cycle” method.

One of the challenges in family-level analyses is the uncertainty of a given family’s domain architectural diversity, as well as their corresponding taxonomic distributions. While orthology databases provide a general view of these attributes, they can be incomplete or misleading as, they erroneously combine or separate families. Examples of this can be seen when navigating the clustered groups and hierarchical relations of the EggNOG (v6) Database. The DUF34 family COG root cluster, LCOG0327, functional annotations include K22391, K07164, and K24730, all of which are incorrectly attributed to this group, the former due to premature EC number assignment in *H. pylori* [24] and the latter two due to DUF34 fusion sequences in bacteria and eukaryotes, respectively (Fig. S4). The aggregation mechanism and presentation manner of annotations by EggNOG implicitly suggests to users that all of these functions are performed by DUF34. However, in truth, these annotations are merely *linked* to DUF34—and aggregated by EggNOG—as a result of DUF34 member fusions with COG1579 (K07164) and CIAO (COG2319, K24730). While this implicit suggestion can be dangerous for propagation, it can also be used to examine the functional associations of DUF34 through comparative genomics, as fusions are often used as a form of guilt-by-association evidence paired with physical clustering data in the prediction of protein function [33,34]. Of note, EggNOG is one of few databases to offer a view feature for family paralog incidence (Fig. S4e), knowledge that can be critical in correctly annotating protein subgroups [35–38].

### Text-based query yields vary quantitatively and qualitatively with both the choice of input and search engine

After establishing and refining the two master lists of keywords and representative sequences (**Fig. 1**, example), text-based queries can be pursued, which will result in the continued accumulation of keywords and representative sequences. The choice of search engine used for text-based queries is ultimately up to the user and examples of the commonly used platforms or “engines” include but are not limited to PubMed, Google Scholar, and Europe PMC. To evaluate how the choices of input keyword and search engine impacted the resulting literature yields and, therein, the information retrieved relevant (or irrelevant) to a target protein family, 10 common keywords associated with the DUF34 protein family were selected and used in iterative searches using 9 popular search engines frequented for the retrieval of scientific literature (i.e., both specialized like PubMed and more broad like Google) (**Table 1**). The results of this systematic survey demonstrated that the total number of text-based hits for this example conserved unknown protein family were highly variable between the distinct combinations of search tool and keyword (**Fig. 2**).

**Figure 2.**
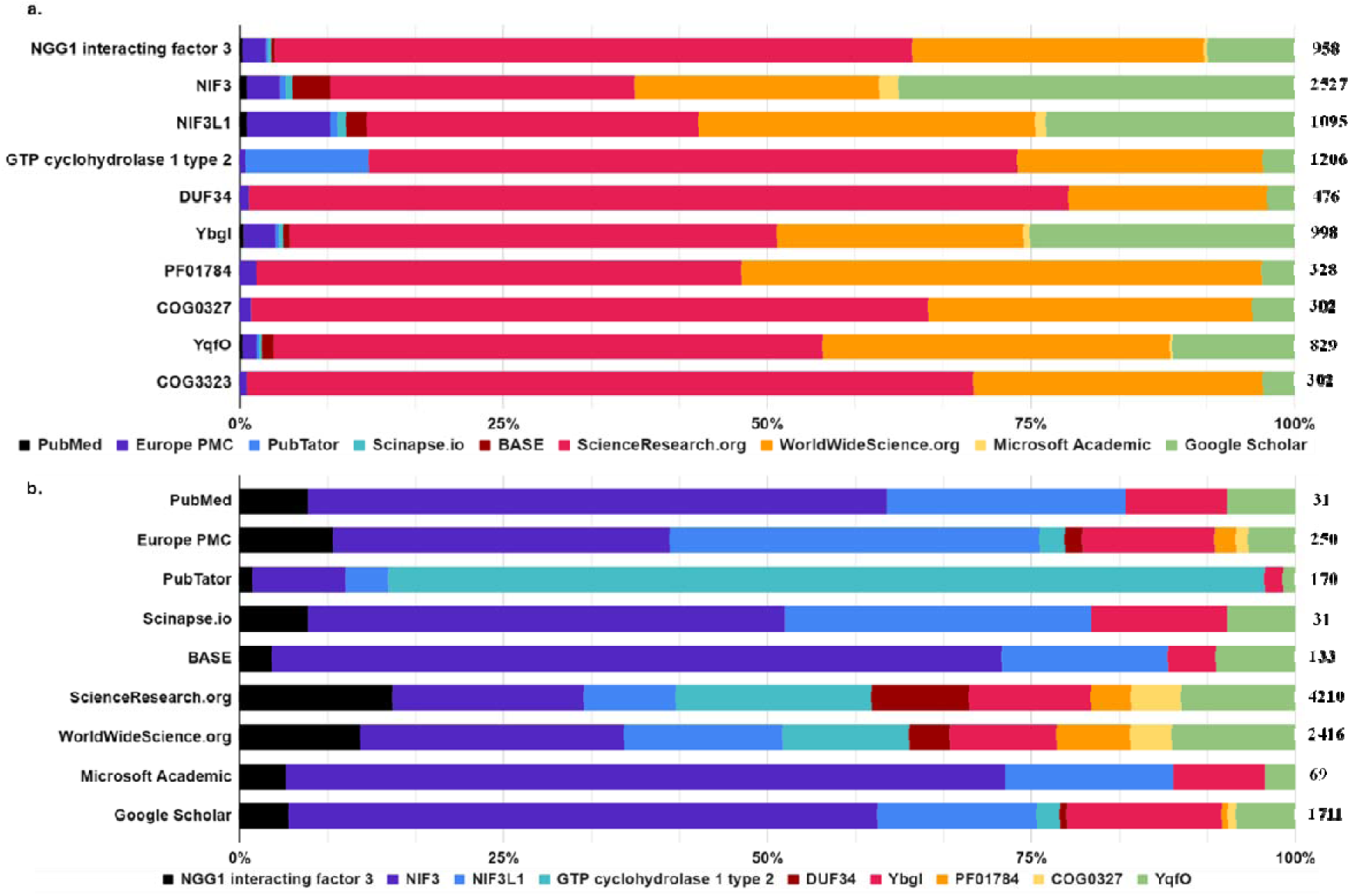
Query yield distributions per search tool as a function of keyword (a) and per keyword as a function of search tool (b). A subset of nine keywords most frequently associated with the target protein family, DUF34, was organized and used to compare the query results (i.e., total hits) across nine distinct search engines commonly used in published data retrieval for scientific research. Totals of each row are shown on the right axis of each figure.

**Table 1.**
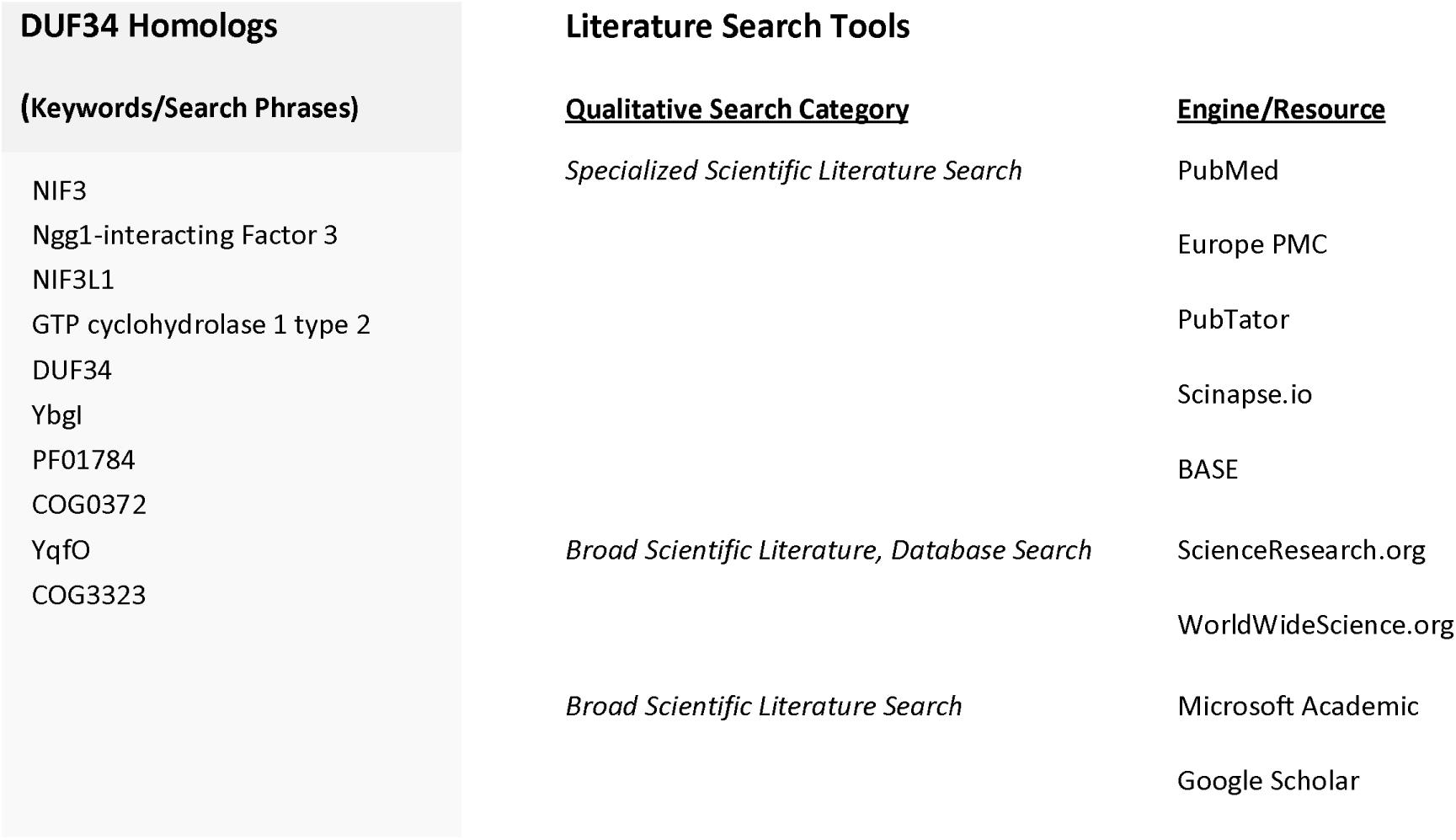
DUF34 homolog search terms and search engines selected for investigating the quantitative impacts of user choice for each upon literature queries.

| DUF34 Homologs | Literature Search Tools |  |
| --- | --- | --- |
| (Keywords/Search Phrases) | Qualitative Search Category | Engine/Resource |
| NIF3<br>Ngg1-interacting Factor 3<br>NIF3L1<br>GTP cyclohydrolase 1 type 2<br>DUF34<br>YbgI<br>PF01784<br>COG0372<br>YqfO<br>COG3323 | Specialized Scientific Literature Search | PubMed |
|  |  | Europe PMC |
|  |  | PubTator |
|  |  | Scinapse.io |
|  |  | BASE |
|  | Broad Scientific Literature, Database Search | ScienceResearch.org |
|  |  | WorldWideScience.org |
|  | Broad Scientific Literature Search | Microsoft Academic |
|  |  | Google Scholar |

To investigate the influence of keyword and search engine choice on the occurrence of false-positive paper yields, a more thorough examination of a subset of specialized search tools was examined (i.e., PubMed, PubTator, Scinapse.io, and Europe PMC) (**Fig. 3**; Data S3). These results indicated a notable presence of false-positive hits among iterations despite the specialized nature of the selected tools. Most of the instances of false-positives were found to be the results of the retrieved publications containing an unrelated scientific term that was identified by the tool as being equivalent to one of the selected keywords (e.g., NiF-3, “Nickel Fluoride 3”) (Fig. S5), the irrelevance of which was only identified through manual curation by the user. In a related observation, Pubtator’s automated query adjustments (a common search engine subroutine intended to improve search result totals by modifying the user-provided keyword, capturing hits for a more generalized version of the term) were found to result in a substantial number of misleading, irrelevant results— specifically for the keyword “GTP cyclohydrolase I type 2”. In these cases, irrelevant results were driven by the expansion of the query to “GTP cyclohydrolase I” without providing the user any notification of the change (**Fig. 3**).

**Figure 3.**
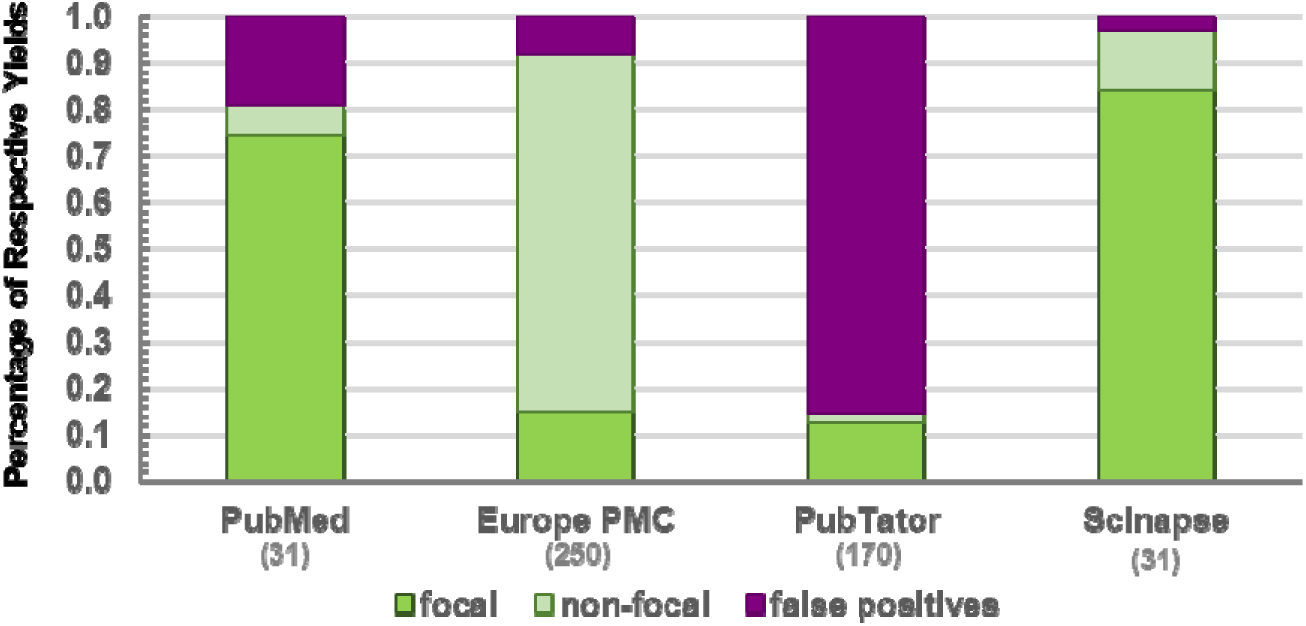
Query yield quality of text-based searches using more research-centered, conservative search tools. Focal publications were labeled as such if they featured relevant keywords specific to the protein family/family homolog in either the title or abstract of the publication. Non-focal publications were labeled if the keywords occurred in any other section of the paper. False positives were manually curated on a case-by-case basis.

Although text-based search tools are widely used by experimentalists, they are vulnerable to false-positives/false-negatives linked to human-computer language “mistranslations” or “miscommunications”. These disconnects between queryable identifiers, and the terms used in publications can be broadly regarded as problems in identifier referenceability. In the case study of DUF34, three major sources of poor referenceability were observed: 1) name/identifier multiplicity (i.e., polyonymy/plurality); 2) mistaken identity (i.e., misleading homonymy/false synonymy); and 3) the ‘published and perished’ phenomenon, discussed further below.

The first source of poor referenceability, name/identifier multiplicity, refers to the problems generated by the many different aliases often assigned to biological entities like protein sequences, which frequently ensure that searches based on any one specific term are always incomplete, as was demonstrated in the DUF34 example (**Fig. 2a**). The second source of poor referenceability can be described as ‘false synonymy’. As gene/protein names are not designed as unique identifiers, the same name can be used for distinct entities by mere coincidence, making it difficult to distinguish, identify, retrieve, or sort them. The DUF34 family alias, ”GTP cyclohydrolase I type 2”, continues to exemplify such problematic homonymy (Fig. S6). In this case, the alias, while widespread in databases across bacterial member sequences, is frequently not recognized by text-based search engine tagging systems/keyword libraries, resulting in a failed primary search that is often automatically generalized by the engine to “GTP cyclohydrolase” in hit retrieval. Although a synonym within reason for the original search term this change invariably retrieves only publications relevant to FolE or RibA, GTP cyclohydrolases I and II. The final source, the so-called ‘published and perished’ phenomenon [39], refers to aliases, descriptions, or characterizations that had been published in the past but have since been overlooked and “re-discovered” by the work of one or more contemporaries, resulting in the independent naming, describing, and/or characterizing of the same entity (Elaboration S1).

### Sequence-based searches stand as a more-than-adequate resource for initial retrieval and review of literature regarding a protein family

As discussed above, text-based searches are defined by their basic functions and reliance upon the design of the engine and user-input preferences, each of which impact the proficiency of natural language-to-computer information processing, searchable data/keyword indexing, and match identification. An alternative to text-based tools are resources that use sequences and family-linked HMMs to search for related publications. To date, several sequenced-based literature search tools have been developed (e.g., Seq2Ref and Pubserver [40,41]) but, at the time of this study, only PaperBLAST [42] remained functional and fully maintained. To determine the publication retrieval productivity differences between these two search tool types, the results for a PaperBLAST query of a single DUF34 homolog, YbgI of *E. coli* (UniProt: P0AFP6), were compared to those of PubMed text-based searches for the same protein. For use in querying PubMed, 9 keywords (i.e., “YbgI”, “P0AFP6”, “b0710”, “JW0700”, “PF01784”, “NIF3”, “Ngg1-interacting factor 3”, “DUF34” and “COG0327”) were collected from the UniProt entry page for this homolog (UniProt: P0AFP6). Using default search settings, a total of 47 unique publications were retrieved with PaperBLAST, of which only 3 were determined to be false positives (6.4%; **Fig. 4** ; Data S4). In contrast, while PubMed returned a total of 19 unique publications, 6 of these were found to be false positives (32%). Although PubMed’s release announcements suggest nigh ubiquity of full-text search across their database and increasingly more advanced search features (https://www.ncbi.nlm.nih.gov/feed/rss.cgi?ChanKey=PubMedNews), the comparative results between these two tools suggest that PubMed is not generally able to recognize search terms in a publication if they are located outside of the title and/or abstract. This is made evident—at least in the case of the keywords associated with this particular DUF34 homolog—by the recognition of “non-focal” publications by PaperBLAST and the complete oversight of the same publications by PubMed (**Fig. 4**), even though these 34 publications were all present in the full-text depository of NCBI (PubMed Central, PMC) of which PubMed is designed to query (Data S4d).

**Figure 4.**
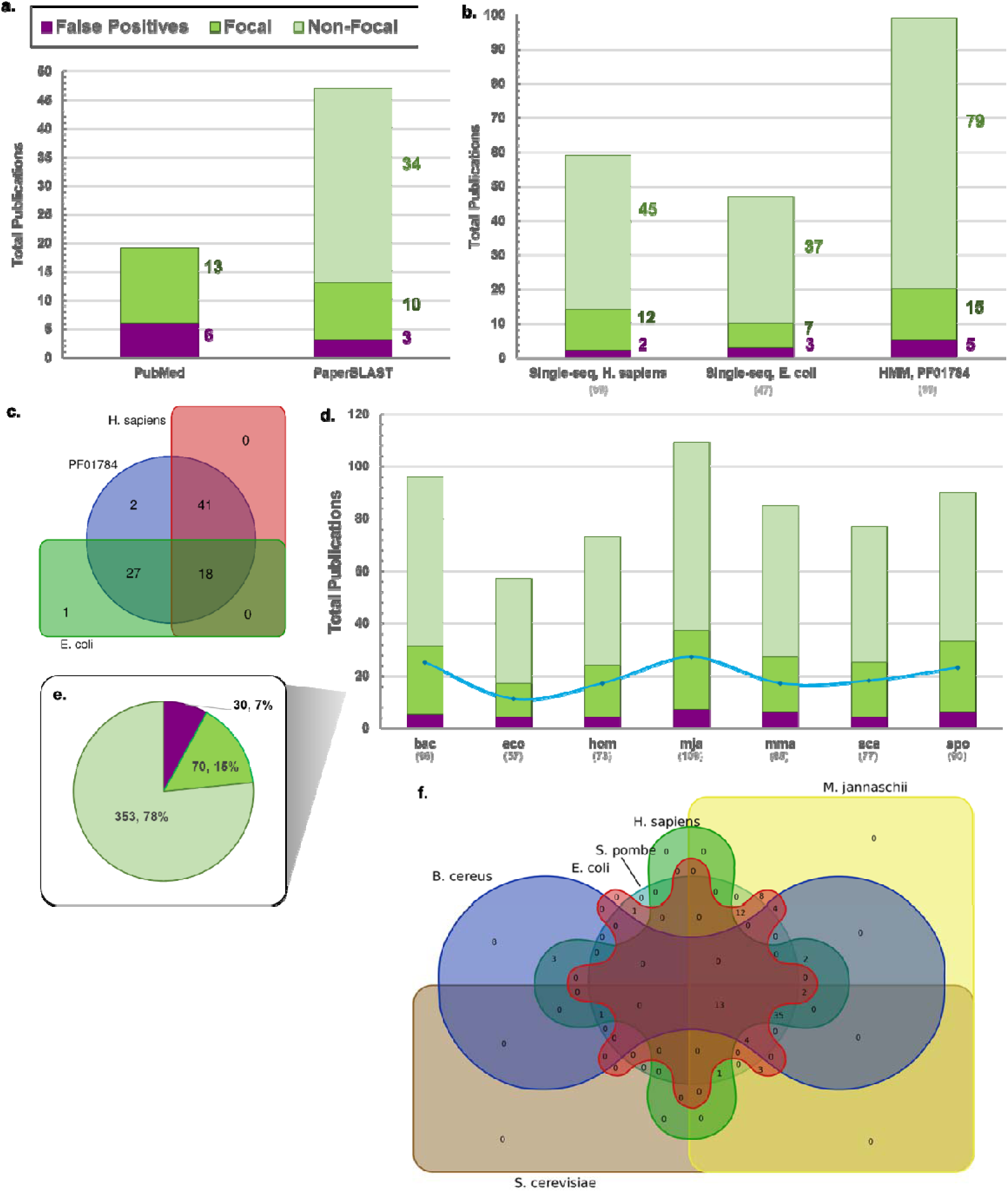
Quantitative and qualitative analyses of sequence-based literature search using PaperBLAST. (a) The yields of a PubMed text-based searches of 9 UniProt entry-derived keywords (source UniProt entry: E. coli DUF34 homolog, P0AFP6; keywords: “YbgI”, “P0AFP6”, “b0710”, “JW0700”, “PF01784”, “NIF3”, “Ngg1-interacting factor 3”, “DUF34”, “COG0327”) compared to those of PaperBLAST using one single-sequence-based query of the same source UniProt entry sequence. Bar color key for “false positives” (purple), “focal” (green), and “non-focal” (light green) designated publications applies to this panel (a) and panels (b), (d), and (e). “Focal” publications were defined as those having mentions of respective homologs in the abstract and/or the title, while “non-focal” were those with homolog mentions anywhere else in the publication and/or supplemental materials. (b) Comparison of publication retrieval methods available through PaperBLAST (i.e., HMM or Pfam identifier-input-based query compared to single-sequence-based queries). Two disparately related DUF34 family homolog sequences (one used in the preceding comparison of text-and sequence-based queries (a) and the other of a model organism, H. sapiens) were chosen for comparison to the family’s linked HMM profile identifier extracted from Pfam (PF01784). (c) Visual comparison (Venn diagram) of unique yields derived from the prior sequence- and HMM-based literature searches using PaperBLAST. The results for the homolog of E. coli (UniProt: P0AFP6) are shown in green area and those for the H. sapiens homolog (UniProt: Q9GZT8) are shown in red. The results of the HMM-based query using the family Pfam (PF01784) are shown in blue. (d) Query quality of PaperBLAST hits per DUF34 protein family member sequence, one query sequence per organism. Organisms selected by tentative domain architectural subclasses and taxonomic distribution of members. A blue line marks the occurrence of redundant results within a single query (avg. ∼23% of hits per query). All selected query sequences have been independently described in a scientific publication [24]. Total hits per sequence/query are shown below the x-axis labels. (e) Overall query quality across all representative sequences used to query PaperBLAST shown in (d). Total of represented hits equals 453. (f) Visual comparison (Venn diagram) of all methods of literature retrieval examined here. Yields shown are 6 of the 7 sets of single-sequence PaperBLAST results; diagram generating tool could not suit the total of 7 lists. With this, the results for the M. maripaludis DUF34 homolog were not observed to have provided any novel results not retrieved by the other 6 sequences and so its yield list was not included in the generation of this figure.

To assess the different uses of PaperBLAST for the retrieval of publications relevant to many members of a protein family, two separate search options were compared: via HMM or via single sequence/sequence ID. These two methods were used to generate itemized lists of publications for two family member sequences (*H. sapiens*, UniProt: Q9GZT8; *E. coli*, UniProt: P0AFP6; default settings) and for one HMM classification identifier representing the same family (Pfam: PF01784). False positives were still present among the results for both PaperBLAST retrieval methods (**Fig. 4** ; Data S5-S6). Despite querying the same database, the two single-sequence queries differed in their results from, both, one another and compared to the HMM-based search. *H. sapiens* and *E. coli* DUF34 homolog sequence results were found to share 9 and 7 unique publications with the HMM-based results, respectively (**Fig. 4**; Data S5-S6). Only the retrieved set for the *E. coli* family member sequence retrieved a publication (1) unique from both the *H. sapiens* homolog results and those of the HMM-based search, suggesting two possibilities: 1) that the curation status of the latter two have the greatest similarity in quality, which appears higher than that for the bacterial family member sequence; and/or 2) that publications of the eukaryotic model system of *H. sapiens* contribute the bulk of the overall family related literature accessioned by PaperBLAST’s database.

Because of the differences observed in PaperBLAST outputs when using the *E. coli* or *H. sapiens* outputs, we repeated PaperBLAST queries using seven diverse sequences from the DUF34 family reflecting different superkingdoms and alternative domain architectures (**Fig. 4**; Data S7-S9). The differences in output first observed with two sequences were confirmed with more sequences tested. One set of results, those of the *B. cereus* DUF34 homolog (UniProt: Q818H0), produced publications that were entirely unique among the seven queries, unshared with the results of other single-sequence queries. Coincidentally, the homolog of *B. cereus* contains an inserted domain distinguishing it and others like it as members of a putative functional subgroup of the DUF34 family [24]. Therefore, these unique retrieved publications reinforce that an understanding of the taxonomic distribution of protein family domain architecture diversities is important to develop prior to selection of representatives for single-sequence-based literature retrieval via PaperBLAST.

In summary, the general lack of standards and guidelines across the community, of which could be implemented by publishers [43], make extracting published information on all members of a protein family a time-consuming and error-prone exercise, particularly when relying upon text-based search tools. Sequence-based searches should be the starting point of any family level review, as demonstrated above. This can be complemented by text-based searches, but any outputs derived from these queries should also be checked by sequence for the expected protein family membership.

### Comprehensive literature search: capture through iterations of queries in parallel with the accumulation of keywords and representative sequences

Ideally, a comprehensive capture of all publications linked to all members of a protein family would require: 1) both text-based and sequence-based search tools; and 2) iterative cycles of querying, curating, and cataloging (“QCC cycle”, **Fig. 5**). However, there exists an inevitable law of diminishing returns with such a process (e.g., fewer total new relevant publications per hour), with productivity gradually decreasing as more time passes until no new publications are retrieved with continued iterations. Even for a dedicated biocuration expert, the total amount of non-redundant data retrieved per unit time exponentially decreases after a certain number of hours. The amount of time necessary to invest, however, will differ between protein families and the time investment determined appropriate will vary between researchers performing the searches.

**Figure 5.**
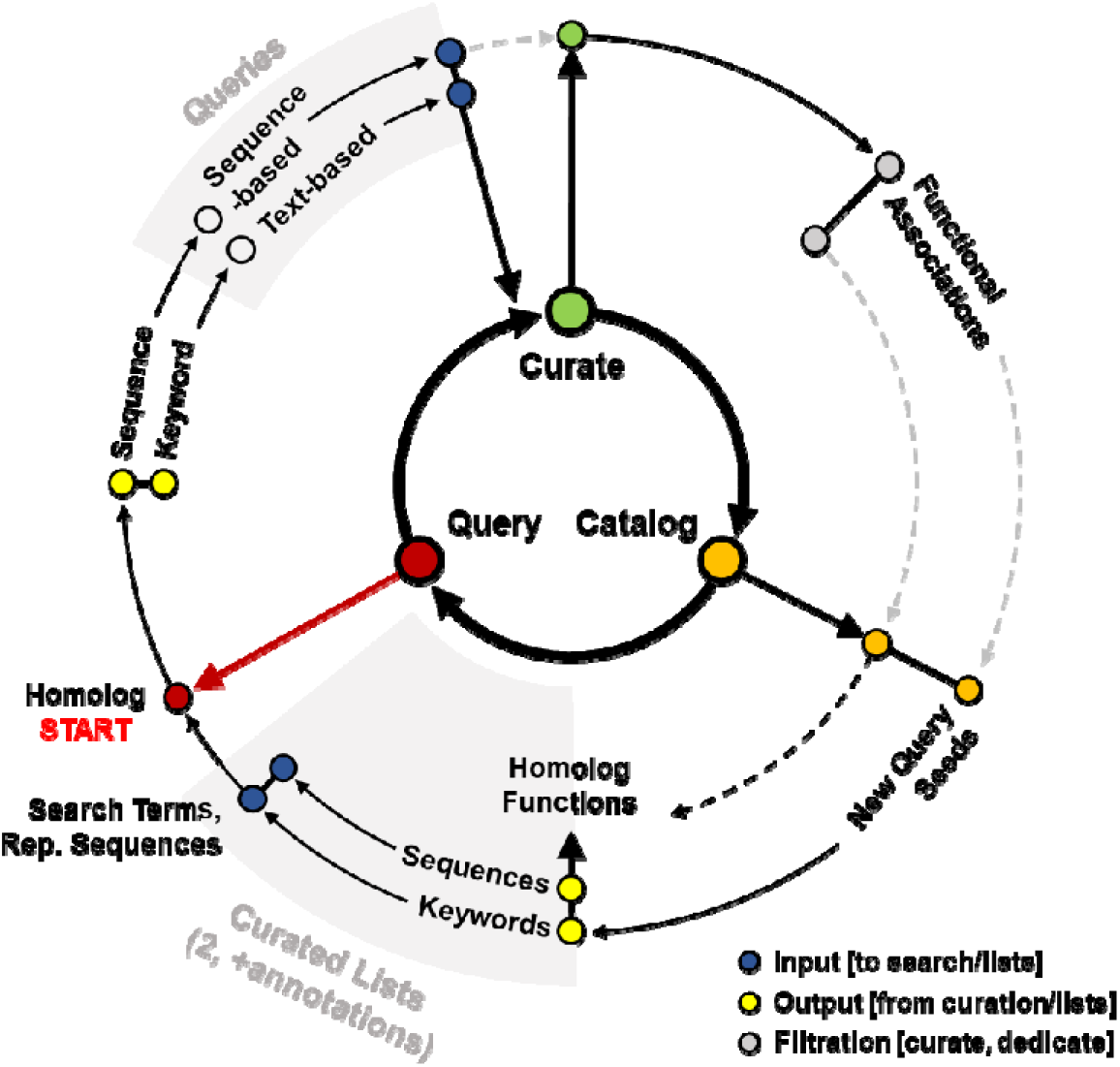
Idealized cycle of accumulating keywords, homolog annotations representative sequences, and publications necessary to optimize the capture of all published data relating to a protein family. Query phase refers to the process (often multiple in parallel) of using various webservers to retrieve literature relevant to target homolog(s). The Curate phase is distinguished by its filtration and review of retrieved information, and, frequently, the identification of experimentally validated functional associations to be noted for select homologs in the subsequent phase. The final phase of the cycle, the Catalog phase, is defined by the multiple diverging paths along which the different identified information will be channeled. These distinct paths of information included the two curated lists of keywords and representative sequences, as well as a collection of experimentally validated functional annotations of select homologs (publications cited). Multiple nodes within a single radial location indicates a split or merge, depending on the direction of the respectively linked arrow. Light gray dashed arrows indicate implicit information flow, whereas the black, solid arrows indicate explicit information flow. The dashed, black arrow denotes explicit information flow out* of the QCC cycle.

To optimize search time, an additional investigation was undertaken to better understand the differences between the sequence- and HMM-based methods of PaperBLAST and the high-investment approach presented by the “QCC cycle”. The results of four different query result sets henceforth referred to as four distinct “methods” of retrieval ({1} HMM-derived, {2} PubMed text-derived, {3} QCC/Curated-derived, and {4} three separate sequence-derived sets: (A) one using only *E. coli* DUF34 homolog sequence; (B) a second that was a merged pair derived from two sequences (DUF34 homologs of *E. coli*, *H. sapiens*); and (C) a third constituted by merging seven sequence-derived query result sets (DUF34 homologs of *E. coli*, *H. sapiens*, *B. cereus*, *M. jannaschii*, *M. maripaludis*, *S. cerevisiae*, *S. pombe*, sequences chosen to represent the putative diversity of the family) were compared for determining overlaps and method-unique yields (**Fig. 6a**; Table S2; Data S9). Only 5 unique publications were shared between all three PaperBLAST-derived result sets (methods {1}, {3} and {4, merged}). There were no results unique to any one of the single-sequence-derived results (method {4}, lists A-C). A comparison of all different types of literature search methods focusing on their relevant results (i.e., those classified as “non-focal” or “focal”) demonstrated that no single method, text- or sequence-based—regardless of the number of sequences—can capture everything available, emphasizing the needs of these distinct methods of search (**Fig. 6b**). Moreover, the sequence-based searches retrieved seven publications that were not captured by the HMM-based results, the two single-sequence-based results (method {4}B), the one single-sequence-based result (method {4}A), or the idealized “QCC cycle” method (**Fig. 6**; Data S10). These data suggest that the HMM-based method was the most efficient approach of the two offered by the PaperBLAST suite when seeking a more comprehensive family-level review of the corpus in the least amount of time and use of only one resource. Finally, when comparing PaperBLAST as a resource overall to the far more tedious “QCC cycle” approach, it was observed that PaperBLAST still misses ∼75% publications relevant to the example DUF34 family, so one cannot avoid the iterative and semi-manual “QCC cycle” approach, for now, to capture the entirety of published knowledge across the corpus for any given protein family.

**Figure 6.**
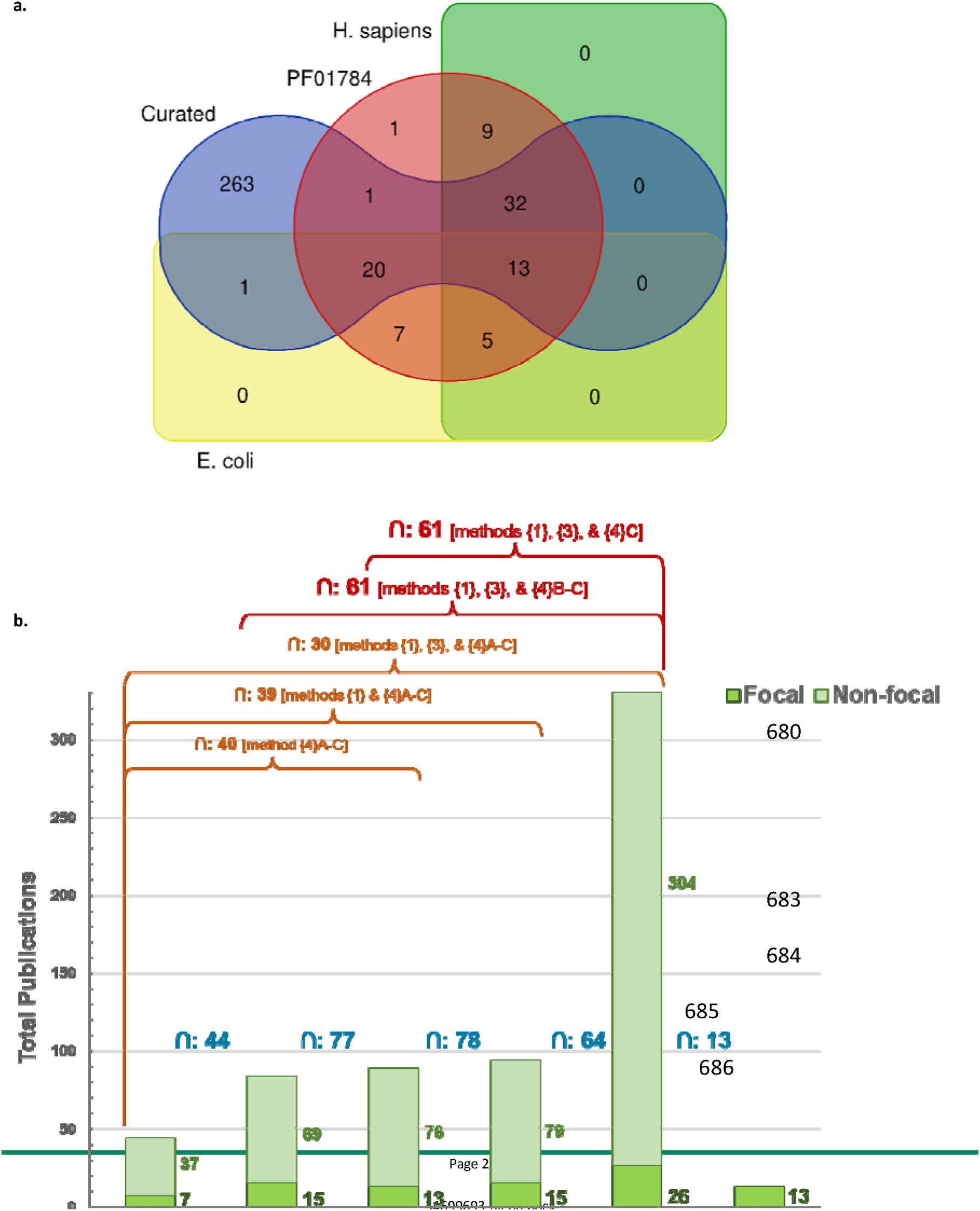
(a) Comparison of literature yields for selected PaperBLAST retrievals and the QCC cycle method. Asymmetric Venn diagram illustrates the distinct hits for each method. “H. sapiens Seq” (green area) denotes the unique results of single-sequence PaperBLAST output for H sapiens DUF34 homolog sequence (UniProt: Q9GZT8). “E. coli Seq” (yellow area) denotes the unique results of single-sequence PaperBLAST output for E. coli DUF34 homolog sequence (UniProt: P0AFP6). “Curated” (blue area) denotes the unique results of the idealized “QCC” Cycle output for the comprehensive investigation of the DUF34 family. “PF01784” (red area) denotes the unique results of the HMM-based PaperBLAST query output (HMM/Pfam: PF01784). (b) Focal and non-focal publications captured across all methods. Stacked bar plot of unique focal and non-focal results of all distinct literature retrieval methods. Intersection of yields between each pair of methods is shown in blue. Orange labels denote the “method” group, as do the red labels, the color of which were changed from those of the orange variety to emphasize the data patterns described in the text.

The taxonomic distribution of protein families and their respective subgroups have been demonstrated to impact the literature capture process through the availability heuristic [44,45] and unique biases influencing researcher keywords and search engines of choice. That is, any given researcher may have a different starting idea of a protein family and this beginning governs their ultimate search results when attempting to broaden their understanding of the same protein family. However, there are many methods one may use in microbiology to better understand proteins at the family level before completing a literature search. In addition to the strategies addressed above, the improvement of your starting conceptions of a family’s diversity and putative subgroups can be accomplished using a variety of phylogenomic and phylogenetic tools. These tools are discussed in detail in the subsequent sections.

### Resources that examine the taxonomic distributions of protein families are essential components of comparative bioinformatic analyses

Before comparing the tools important for family-level analyses, it is important to, first, define some of the key terms used to describe them. Family-level bioinformatic tools are primarily split into two categories: 1) phylogenomic and 2) phylogenetic. “Phylogenomic” describes analyses considering whole genomes or large regions of genomes. Tools investigating gene synteny and neighborhoods are often considered “phylogenomic”, depending on the nature of their genomic comparisons. Phylogenetic tools focus on the presence of individual gene/protein sequences compared between genomes. For years, PubSEED [46] had been an ideal resource for examining taxonomic distributions of protein families, as a user could rename member proteins and visualize their distributions in sets of genomes or user-defined “Subsystems” with a color-coding system that highlighted synteny and that could be used to gather both phylogenomic and phylogenetic data. If PubSEED is still functional it is frozen at ∼10,000 genomes and so cannot be considered a main source for analyzing taxonomic distributions when over 200,000 complete genomes are now available. Here, we surveyed different types of webserver-based resources designed for phylogenomic and phylogenetic analyses (Data S2a). These resources can be separated into several types: A) general orthology databases (precomputed phylogenetic distributions; can also include protein family classification databases); B) synteny/gene neighborhood databases (precomputed phylogenomic record data, distributions; often features within larger databases); and C) phylogenetic pattern/profile databases (often features within larger databases; precomputed phylogenetic distributions) (**Fig. 7**). These three types of analyses can be further defined by the respective parametric restrictions of the tools used to execute them into four subtypes: 1) custom genome selection with single target family selection; 2) tool-defined genome selection with single target family selection; 3) tool-defined genome selection with multiple target family selection; and the rarest of the subtypes, 4) custom genome selection with multiple target family selection. An additional feature among these tools is sequence-based tool input or sequence similarity-based thresholds in family identification across genomes (also noted in **Fig. 7** as a minor subcategory of “Target Selection”). These four phylogenetic/phylogenomic tool subtypes are detailed further below, and a subset we use most frequently for both teaching and research is given in **Table 2** . The strengths and weaknesses of these tools are compared and discussed below using the specific DUF34 family-linked clustered orthologous groups (COGs) identified previously: COG0327, COG1579, and COG3323 [24].

**Figure 7.**
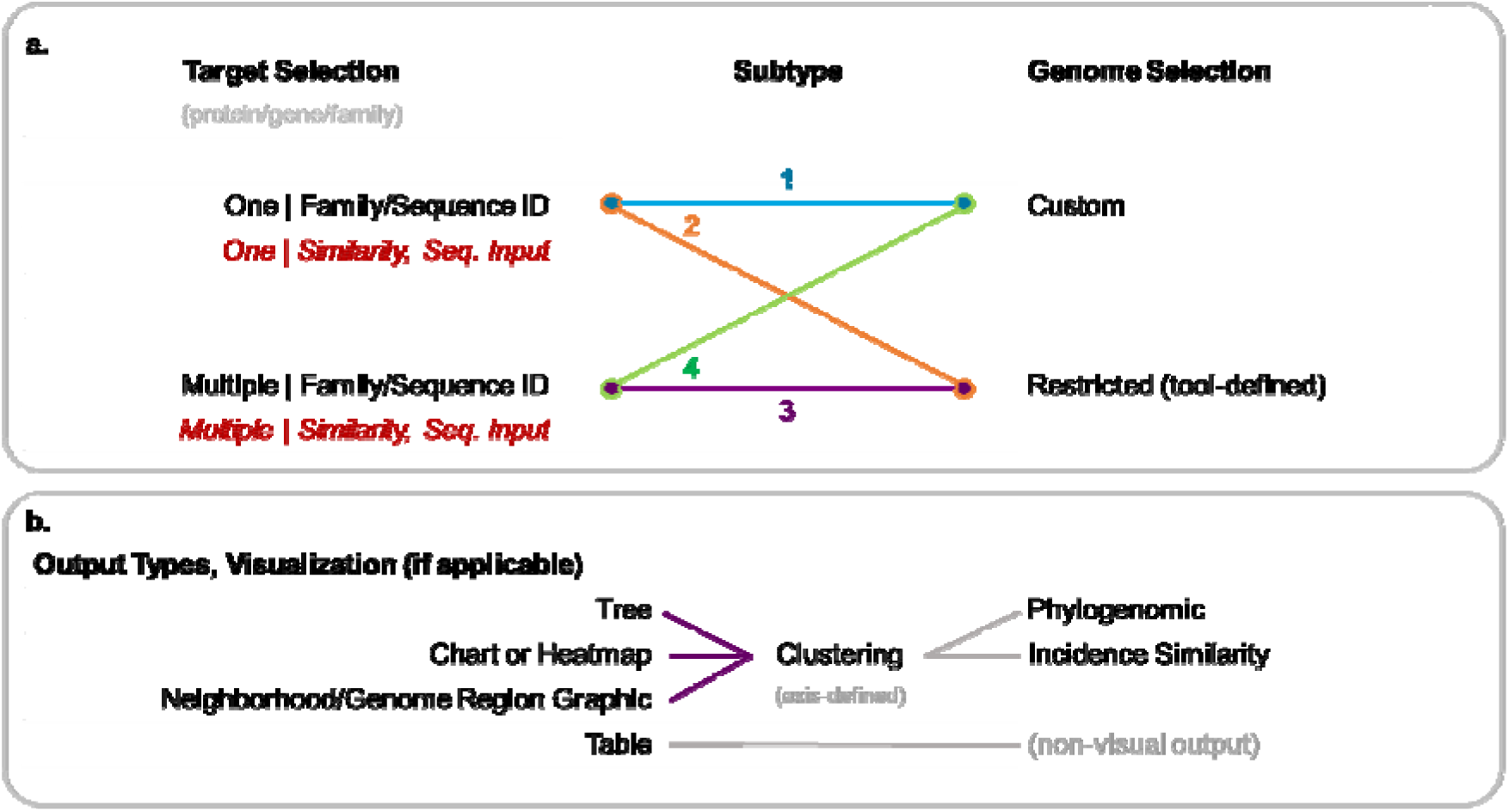
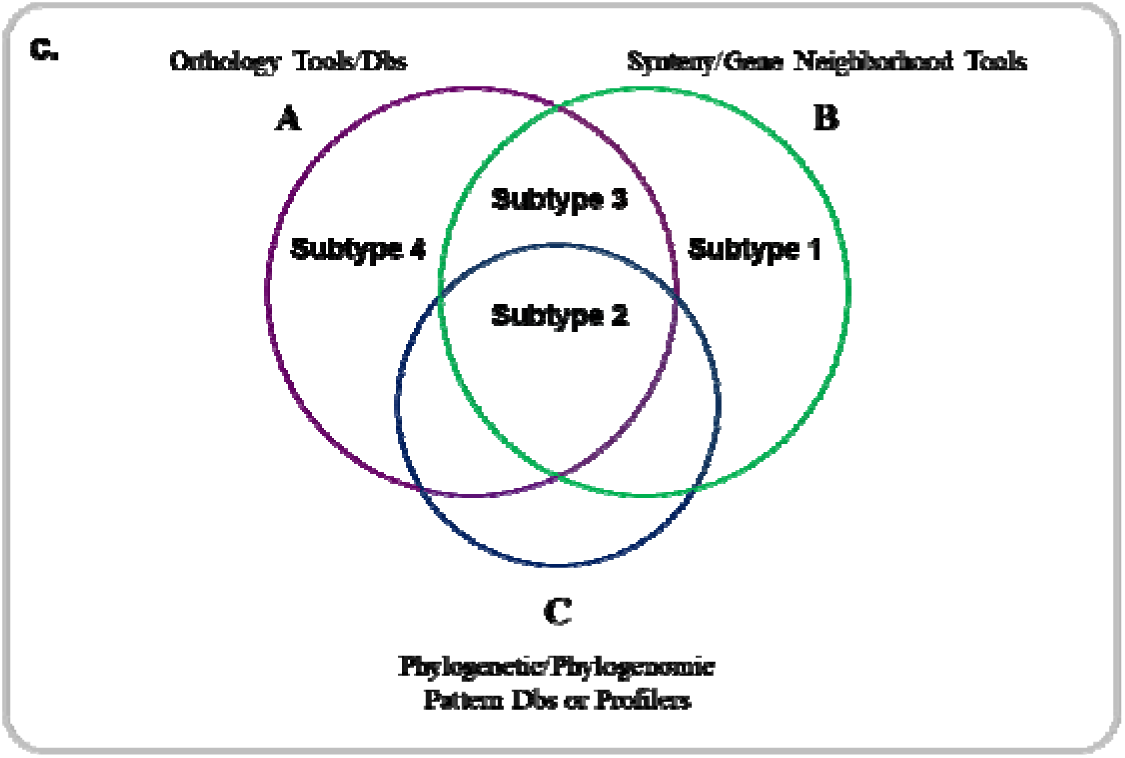
Categorization of phylogenomic, phylogenetic analyses and the diversity of tools available for performing them. (a) A list of three major categories of phylogenetic/phylogenomic analyses are listed first, A-C. (b) These major categories of analyses are further broken down into subtypes of tools used to perform those analyses. (c) All tools have a variety of outputs and result visualizations, and are, below the prior, examined in the context of the root output type, as well as any kernel-based clustering and corresponding type of clustering relative to axis/data-type being clustered (if applicable). (d) A summary diagram that illustrates the effective overlap of tool subtypes relative to the three major analysis types.

**Table 2.**
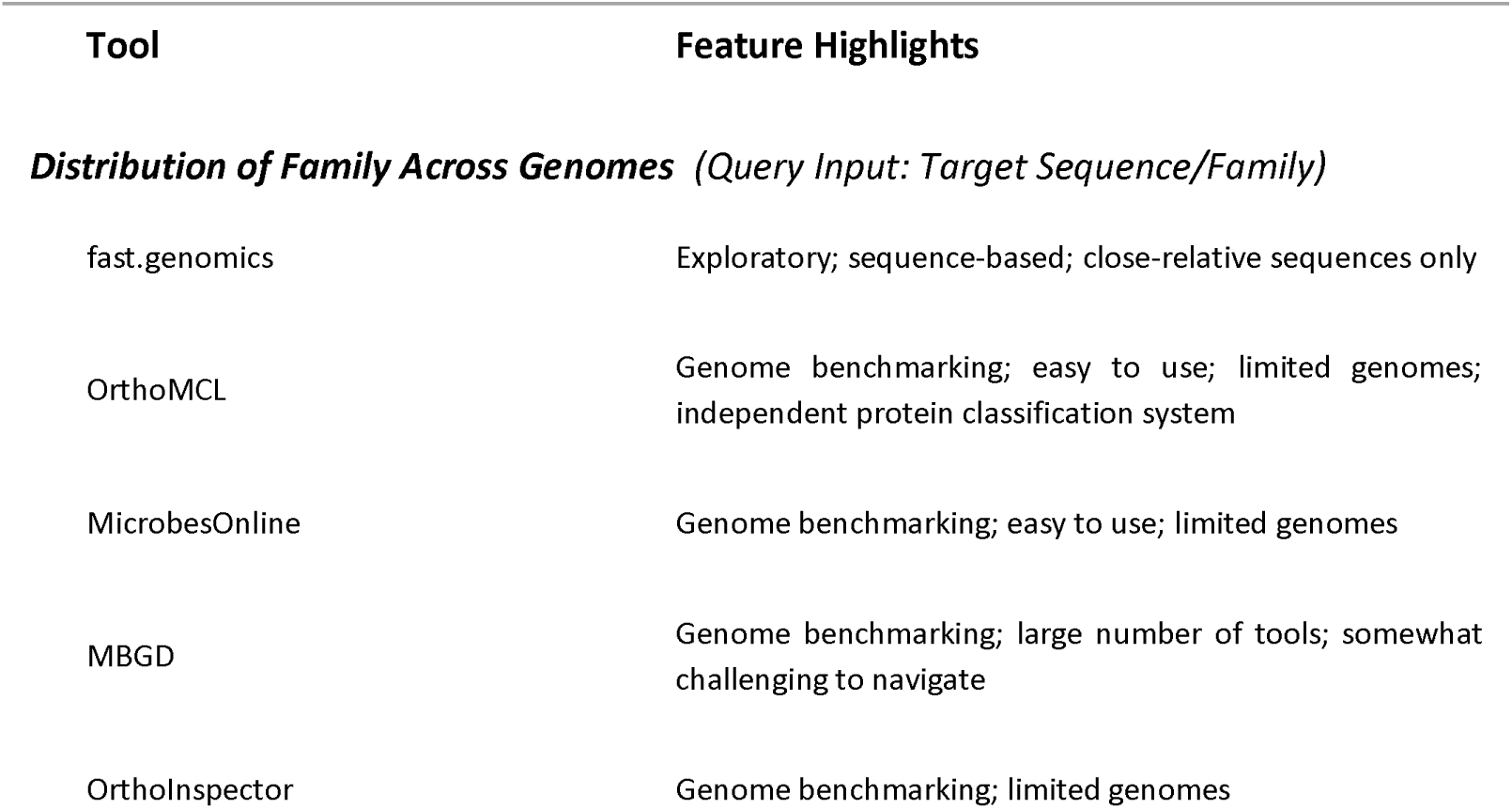
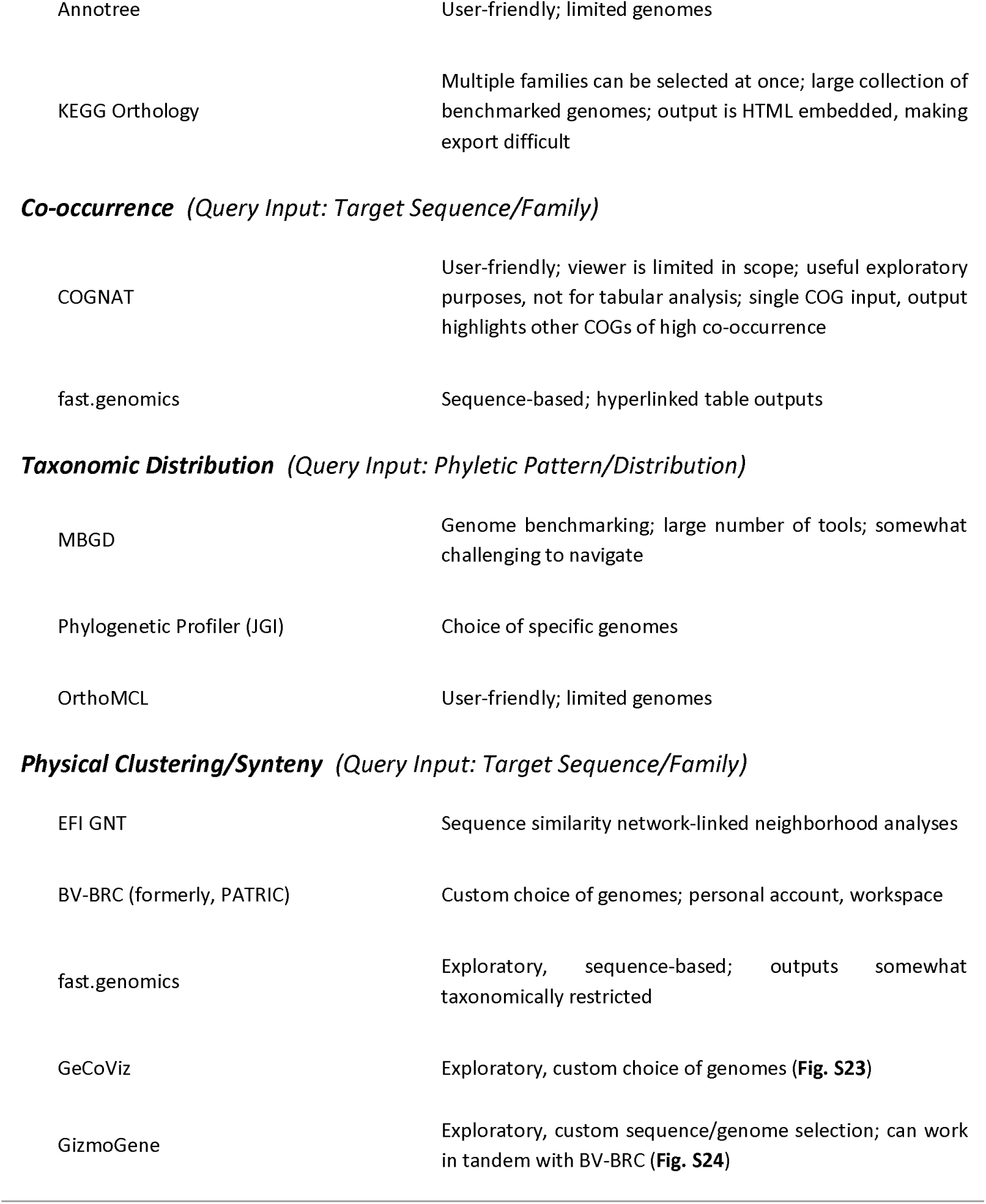
Preferred tools by phylogenomic/phylogenetic objective.

| Tool | Feature Highlights |
| --- | --- |
| *Distribution of Family Across Genomes (Query Input: Target Sequence/Family)* |  |
| fast.genomics | Exploratory; sequence-based; close-relative sequences only |
| OrthoMCL | Genome benchmarking; easy to use; limited genomes; independent protein classification system |
| MicrobesOnline | Genome benchmarking; easy to use; limited genomes |
| MBGD | Genome benchmarking; large number of tools; somewhat challenging to navigate |
| OrthoInspector | Genome benchmarking; limited genomes |
| Annotree | User-friendly; limited genomes |
| KEGG Orthology | Multiple families can be selected at once; large collection of benchmarked genomes; output is HTML embedded, making export difficult |
| **Co-occurrence** (*Query Input: Target Sequence/Family*) |  |
| COGNAT | User-friendly; viewer is limited in scope; useful exploratory purposes, not for tabular analysis; single COG input, output highlights other COGs of high co-occurrence |
| fast.genomics | Sequence-based; hyperlinked table outputs |
| **Taxonomic Distribution** (*Query Input: Phyletic Pattern/Distribution*) |  |
| MBGD | Genome benchmarking; large number of tools; somewhat challenging to navigate |
| Phylogenetic Profiler (JGI) | Choice of specific genomes |
| OrthoMCL | User-friendly; limited genomes |
| **Physical Clustering/Syteny** (*Query Input: Target Sequence/Family*) |  |
| EFI GNT | Sequence similarity network-linked neighborhood analyses |
| BV-BRC (formerly, PATRIC) | Custom choice of genomes; personal account, workspace |
| fast.genomics | Exploratory, sequence-based; outputs somewhat taxonomically restricted |
| GeCoViz | Exploratory, custom choice of genomes (**Fig. S23**) |
| GizmoGene | Exploratory, custom sequence/genome selection; can work in tandem with BV-BRC (**Fig. S24**) |

#### Subtype 1: custom genome selection, single target family selection

Phylogenetic and phylogenomic analyses, in general, are largely governed by two variables, as alluded to in **Fig. 7** : 1) the number of targets (i.e., families, groups, neighborhoods) viewable/analyzed at once, and 2) the number of genomes one can view these data across at once (and whether those genomes can be custom selected). Many of the tools available via webserver are restricted by one or the other, often both. Additionally, it is common for a webserver’s workflow to begin with a single sequence or family identifier, a paradigm common across phyletic tool subtypes defined in this work. Divergences from this framework are discussed in later subsections (e.g., see discussions of BV-BRC), but are also more commonplace in the first of our four defined subtypes. One of the subtype 1 example resources for phylogenetic analyses, JGI-IMG, can begin with a single sequence/family but can also begin at the level of user-selected genomes, taxonomic ranges. Most subtype 1 tools are components or features within a larger suite or database, with or without an account-linked workspace. Physical clustering is a key type of association-based inference derived from genomic sequences and links genes to putative functions based on the annotations of their encoded neighbors, given that strong conservation is observed [33]. Many gene neighborhood or physically clustered gene viewers also fall within this subtype (**Fig. 7**). One example of a free-standing subtype 1 gene neighborhood viewer is WebFlaGs [47] (https://server.atkinson-lab.com/webflags) (Fig. S7), which permits the user to input many sequence identifiers (i.e., NCBI protein accessions) within a protein family for the generation of a taxonomically clustered set of gene neighborhoods.

Sequence similarity networks (SSN), while not classified as a major analysis type to be shown in **Fig. 7**, are becoming more common across comparative genomics [48,49]. A popular SSN-generation tool, EFI’s Enzyme Similarity Tool (EST) [8,50] (https://efi.igb.illinois.edu/efi-est/) (Fig. S8), falls within the subtype 1 group, specifically for its primary means of job submission (“Sequence BLAST” and “Families”). Additionally, this suite provides options for gene neighborhood analyses, either linked to submitted EST jobs or independently generated user-submitted SSNs. Because of the numerous options made available within the EFI suite, this tool could also be considered subtype 3 or even subtype 4, depending upon a user’s creativity in implementing the sequence list-based job submission form and other features.

#### Subtype 2: tool-defined genome selection, single target family selection

Subtype 2 phylogenomic/phylogenetic tools are by far the most common and are often embedded as part of a larger database (e.g., CDD, PANTHER, MBGD). Because they usually require little computation or are precomputed, they are ideal for a first pass investigation of a protein family.

A user-friendly subtype 2 tool for deriving family-level distribution information is the phylogenetic tree viewer, Annotree [51] (http://annotree.uwaterloo.ca/annotree/). Building on the protein family information derived from Pfam (now integrated into InterPro), TIGRFAM (no longer maintained), or KEGG (KO families), Annotree provides a practical “first pass” in examining a protein family’s taxonomic distribution [51] (Fig. S9). Several output parameters can be actively modified by the user in-browser (e.g., taxonomic ranges for tree branching and, separately, labeling of those ranges). However, Annotree is restricted to bacterial and archaeal taxa and does not allow for the examination of multiple target families.

Gene neighborhood and synteny tools that fall within subtype 2 are usually free-standing while also being dependent upon the aggregation of information from other databases (e.g., COGNAT [52] (https://depo.msu.ru/module/cognat) (Fig. S10)). An embedded feature of the KEGG Database makes it possible to extract “Gene Cluster” (gene neighborhood) information for individual genes, sometimes provided in parallel to neighborhood data of closely related organisms (that is, if the majority of the Cluster is also conserved across those closely related target homologs). However, this tool does not provide an option of viewing these data across taxa as a distribution per target gene/protein family. Even more, this viewer is without an option for tabular export. Another subtype 2 comparative gene neighborhood tool that is also a component within another database is the Genomic Neighborhood Comparison viewer subsection each *B. subtilis* gene entry within the recent beta release (“December update”) of SubtiWiki’s “CoreWiki” (http://corewiki.uni-goettingen.de/welcome) [53] (Fig. S11). This feature provides a swift overview of the homologous gene neighborhoods of select model bacterial alongside that of the entry page’s target gene and corresponding neighborhood. The additional genomes featured in the viewer are fixed by the database and the visualization is generated within-page using the beforementioned subtype 1 tool, WebFlaGs.

#### Subtype 3: tool-defined genome selection, multiple target family selection

Phylogenetic/phylogenomic tools that are classified here as “subtype 3” allow for the selection of multiple target gene/protein families but are restricted in the genomes across which those families may be viewed or analyzed. Examples of precomputed phylogenetic databases of this subtype include MicrobesOnline, STRING-DB, *fast.genomics*, and KEGG Orthology (KO). MicrobesOnline, while being a multi-faceted sequence database, allows the user to choose a set of input families using different types of systematic identifiers like COGs or Enzyme Commission (EC) numbers for generating phylogenetic profiles, which are graphically produced and clustered taxonomically with the absence-presence of the families/members across the database’s benchmark 1,965 organisms (Fig. S12). This tool is notably user-friendly with different methods of family member identification/filtration possible for selection per target; in addition to systematic identifier annotations, options for these filters also include several BLAST cut-offs (Fig. S13). Users may also view the precomputed phyletic profile for a single family via any gene entry’s “Gene Info” tab (Fig. S14). Unfortunately, MicrobesOnline is, to-date, frozen at a total of 3,707 genomes (retrieved January 14^th^, 2022), and, further, these genomes are largely limited to bacterial organisms with only 94 archaea and 119 eukaryotes, the latter of which are mostly fungi.

While primarily an annotation network visualization tool, the STRING Database also features a tool designed for the rapid survey of phylogenetic co-distribution of protein families (Fig. S15). While useful for exploring annotations and hypothesis generation, the tool is a poor source of primary data without the paired implementation of much more systematic, stringent analytic pipelines.

Because the KEGG database uses relatively stringent family relationships to create their orthologous groups (i.e., “KOs” or K numbers [54]), we find that the KO database can be quite useful to analyze the phylogenetic distribution of specific families. The tool produces a table of protein distribution among all genomes present in the KEGG dataset using KO identifiers (Fig. S16a), which the user provides in a space-separated list (Fig. S16b). However, not all protein families have been assigned to a KO group and the genomes are organized in an order without clear reference to taxonomic relations and the data shown is generated based upon the genomes in which at least one of the submitted KOs occurs. Because of the latter feature, it is recommended that, with tools like this, the user co-submit a positive control KO (i.e., a group that is known to be universally conserved across database genomes) to ensure that all genomes benchmarked within KEGG are called in the results. Further, export for this webserver output is not necessarily tabular or tabular-compatible (i.e., HTML-embedded table) and, therefore, will require additional data tidying due to paralog-related row duplications (i.e., duplicate rows lack names, which may be particularly troublesome for tidying without specialized programmatic script development). Recently, KEGG Synteny database, queries of which also use K numbers for input (two or more at a time), has been expanded to include the entirety of genomes in KEGG.

A recently developed tool of particularly exciting functionality is *fast.genomics*, a tool built within the suite of PaperBLAST (**Fig. 8**) [55] and that uses the genomes of MicrobesOnline. This tool is one of very few that pairs the power of a sequence-based search in tandem with the inferential value of clustering analysis for pairs of distinct protein families (**Fig. 4**). Several databases specifically designed for biosynthetic gene cluster analyses also fall within tool subtype 3, allowing for multiple target sequences/families as input while restricting genomes included in the analyses. An example of this subtype is CAGECAT [56] (https://cagecat.bioinformatics.nl) (Fig. S17), which uses NCBI sequence identifiers for input (multiple) chosen by the user. While users can also select specific organisms or taxonomic clades (by Genus), the queries are somewhat limited and predefined by the server’s database. A final example of this tool subtype, FunCoup [57] (https://funcoup.org/search/) (Fig. S18), takes a meta-analytical approach to family-level analyses, aggregating and summarizing data supporting functional coupling between proteins across a limited number of genomes, phylogenetic profiling via InParanoid [58] being one of several evidentiary criteria.

**Figure 8.**
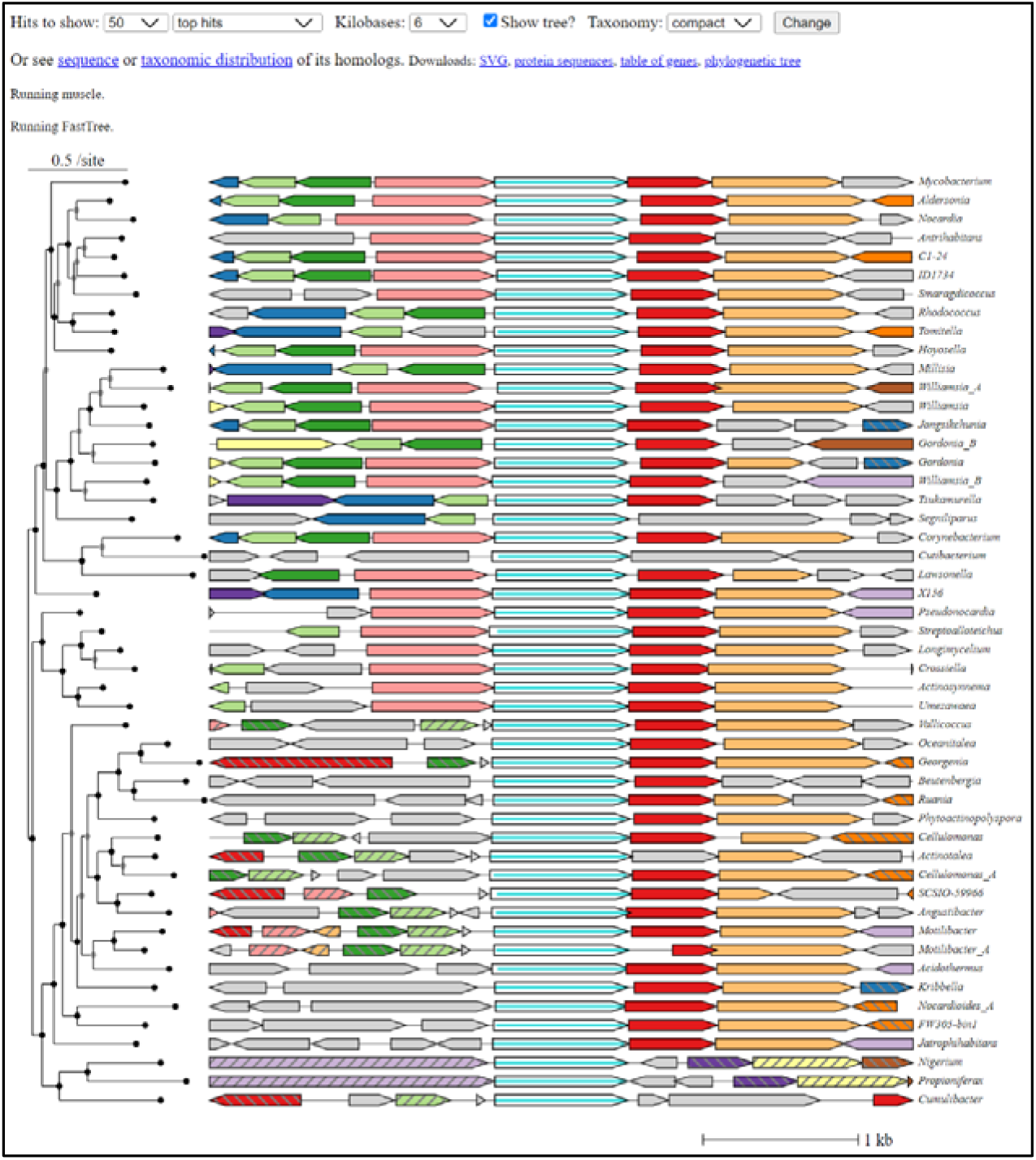
Graphical output of fast.genomics for one protein, Rv2230c (DUF34 family homolog, COG0327, P9WFM1 of Mycobacterium tuberculosis str. ATCC 25618 / H37Rv). A tree clustered view of physical clustering for closely related homologs is shown.

#### Subtype 4: custom genome selection, multiple target family selection

Subtype 4 tools are most commonly available in the form of online suites and custom account-linked workspaces (e.g., Galaxy [59] (https://usegalaxy.org/), BV-BRC (formerly, PATRIC)). Like Annotree, BV-BRC’s Comparative Systems tool is restricted taxonomically to bacteria and archaea [32] (https://www.bv-brc.org), but with the additional inclusion of viruses (Fig. S19-S20). The output of this tool includes a searchable heatmap for all identified gene families across a custom selection of genomes, the results of which can then be filtered using family identifiers (PGF IDs). MicroScope (the microbial platform of GenoScope) also possesses a Gene Phyloprofile tool. Multiple genomes (PkGDB, i.e., reannotated bacterial RefSeq genomes/proteomes) can be compared based on single or multiple genes/proteins, in addition to whole genome-to-genome phylogenomics. The ultimate result of this program is an output in the form of an HTML embedded table with each selected genome represented in a separate column (not row). Finally, JGI-IMG provides a tool suite that allows for the examination of custom genome lists with the use of many common systematic identifiers, such as KOs, COGs, and Pfams (i.e., “Find Function” feature of the suite). Again, the output for this tool is restricted to an HTML embedded table format but can be customized and exported in tabular format. In general, if all these tools are quite user-friendly and useful for first pass analyses, they are currently limited by the reliance on precomputed family annotations that can be partial, too broad, or simply incorrect [60].

One phylogenetic analysis tool of subtype 4 that is not dependent upon a user account-linked workspace is EggNOG’s Phylogenetic Profile tool [61] (http://eggnog6.embl.de/app/phyloprofile/) (Fig. S21). Uniquely, this browser-accessible resource allows for the input of multiple COGs and multiple user-selected genomes, the latter input being taxonomic identifiers. The submitted job results in a heatmap visualization showing absence-presence of selected orthologous groups across the user’s custom-selected genomes.

The Department of Energy Systems Biology Knowledgebase (KBase) has made analytic modules and pipelines available for researchers who lack programming skills [62]. Any KBase user account allows for browser-mediated access to complete suites of common bioinformatic analyses using either publicly accessible or user-uploaded data. Resources provided by KBase map out specially ordered “narratives” (i.e., an organized set of data objects and application queues within a digital notebook) for completing phylogenomic analysis starting from species trees, but such a pipeline can be unwieldy for novice users (Fig. S22; figure adapted from KBase 2020 phylogenetics narrative diagram). It should also be noted that analyses can take many hours depending on the number of genomes being analyzed, and such investments may be important timeline considerations for experimentalists.

Because of those described and many other challenges to tool usage, resources providing guidance appropriate for the microbiological field’s diverse audiences are increasingly necessary. Support of resources are often directly tied to the perceived use on part of the target community (frequently measured by citations, link traffic when reported to supporting agencies, organizations). Therefore, if we are to ensure the continued success and access to many high-quality resources, it is sometimes necessary to assist in the effective accessibility of their use by diverse userbases.

### Creating a Wiki compiling a non-exhaustive list of web-based resources organized into pedagogical modules for microbiologist

A persistent challenge exists within the bioinformatic community; that is, the ability to know which tools are available and which are most suitable for fulfilling our data analysis and visualization objectives. As time passes, more tools are published with others being decommissioned nearly at much the same rate, the longevity of most tools maintaining uncertainty across their lifetime due to funding instability [63]. In 2015, Attwood and colleagues found that after 18 years (1997-2015) over 60% of catalogued databases within DBcat (www.infobiogen.fr/services/dbcat) were “dead” (i.e., server was unresponsive, or search/other major functionality was not functioning) [63]. More recently, Kern and colleagues found, in a survey of 2618 tools published between 2019 and 2020, that 10% of them were unreachable, the rate of which they observed to increase linearly with earlier dates of publication [64]. A few sites have been dedicated to the aggregation of the totality of useful bioinformatics resources (e.g.: bio.tools [65], https://bio.tools; CNCB Database Commons [66], https://ngdc.cncb.ac.cn/databasecommons/; Nucleic Acids Research’s regular Database Issue [67]), but—in addition to being understandably challenging to maintain—the lack of grassroots- or leadership-level efforts to popularize some of these resources have left them of low findability and, therein, in deficit of broad use by the community. Only more recently have sites like CNCB Database Commons been recommended by the likes of *Cell Press* (https://marlin-prod.literatumonline.com/pb-assets/journals/research/cellpress/data/RecommendRepositories.pdf) or *Bioinformatics Advances* (https://academic.oup.com/bioinformaticsadvances/pages/instructions-to-authors). As of November 29, 2023, CNCB Database Commons indicated that 5213 of 6380 (82%) biological databases were annotated as being “alive”.

In response to our own difficulties in navigating the ever-changing frontier of bioinformatic tools, a wiki of webtools was established, initially, for our laboratory’s in-house use, and, later, was further developed with the intention of aiding other microbiologists (https://vdclab-wiki.herokuapp.com/). With 15 years of instructional experience in bioinformatics specifically for microbiologists, this resource was designed with our own graduate-level courses in mind, in addition to featuring some of our lab’s own commonly used bioinformatic workflows. The wiki was created using the pedagogic modules of already well-established bioinformatic courses and their learning objectives, which were used to model the website’s subsets of information, keywords, tags, and relationships between links. Additional pages provided in the main navigation like “VDC Favorites” and “Recent Finds” contribute a personal touch—one curated and the other a chronological, ongoing, and uncurated feed—to the website’s suite of information. Ideally, this custom collection of tools and workflows will aid other microbiological experimentalists that are less familiar with the user-friendly bioinformatic resources available today.

## Conclusions

Comprehensively extracting, synthesizing, and properly propagating scientific observations among databases, all in a manner that adheres to and further fosters the use of FAIR data guidelines remains a challenge [69]. Here, we examined the challenges pertaining to the capture of the literature on whole protein families. The curation of published data, alongside the interrogation of available tools common to this process, were surveyed and workflows incorporating different iterations of them were compared to provide a minimal workflow that can be followed by users to optimize search time (Fig. 1 and 5). Several potential pitfalls and stumbling blocks commonly encountered by researchers during biological publishing were identified and described, and further supplemented by examples of each using the case study of the DUF34 protein family. Importantly, it was observed that the choice of keywords and search engines, though equally important, vary both together and independently in how they influence published data capture results. Additionally, false positives across search engine types illustrated the importance of thorough, well-informed curation efforts, and the need for more stringent standards among publishers. Related and although tools that allow this form of search are limited in number, sequence-based searches were determined to be critical first steps of the data capture process at the protein family level. A comparison of merged query result lists derived from varying numbers of single-sequence-based searches (i.e., 1-sequence, 2-sequences, and 7-sequences) to those of PaperBLAST’s HMM-based search tool demonstrated that, despite being drawn from the same publication-bioentity/sequence ID crosslink network, each method—no matter the number of representative sequences used to generate a consolidated list for the sequence-based results—provided yields dissimilar in total and quality. The greatest distinctions between lists were observed between the sequence-based list derived from only one family member sequence of *E. coli* and that of the HMM-based results. Interestingly, the yield increase of single-sequence-based queries was shown to have largely plateaued between the lists derived from one sequence to two sequences, with only a marginal increased yield of publications between lists derived from two and seven sequences. These results implied that the most impactful factor for improving single-sequence-based results was not necessarily the total number of sequences used to compile the final results list. Moreover, and consistent with this hypothesis, the member sequence representing the known most-divergent domain architecture of the DUF34 protein family, that of *B. cereus*, was also the sequence to derive the greatest number of publications unique to its query when comparing the individual single-sequence-based yields. Consideration of the QCC cycle approach in these comparisons further emphasized that, while no single method appeared optimal, it was this more tedious method that was observed to outperform all others in total unique publications and rate of false positives, the latter of which was largely driven by the curation-based nature of the QCC cycle approach. As it relates to the interest of increasing the efficiency with which a given experimentalist might capture a sufficient portion of literature relevant to a protein family, these analyses suggested that the HMM-based method available through PaperBLAST would likely best serve this purpose as it was the approach that retrieved the most unique and relevant publications with the least amount of time and resource investment.

Because of the importance of protein family-level information in guiding the published data capture process, a survey of web-based phylogenomic and phylogenetic tools was performed highlighting those of higher usability and interoperability. Mapping identifiers between databases is central to comparative genomics approaches as users often must use a variety of resources with distinct features and formats in order to accomplish their analytic objectives. However, the more work and time necessary to transform and/or map data between resources, the less likely they are to be used together or in tandem, regardless of how high-quality their visualizations or how well-curated their data [70–74]. Although not thoroughly discussed, interoperability was observed to be notably bereft between the phylogenetic/phylogenomic tools surveyed, as well as with those of other types of biological databases. While it has been argued to simultaneously contribute to the larger systemic problem, the growing diversity of tools also suggests that the challenges driven by persisting interoperability deficits between resources have yet to be sufficiently ameliorated and continue to obstruct analytic processes. That is, publishing behavior of biological tools suggests that many researchers continue to resort to “reinventing the wheel” instead of developing tools to bridge the gaps between different resources or, alternatively, work with existing resources to improve their interoperability. To confront these trends in our recommendations to experimentalists, we designed a pedagogical wiki of bioinformatic workflows using tools with long, well-tried histories of continued support, adequate quality curation, and maintenance. Ultimately, it is our hope that this work provides a framework with which experimental microbiologists can perform routine operations on genes and proteins of a given family even as the genomic data size increases.

## Author statements

### Author contributions

Conceptualization: Valérie de Crécy-Lagard; Data Curation: Colbie J. Reed; Formal Analysis: Colbie J. Reed; Funding Acquisition: Valérie de Crécy-Lagard; Investigation: Colbie J. Reed; Methodology: Colbie J. Reed, Valérie de Crécy-Lagard; Project Administration: Valérie de Crécy-Lagard, Colbie J. Reed; Resources: Valérie de Crécy-Lagard; Software: Remi Denise; Supervision: Valérie de Crécy-Lagard; Validation: Colbie J. Reed, Valérie de Crécy-Lagard; Visualization: Colbie J. Reed; Writing – Original Draft: Colbie J. Reed, Valérie de Crécy-Lagard; Writing – Review & Editing: Colbie J. Reed, Valérie de Crécy-Lagard, Geoffrey Hutinet, Jill Babor, Rémi Denise; Tool/Resource Curation, Review *for Website Collections: Colbie J. Reed, Rémi Denise, Jacob Hourihan, Jill Babor, Marshall Jaroch, Maria Martinelli, Geoffrey Hutinet, and Valérie de Crécy-Lagard*.

## Acknowledgements

We thank Marc Chevrette and Svetlana Gerdes for critical reading and suggestions and all students and participants in the training and courses given by the Crécy lab in the last 20 years for spurring the idea of this manuscript.

