## Supplemental Materials Document for "Beyond Blast: Enabling Microbiologists to Better Extract Literature, Taxonomic Distributions and Gene Neighborhood Information for Protein Families"

**Index**

**Headings for each section of supplementary content are hyperlinked to facilitate document navigation.*

**Supplementary Data**

**S1.** All keywords accumulated for DUF34 protein family during the comprehensive published data capture process.

**S2.** Curated sets of tools organized by: (a) orthology-/homology-limited and phyletic patterning tools; (b) gene neighborhood; and (c) synteny tools.

**S3.** All curated yields of specialized corpus search tools: PubMed, EuropePMC, PubTator, Scinapse.

**S4.** Simple Comparison of Text-based vs. Sequence-based Publication Retrieval.

**S5.** Publication Retrieval Method Comparison — Single-sequence vs. HMM-based PaperBLAST Methods.

**S6.** Venn Diagram Data for the Comparison of the Two Major PaperBLAST Retrieval Methods.

**S7.** Publication Retrieval Method Comparison — Single-sequence vs. HMM-based vs. Idealized "QCC" Cycle Method.

**S8.** Venn Diagram Data Comparing PaperBLAST Retrieval Methods to "QCC Cycle" Approach.

**S9.** Comparison of the PaperBLAST Results of Several, Disparately Related DUF34 Homolog Sequences.

**S10.** Final Summary Venn Diagram of all Methods' Results Relevant to DUF34.

**Supplementary Tables**

**S1.** All 65 gene families characterized by the Laboratory of Valérie de Crécy-Lagard, PhD.

**S2.** Results of investigation comparing sequence-based search tool PaperBLAST (two methods) to those of the idealized “QCC” method.

**Supplementary Elaborations**

**S1.** Explanation of XynX “productive ignorance” example.

**Supplementary Figures**

**S1.** Seed Information Assessment: establish understanding of starting information in preparation for the capture of published data.

**S2.** Retrieve sequences using known names/aliases.

**S3.** Family-level analyses.

**S4.** Example of over-propagation of root family attribution in EggNOG Database as a result of fusions.

**S5.** WorldWideScience.org-, ScienceResearch.com-generated diagrams of keywords.

**S6.** Coincidental homonyms are challenges for text-based search engines, even those as specialized as PubTator.

**S7.** FlaGs example output.

**S8.** EFI-GNT example output.

**S9.** Annotree example: single family, PF02591, in bacteria.

**S10.** COGNAT example output.

**S11.** SubtiWiki (CoreWiki) beta feature of the database’s Genomic Neighborhood Comparison viewer of the DUF34 homolog of B. subtilis, YqfO (BSU_25170).

**S12.** Example tree output of MicrobesOnline.

**S13.** Screengrab demonstrating different “filters” for target family recognition, particularly used differentially with multiple targets (MicrobesOnline).

**S14.** Screengrab demonstrating navigation to family-specific trees in MicrobesOnline via the “Gene Info” tab of each entry.

**S15.** “Families” (COGs) in STRING search tool options.

**S16.** KO identifiers being supplied to KEGG orthology tool (multiple families, limited genome customization).

**S17.** Example CAGECAT output using DUF34 and COG1579 homolog sequences: MSMEG_4307 (NCBI-ProteinID: ABK73705); MSMEG_4306 (NCBI-ProteinID: ABK70599).

**S18.** Example output for FunCoup using “YbgI” of *E. coli*.

**S19.** BV-BRC protein family sorter, general output example (a) with additional viewer examples.

**S20.** BV-BRC protein family sorter output, Families.

**S21.** EggNOG Phylogenetic Profile tool example output using COG0327, COG3323, and COG1579.

**S22.** KBase workflow for generating a “gene tree” (as regarded by KBase) from several select genomes.

**S23.** Genomic Context Visualizer example output.

**S24.** GizmoGene example output.

**Supplementary References**

### Supplementary Data

**Supplementary data files in submission packet are also accessible via the accompanying FigShare.*

Data S1. All keywords accumulated for DUF34 protein family during the comprehensive published data capture process.

Data S2. Curated sets of tools organized by: (a) orthology-/homology-limited and phyletic patterning tools; (b) gene neighborhood; and (c) synteny tools.

**Data S3.** All curated yields of specialized corpus search tools: PubMed, EuropePMC, PubTator, Scinapse. Each search engine was queried using the 10 selected keywords thought to represent the most commonly associated names/aliases associated with the target protein family (i.e., “NGG1 interacting factor 3”, “NIF3”, “NIF3L1”, “GTP cyclohydrolase 1 type 2”, “DUF34”, “YbgI”, “PF01784”, “COG0327”, “YqfO”, and “COG3323”). Respective yields were exported from each engine’s results and, subsequently, manually reviewed for relevance rated at three levels: 1) “Focal”, a true positive hit in which the target homolog was mentioned in either the title and/or the abstract; 2) “Non-focal”, a true positive hit in which the target homolog, while not mentioned in the title or abstract, was mentioned in another subsection or supplemental data of the publication; and 3) “False Positive”, a curated and confirmed false positive hit in which the target homolog was falsely identified among any of the yielded publication’s text. These three ratings constitute the three left-most columns of each subset of exported hits per search engine (respective publication’s row assigned a “yes” in the column corresponding to one of the three rating categories for which it was found affirmative, reflecting the curation results or relevance status). Tables reflecting the curation summaries of each raw export (i.e., each keyword/query per search engine) were generated to document curation process (see “Export Subsets, Curation”). The total number of raw yields varied per search engine due to the variable productivity of individual engines and select keywords within those search engines. Raw yields (i.e., returned publications) for each keyword per search engine were also included in this supplementary dataset, accompanying the export summaries; queries returning no hits were unable to be exported and, therefore, are not represented within a raw file (see export summaries for which queries resulted in no hits, and, therein, which query yields were left out of the raw data files; Data S3 table includes a key for all raw export files). The results of this survey are illustrated in **Figure 4** of the main text.

### Supplementary Tables

**Table S1.** All 65 gene families characterized by the Laboratory of Valérie de Crécy-Lagard, PhD. (Asterisks indicate homologs in human)

| **Pathway** | **Name** | **UniProt Representative** | **COG/DUF/PFAM** | **PMID** |
| --- | --- | --- | --- | --- |
| **RNA Modification** | | | | |
| Agmitidine Synthase | TiaS | O59476 | COG1571 | 18844986; 20139989 |
| Archaeosine Synthesis | QueF-like | A3MSP1 | COG0780 QueF-paralog | 22032275; 28383498 |
| Archaeosine Synthesis | Gat/QueC fusion | Q981C9 | COG0449/COG0603 | 22032275 |
| Archaeosine Synthase | ArcS | Q58428 | COG1549, DUF5591, TGT paralog | 20129918 |
| 5-methylaminomethyl-2-thiouridine synthase | MnmC1/C2 fusion | P77182 | COG4121/COG0579 | 17673083 |
| Pre-Q_0_ Reductase (Queuosine Biosynthesis | QueF | Q46920 | COG2904/0780 | 14660578; 15767583 |
| Queuosine/Archaeosine Biosynthesis | QueC | O31675 | COG0603 | 14660578 |
| Queuosine/Archaeosine Biosynthesis | QueD | O31676 | COG0720 | 14660578 |
| Queuosine/Archaeosine Biosynthesis | QueE | O31677 | COG0602 | 14660578 |
| Queuosine Biosynthesis | QueH | Q9WZJ0 | DUF208 | 28128549 |
| PreQ0/preQ1 transport | YhhQ | P37619 | COG1738 | 28208705 |
| Queuosine (Q) hydrolase | CD1682/QueK | Q186N9 | COG1957 paralog | 31481610 |
| Queuine lyase | CD1684/QueL | Q186P0 | COG1244 paralog | 31481610 |
| Q transport | CT140 | O84142 | COG1738 paralog | 31481610 |
| Q Synthesis | CT193 | O84196 | COG0343 paralog | 31481610 |
| Queuosine Salvage | Qng1* | Q9HDZ9 | DUF2419 | 24911101; 36610787 |
| tRNA-ac4C Biosynthesis | TmcA | D4GW73 | COG1444 | 19478918 |
| tRNA-m1A22 Biosynthesis | TrmK | P54471 | COG2384 | 18420655; 17852564 |
| tRNA-m1Psi54 synthesis | PusY | D4GTL8 | COG1901/DUF358 | 22274953; |
| tRNA-t6A37 Synthesis (Universal) | TsaC * | P45748 | COG0009 | 19287007 |
| tRNA-t6A37 Synthesis (Universal) | TsaC2, Sua5 | P32579 | PF03481 | 19287007 |
| tRNA-t6A37 Synthesis (Bacteria) | TsaB | O05516 | COG1214 | 21285948 |
| tRNA-t6A37 Synthesis (Bacteria) | TsaD | P05852 | COG0533 | 21285948 |
| tRNA-t6A37 Synthesis (Bacteria) | TsaE | P0AF67 | COG0802 | 22378793 |
| tRNA-t6A37 Synthesis (Eukarya/Archaea Cyto) | Kae1* | P36132 | PF00814 | 21285948 |
| tRNA Dihydrouridine Synthase (Bacteria) | DusABC* | P32695 | COG0042 | 11983710 |
| Wybutosine Biosynthesis | TYW1, WyeA* | Q08960 | PF00258/PF04055/PF08608 | 16162496 |
| Wyosine Derivative Biosynthesis (Archaea) | Taw22 | Q9V2G1 | COG2520 | 20382657 |
| tRNA Psi 32/32 Synthase (*B. subtilis*) | YjbO | O31613 | COG0564 paralog | 32629984 |
| rRNA Psi Sythase (*B. subtilis*) | YhcT | P54604 | COG0564 paralog | 32629984 |
| tRNA m6t6A | TrmO* | P28634 | COG1720 | 25063302 |
| **DNA Modification** | | | | |
| dADG modification | DpdA | A0A3A3JBR7 | COG0343 TGT paralog | 30159947 |
| dADG modification | DpdB | A0A3A3NHD9 | pfam14072 DndB paralog | 30159947 |
| dADG modification | DpdC | A0A3A3IHU3 | DUF308 | 30159947 |
| **Small Molecule/Cofactor Metabolism** | | | | |
| Folate Synthesis (Bacteria, FolB Shunt) | PTPS-III | A0A098MZ39 | COG0720 paralog | 19395485 |
| Folate FolC-like (Chlamydiae) | CT611 | O84617 | COG1478 | 17645794 |
| GTP Cyclohydrolase I type 2 | FolE2 | P94398 | COG1469 | 17032654 |
| Folate, New PABA Synthesis | CT610 | O84616 | COG5424 | 25006229 |
| Iron-sulfur Cluster Repair | YgfZ* | P0ADE8 | COG0354, GcvT paralog | 20489182 |
| Pterin Biosynthesis (Archaea) | FolB2/MptD | Q57851 | COG2098/DUF372/381 | 22931285 |
| Pterin Biosynthesis (Archaea) | FolK2/MptE | Q59028 | COG1634/DUF115 | 22931285 |
| Tetrahydromonapterin Synthesis | FolX | P0AC19 | COG1539 | 19897652 |
| Tetrahydromonapterin Synthesis | FolM | P0AFS3 | COG1028 paralog | 19897652 |
| Thiamine Metabolism | TenAE | Q9ASY9 | COG0812 | 25014715 |
| Zinc Homeostasis | YeiR* | P33030 | COG0523 paralog | 15690043 |
| R,S SAM hydrolase | Sare_1364 | A8M783 | DUF62 | 32776704 |
| Oxoproline Metabolism | YbgJ | P0AAV4 | COG20149 | 28830929 |
| Oxoproline Metabolism | YbjK | P75745 | COG1984 | 28830929 |
| Oxoproline Metabolism | YbgL | P75746 | COG1540 paralog | 28830929 |
| Riboflavin | YbiA/RibX | P30176 | COG3236 | 25431972 |
| Thiamine Metabolism | At4g29530 | Q9SU92 | PF06888 paralog | 26537753 |
| Thiamine Metabolism (Plants) | TenAC | F4KFT7 | IPR036412/IPR016084 /IPR004305 | 27677881 |
| Phosphopanteine Hydrolase | At4g32180* | Q8L5Y9 | DUF89_II (PanK fusion) | 27322068 |
| B12 Metabolism | PF0295 | Q8U404 | COG2103/DUF71 paralog | 23013770 |
| PLP Homeostasis | YggS/PLPHP/PLPBP* | P67080 | COG0325 | 26872910 |
| **Protein Modification** | | | | |
| Beta-Lysylation of Elongation Factor P | YjeA | P0A8N7 | COG226 paralogs | 20070887; 21841797 |
| Beta-Lysylation of Elongation Factor P | YjeK | P39280 | COG0535 paralogs | 20070887; 21841797 |
| Diphthamide Biosynthesis | Dph6* | Q12429 | COG2103/DUF71 paralog | 23013770 |
| Deoxyhypusine Biosynthesis | HVO_2299 | D4GWG4 | COG0010 paralogs | 28053595 |
| **Carbon Source Catabolism** | | | | |
| Mannitol-phosphate Dehydrogenase/Phosphatase | MtlD | Q6FBP5 | PF13419/PF08125 | 24800891 |
| Sugar-phosphate and Nucleotide Hydrolase | PH1575 | O59272 | DUF89_I | 27322068 |
| Sugar-phosphate Hydrolase | YMR027W | Q04371 | DUF89_III | 27322068 |
| 3-Oxo-tertronate Kinase | YgbK | Q46889 | DUF1537 paralog | 27402745; 27294475 |
| D-Threonate Kinase | DtnK | Q8ZRS5 | DUF1537 paralog | 27402745; 27294475 |
| D-Threonate 4-phosphate Dehydrogenase | PdxA2 | P58718 | COG1995 paralog | 27402745; 27294475 |
| **Unpublished (papers in-prep)** | | | | |
| nm5U34 Synthase | YtqA | O35008 | COG1242 |  |
| nm5U34 Methylase | YtqB | O34614 | COG2519 |  |
| dPreQ1 Methylase | DpdM | M4PNV1 | PF03692 (paralog?) | <https://biorxiv.org/cgi/content/short/2023.04.13.536721v1> |
| dPreQ1 Formyltransferase | DpdN | NA, ncbi: YP_010114479.1 | PF00551 paralog | <https://biorxiv.org/cgi/content/short/2023.04.13.536721v1> |
| CDG Decarboxylase | DpdL | M4SNA7 | COG0720 paralog | <https://biorxiv.org/cgi/content/short/2023.04.13.536721v1> |
| preQ_0_/preQ_1_ Transporter | Bifidobacterium breve CREST homolog | A0A0M3T8W5 | PF03006/COG1272 paralog |  |
| preQ_0_/preQ_1_ Transporter | *Bartonella henselae* MFS homolog | WP_011180872.1 | PF07690/COG2814 paralog |  |
| preQ_0_/preQ_1_/q | *Acidobacteria bacterium* DMT homolog | A0A2V9U0M9 | PF07168 paralog |  |

**Table S2.** Results of investigation comparing sequence-based search tool PaperBLAST (two methods) to those of the idealized “QCC” method. “Focal” and “False Positive” publication labels have been bolded in the table.

|  | **Method** | | | |
| --- | --- | --- | --- | --- |
|  | **PaperBLAST** | | | **QCC** |
|  | **HMM** | **Single Sequence/UniProt ID** | |  |
| **Relevance** | **PF01784** | ***H. sapiens* (Q9GZT8)** | ***E. coli* (P0AFP6)** |  |
| Non-Focal |  |  |  | Ahmed 2011 |
| **Focal** | Akiyama 2003 | Akiyama 2003 |  | Akiyama 2003 |
| Non-Focal |  |  |  | Alalouf 2011 |
| Non-Focal | Alam 2011 | Alam 2011 | Alam 2011 | Alam 2011 |
| Non-Focal | Anderson 2010 | Anderson 2010 |  |  |
| Non-Focal |  |  |  | Alderman 2019 |
| Non-Focal |  |  |  | Alqazlan 2020 (thesis) |
| Non-Focal |  |  |  | Amon 2010 (thesis) |
| Non-Focal |  |  |  | Antelmann 2005 |
| Non-Focal |  |  |  | Antoniali 2017 |
| Non-Focal | Araújo 2020 | Araújo 2020 |  | Araújo 2020 |
| Non-Focal |  |  |  | Ashburner 1999 |
| Non-Focal |  |  |  | Aurass 2009 |
| Non-Focal |  |  |  | Avican 2021 |
| **Focal** |  |  |  | Bagautdinov 2008 |
| Non-Focal |  |  |  | Bai 2017 |
| Non-Focal |  |  |  | Bashir 2020 |
| Non-Focal |  |  |  | Baysal 2013 |
| Non-Focal | Belvin 2019 | Belvin 2019 |  | Belvin 2019 |
| Non-Focal |  |  |  | Bermudez 2015 |
| Non-Focal |  |  |  | Bhasin 2008 |
| Non-Focal |  |  |  | Bhatraju 2015 |
| Non-Focal | Bleichert 2020 | Bleichert 2020 |  | Bleichert 2020 |
| Non-Focal |  |  |  | Boot 2016 |
| Non-Focal |  |  |  | Bootsma 2010 (patent) |
| Non-Focal |  |  |  | Boswell 2018 |
| Non-Focal |  |  |  | Breker 2014 |
| Non-Focal |  |  |  | Brosnahan 2019 |
| Non-Focal |  |  |  | Brosnahan 2020 (thesis) |
| **False Positive** | Brady 1989 |  | Brady 1989 |  |
| Non-Focal |  |  |  | Bruce 2012 (patent) |
| Non-Focal |  |  |  | Burnstein 2016 |
| Non-Focal | Byrne 2014 |  | Byrne 2014 | Byrne 2014 |
| Non-Focal |  |  |  | Camp 2016 |
| Non-Focal |  |  |  | Cardenas 2009 |
| Non-Focal |  |  |  | Chang 2016 |
| Non-Focal |  |  |  | Charlton 2015 |
| Non-Focal |  |  |  | Chauhan 2015 |
| Non-Focal |  |  |  | Chen 1999 |
| **Focal** |  |  |  | Chen 2010a |
| Non-Focal |  |  |  | Chen 2010b |
| **Focal** | Chen 2014 | Chen 2014 | Chen 2014 | Chen 2014 |
| Non-Focal |  |  |  | Cherkas 2016 |
| Non-Focal |  |  |  | Chiesa 2020 |
| **Focal** | Choi 2013 | Choi 2013 |  | Choi 2013 |
| Non-Focal |  |  |  | Chung 2013 |
| Non-Focal |  |  |  | Codrich 2019 |
| Non-Focal |  |  |  | Comas 2008 |
| Non-Focal |  |  |  | Conn 2017 |
| **Focal** |  |  |  | Constantine 2009 |
| Non-Focal | Cox 2020 |  | Cox 2020 |  |
| Non-Focal | Crapoulet 2006 | Crapoulet 2006 |  | Crapoulet 2006 |
| Non-Focal |  |  |  | Cury 2020 |
| Non-Focal | Czubat 2020 | Czubat 2020 |  |  |
| Non-Focal | Da Costa 2018 | Da Costa 2018 |  | Da Costa 2018 |
| Non-Focal |  |  |  | Daas 2016 |
| Non-Focal |  |  |  | Daas 2018 |
| Non-Focal |  |  |  | Dalia 2011 (thesis?) |
| Non-Focal |  |  |  | Damianou 2020 |
| Non-Focal | Danelishvili 2005 | Danelishvili 2005 |  | Danelishvili 2005 |
| Non-Focal |  |  |  | Daou 2019 |
| Non-Focal |  |  |  | Dapas 2019 (thesis) |
| Non-Focal |  |  |  | Dartigalongue 2001 |
| Non-Focal |  |  |  | Degnan 2005 |
| Non-Focal |  |  |  | DeLoney 2002 |
| Non-Focal | De Sordi 2015 | De Sordi 2015 |  |  |
| Non-Focal | Díaz-Mejía 2009 |  | Díaz-Mejía 2009 | Díaz-Mejía 2009 |
| Non-Focal |  |  |  | Dickey 2013 (thesis) |
| Non-Focal |  |  |  | Ding 2015 |
| Non-Focal |  |  |  | Dionisio 2016 |
| Non-Focal |  |  |  | Ditse 2017 |
| Non-Focal | Dong 2008 | Dong 2008 |  | Dong 2008 |
| Non-Focal |  |  |  | Drummelsmith 2007 |
| Non-Focal |  |  |  | Ducret 2021 |
| Non-Focal |  |  |  | Dudin 2017 |
| Non-Focal |  |  |  | Duncan 2017 |
| Non-Focal |  |  |  | Dunman 2001 |
| Non-Focal |  |  |  | Duzyj 2014 |
| Non-Focal |  |  |  | Edwards 2013 |
| Non-Focal |  |  |  | El Bounkari 2009 |
| Non-Focal | Esser 2008 |  | Esser 2008 | Esser 2008 |
| Non-Focal |  |  |  | Facciuolo 2013 |
| Non-Focal |  |  |  | Facciuolo 2016 |
| Non-Focal |  |  |  | Falkenberg 2019 |
| Non-Focal |  |  |  | Fang 2010 |
| Non-Focal | Fenner 2019 | Fenner 2019 |  | Fenner 2019 |
| Non-Focal |  |  |  | Fontes 2018 (thesis) |
| Non-Focal |  |  |  | Freiberg 2016 |
| **Focal** | Fujishiro 2014 | Fujishiro 2014 | Fujishiro 2014 | Fujishiro 2014 |
| Non-Focal |  |  |  | Fujishiro 2015 |
| Non-Focal |  |  |  | Fujishiro 2016 |
| Non-Focal |  |  |  | Fukushima 2020 |
| Non-Focal | Furuta 2010 |  | Furuta 2010 | Furuta 2010 |
| Non-Focal |  |  |  | Galperin 2010 |
| Non-Focal | Gangaiah 2013 |  | Gangaiah 2013 |  |
| Non-Focal | Gangaiah 2014 |  | Gangaiah 2014 | Gangaiah 2014 |
| Non-Focal |  |  |  | Gao 2019 |
| Non-Focal | Gaudet 2011 |  | Gaudet 2011 |  |
| Non-Focal | Gawinecka 2012 | Gawinecka 2012 |  | Gawinecka 2012 |
| Non-Focal |  |  |  | Geislet 1992 |
| Non-Focal |  |  |  | Geyer 2019 |
| Non-Focal | Ghaemmaghami 2003 | Ghaemmaghami 2003 |  | Ghaemmaghami 2003 |
| Non-Focal |  |  |  | Ghersi 2011 |
| Non-Focal |  |  |  | Giannangelo 2018 (thesis) |
| Non-Focal |  |  |  | Gifford 2000 |
| Non-Focal |  |  |  | Giotti 2019 |
| Non-Focal |  |  |  | Giovannelli 2017 |
| Non-Focal |  |  |  | Girinathan 2018 |
| Non-Focal |  |  |  | Giuffrida 2006 |
| **Focal** | Godsey 2007 | Godsey 2007 | Godsey 2007 | Godsey 2007 |
| Non-Focal |  |  |  | Gooderham 2008 (thesis) |
| Non-Focal |  |  |  | Graf 2018 |
| Non-Focal | Gupta 2017 | Gupta 2017 |  | Gupta 2017 |
| Non-Focal |  |  |  | Gupta 2018 |
| Focal |  |  |  | Hadano 2001 |
| Non-Focal |  |  |  | Hadj-Hamou 2010 (thesis) |
| Non-Focal | Han 2006 |  | Han 2006 | Han 2006 |
| Non-Focal | Happonen 2019 | Happonen 2019 | Happonen 2019 | Happonen 2019 |
| Non-Focal |  |  |  | Harrison 2021 |
| Non-Focal |  |  |  | Hart 2018 |
| Non-Focal |  |  |  | Hayles 2013 |
| Non-Focal |  |  |  | He 2019 |
| Non-Focal | Hendrickson 2010 | Hendrickson 2010 |  |  |
| Non-Focal |  |  |  | Hermans 2008 (patent) |
| Non-Focal |  |  |  | Hews 2019 |
| Non-Focal |  |  |  | Hladik 2019 |
| Non-Focal | Ho 2015 | Ho 2015 |  | Ho 2015 |
| Non-Focal |  |  |  | Ho 2016 |
| Non-Focal |  |  |  | Hossain 2021 |
| Non-Focal | Huang 2015 | Huang 2015 | Huang 2015 |  |
| Non-Focal |  |  |  | Huang 2019 |
| Non-Focal |  |  |  | Huston 2017 (lecture) |
| Non-Focal |  |  |  | Huttlin 2010 |
| Non-Focal |  |  |  | Iyer 2016 |
| Non-Focal |  |  |  | Jackson 2013 |
| Non-Focal |  |  |  | Jama 2014 (thesis) |
| Non-Focal |  |  |  | Jiang 2013 |
| Non-Focal | Jiang 2023 | Jiang 2023 | Jiang 2023 |  |
| Non-Focal |  |  |  | Johnson 2018 |
| Non-Focal |  |  |  | Johnson 2019 (preprint) |
| Non-Focal |  |  |  | Jones 2006 |
| Non-Focal |  |  |  | Jostes 2019 (thesis) |
| Non-Focal | Joshi 2020 |  | Joshi 2020 |  |
| Non-Focal |  |  |  | Jothi 2006 |
| Non-Focal |  |  |  | Kai 2016 |
| Non-Focal |  |  |  | Kalari 2013 |
| **Focal** | Kanesaki 2021 | Kanesaki 2021 |  |  |
| **Focal** |  |  |  | Karniely 2006 |
| Non-Focal | Kawabata 2019 | Kawabata 2019 |  |  |
| Non-Focal |  |  |  | Kaysser 2010 |
| Non-Focal |  |  |  | Kerns 2020 (thesis) |
| Non-Focal |  |  |  | Kim 2018 |
| Non-Focal | Kölbel 2019 | Kölbel 2019 |  | Kölbel 2019 |
| Non-Focal | Kolker 2004 |  | Kolker 2004 | Kolker 2004 |
| **Focal** | Kuan 2013 | Kuan 2013 | Kuan 2013 | Kuan 2013 |
| Non-Focal |  |  |  | Kumar 2018 |
| Non-Focal |  |  |  | Kusonmano 2018 |
| Non-Focal |  |  |  | Kwon 2005 |
| Non-Focal | Labandeira-Rey 2009 |  | Labandeira-Rey 2009 | Labandeira-Rey 2009 |
| **Focal** | Ladner 2003 |  | Ladner 2003 | Ladner 2003 |
| **Focal** |  |  |  | Lamba 2013 (poster abstract) |
| Non-Focal | Lasserre 2006 |  |  |  |
| Non-Focal | Leang 2005 | Leang 2005 | Leang 2005 | Leang 2005 |
| Non-Focal |  |  |  | Lee 2008 (thesis) |
| Non-Focal | Li 2010 | Li 2010 | Li 2010 | Li 2010 |
| Non-Focal | Li 2017 | Li 2017 |  | Li 2017 |
| Non-Focal | Liang 2013 | Liang 2013 | Liang 2013 | Liang 2013 |
| Non-Focal |  |  |  | Lie 2013 |
| Non-Focal |  |  |  | Lin 2004 |
| Non-Focal |  |  |  | Lin 2019 |
| Non-Focal |  |  |  | Liu 2013 |
| Non-Focal |  |  |  | Liu 2014 |
| Non-Focal |  |  |  | Liu 2019 |
| Non-Focal |  |  |  | Lively 2010 |
| Non-Focal | Livengood 2008 |  | Livengood 2008 | Livengood 2008 |
| Non-Focal |  |  |  | Long 2020 |
| Non-Focal |  |  |  | López-Madrigal 2013 |
| Non-Focal |  |  |  | Lu 2015 |
| Non-Focal |  |  |  | Luo 2005 (thesis) |
| Non-Focal |  |  |  | Luo 2007 |
| Non-Focal |  |  |  | Lv 2021 |
| Non-Focal |  |  |  | Mahdi 2013 |
| Non-Focal |  |  |  | Makarova 2010 |
| Non-Focal |  |  |  | Malan-Müller 2020 |
| **Focal** |  |  |  | Malik 2020 (preprint) |
| Non-Focal | Malik-Kale 2008 | Malik-Kale 2008 | Malik-Kale 2008 | Malik-Kale 2008 |
| **Focal** | Manan 2018 | Manan 2018 | Manan 2018 | Manan 2018 |
| Non-Focal |  |  |  | Manzano-Marin 2019 |
| Non-Focal | Manzourolajdad 2015 | Manzourolajdad 2015 |  | Manzourolajdad 2015 |
| **Focal** | Martens 1996 | Martens 1996 |  | Martens 1996 |
| Non-Focal |  |  |  | Martin 2002 |
| Non-Focal |  |  |  | Martin 2018 |
| Non-Focal |  |  |  | Martinot 2018 |
| Non-Focal |  |  |  | Massart 2015 |
| Non-Focal |  |  |  | McBrearty 2019 (thesis) |
| Non-Focal |  |  |  | McKee 2018 |
| Non-Focal |  |  |  | McKenzie 2012 |
| Non-Focal | McKenzie 2015 | McKenzie 2015 |  | McKenzie 2015 |
| **False Positive** | McLeod 2017 | McLeod 2017 |  |  |
| Non-Focal | Medrano 2009 |  | Medrano 2009 | Medrano 2009 |
| Non-Focal |  |  |  | Meibom 2004 |
| Non-Focal | Melian 2021 | Melian 2021 | Melian 2021 |  |
| **Focal** |  |  |  | Merla 2004 |
| Non-Focal |  |  |  | Meshram 2019 |
| Non-Focal |  |  |  | Minias 2015 |
| Non-Focal |  |  |  | Minias 2018 |
| Non-Focal |  |  |  | Moreira 2008 |
| Non-Focal | Moreno 2018 | Moreno 2018 |  | Moreno 2018 |
| Non-Focal |  |  |  | Morgenstern 2017 |
| Non-Focal |  |  |  | Munoz 2020 |
| Non-Focal |  |  |  | Nalls 2009 |
| Non-Focal | Naqvi 2015 |  | Naqvi 2015 | Naqvi 2015 |
| Non-Focal | Naqvi 2018 |  | Naqvi 2018 |  |
| Non-Focal |  |  |  | Nelson 2007 |
| Non-Focal | Neville 2020 | Neville 2020 | Neville 2020 |  |
| Non-Focal |  |  |  | Niehaus 2015 |
| Non-Focal |  |  |  | Nowlan 2020 |
| Non-Focal |  |  |  | Nowlan 2021 |
| Non-Focal |  |  |  | O'Connor 2012 |
| Non-Focal |  |  |  | Ogura 2010 |
| Non-Focal |  |  |  | Ogura 2016 |
| Non-Focal |  |  |  | Ogura 2018 |
| **Focal** |  |  |  | Ogura 2019 |
| Non-Focal |  |  |  | O'Hanlon 2011 |
| Non-Focal |  |  |  | Oja 2018 |
| Non-Focal | Page 2005 | Page 2005 |  | Page 2005 |
| Non-Focal |  |  |  | Pan 2012 |
| Non-Focal | Parente 2016 | Parente 2016 |  | Parente 2016 |
| Non-Focal |  |  |  | Park 2007 |
| Non-Focal |  |  |  | Passalacqua 2012 |
| Non-Focal |  |  |  | Patel 2015 |
| Non-Focal |  |  |  | Pattrick 2019 |
| Non-Focal | Paul 2017 | Paul 2017 |  | Paul 2017 |
| Non-Focal |  |  |  | Peng 2019 |
| Non-Focal | Pereira 2005 | Pereira 2005 |  |  |
| Non-Focal | Pereira 2011 | Pereira 2011 |  | Pereira 2011 |
| Non-Focal |  |  |  | Perez-Samper 2018 |
| Non-Focal |  |  |  | Phuphisut 2018 |
| Non-Focal | Pompesiello 2001 |  | Pompesiello 2001 | Pompesiello 2001 |
| Non-Focal | Porcelli 2013 | Porcelli 2013 | Porcelli 2013 | Porcelli 2013 |
| Non-Focal |  |  |  | Prahlad 2020 |
| Non-Focal | Prunetti 2016 |  | Prunetti 2016 | Prunetti 2016 |
| **False Positive** | Prusty 2013 |  | Prusty 2013 |  |
| Non-Focal |  |  |  | Pulido 2020 (thesis) |
| Non-Focal |  |  |  | Punekar 2016 |
| **Focal** |  |  |  | Qijing 2007 (translated) |
| Non-Focal |  |  |  | Qiu 2019 |
| Non-Focal |  |  |  | Qu 2020 |
| Non-Focal |  |  |  | Quigley 2014 |
| Non-Focal |  |  |  | Rachel 2012 |
| Non-Focal |  |  |  | Rahmani-Badi 2016 |
| Non-Focal |  |  |  | Rajathei 2013 |
| Non-Focal |  |  |  | Raman 2006 |
| Non-Focal |  |  |  | Ramaniuk 2018 |
| Non-Focal |  |  |  | Rankin 2002 |
| **Focal** | Reed 2021 | Reed 2021 | Reed 2021 | Reed 2021 |
| Non-Focal | Reinders 2006 | Reinders 2006 |  | Reinders 2006 |
| Non-Focal |  |  |  | Resch 2016 |
| Non-Focal |  |  |  | Rijlaarsdam 2016 |
| Non-Focal |  |  |  | Risler 2012 |
| Non-Focal |  |  |  | Rollins 2006 |
| Non-Focal | Rooney 2009 |  | Rooney 2009 | Rooney 2009 |
| Non-Focal | Rooney 2011 | Rooney 2011 | Rooney 2011 | Rooney 2011 |
| Non-Focal |  |  |  | Rosenbaum 2011 |
| Non-Focal |  |  |  | Ryan 2005 |
| Non-Focal |  |  |  | Rylander 2011 |
| Non-Focal |  |  |  | Saez 2019 |
| **Focal** | Saikatendu 2006 | Saikatendu 2006 |  | Saikatendu 2006 |
| Non-Focal |  |  |  | Sailani 2015 |
| Non-Focal | Sainsbury 2012 |  | Sainsbury 2012 | Sainsbury 2012 |
| Non-Focal |  |  |  | Samper 2019 |
| Non-Focal |  |  |  | Samudrala 2009 |
| Non-Focal |  |  |  | Sarker 2012 |
| Non-Focal |  |  |  | Schlenker 2011 (thesis) |
| Non-Focal |  |  |  | Schneeweiss 2020 |
| Non-Focal |  |  |  | Schrader 2016 |
| Non-Focal |  |  |  | Selby 2017 |
| Non-Focal |  |  |  | Selim 2021 |
| **Focal** | Sergeeva 2018 |  | Sergeeva 2018 | Sergeeva 2018 |
| Non-Focal | Shahbaaz 2013 |  | Shahbaaz 2013 | Shahbaaz 2013 |
| Non-Focal |  |  |  | Shakya 2020 (preprint) |
| Non-Focal |  |  |  | Shimada 2011 |
| Non-Focal |  |  |  | Shin 2011 |
| Non-Focal |  |  |  | Shulami 1999 |
| Non-Focal |  |  |  | Shulami 2007 |
| **Focal** | Shulami 2014 |  | Shulami 2014 | Shulami 2014 |
| Non-Focal |  |  |  | Sigdel 2011 |
| Non-Focal |  |  |  | Simon 2016 |
| Non-Focal |  |  |  | Simonis 2012 |
| Non-Focal |  |  |  | Simpson 2015 |
| Non-Focal |  |  |  | Skottman 2005 |
| Non-Focal |  |  |  | Skunca 2013 |
| Non-Focal | Spinola 2010 |  | Spinola 2010 | Spinola 2010 |
| Non-Focal | Stabler 2010 |  |  | Stabler 2010 |
| Non-Focal |  |  |  | Starkey 2009 |
| Non-Focal |  |  |  | Stenger 2019 (thesis) |
| Non-Focal |  |  |  | Stern 2020 |
| Non-Focal |  |  |  | Steyn 2017 (thesis) |
| Non-Focal |  |  |  | Strillacci 2020 |
| Non-Focal |  |  |  | Su 2012 |
| Non-Focal |  |  |  | Suzuki 2006 |
| Non-Focal |  |  |  | Ta 2010 |
| Non-Focal |  |  |  | Tajkarimi 2017 |
| **Focal** | Tascou 2000 | Tascou 2000 |  | Tascou 2000 |
| **Focal** | Tascou 2003 | Tascou 2003 |  | Tascou 2003 |
| Non-Focal |  |  |  | Thankam 2018 |
| Non-Focal |  |  |  | Thorenoor 2010 |
| Non-Focal |  |  |  | Tian 2014 |
| Non-Focal |  |  |  | Tian 2020 |
| **Focal** | Tomoike 2009 |  | Tomoike 2009 | Tomoike 2009 |
| Non-Focal |  |  |  | Tran 2018 |
| Non-Focal |  |  |  | Tsuiko 2016 |
| Non-Focal |  |  |  | Turlin 2006 |
| Non-Focal |  |  |  | Uxa 2019 |
| Non-Focal | van Diemen 2017 | van Diemen 2017 |  | van Diemen 2017 |
| Non-Focal | Van Dyke 2016 |  | Van Dyke 2016 | Van Dyke 2016 |
| Non-Focal |  |  |  | van Ouwerkerk 2019 |
| Non-Focal |  |  |  | Volha 2019 (thesis) |
| Non-Focal |  |  | Wagley 2014 | Wagley 2014 |
| Non-Focal |  |  |  | Waldor 2013 |
| Non-Focal |  |  |  | Wang 2010 |
| **False Positive** | Wang 2012 | Wang 2012 |  | Wang 2012 |
| Non-Focal |  |  |  | Wang 2015 |
| Non-Focal |  |  |  | Wang 2018 |
| Non-Focal |  |  |  | Wang 2021 |
| Non-Focal |  |  |  | Watkins 2008 |
| Non-Focal |  |  |  | Watkins 2010 |
| Non-Focal |  |  |  | Wei 2001 |
| Non-Focal | Wei 2016 | Wei 2016 |  | Wei 2016 |
| Non-Focal |  |  |  | Weinzierl 2008 |
| Non-Focal |  |  |  | Wilkinson 2018 (thesis) |
| Non-Focal | Williams 2013 | Williams 2013 |  | Williams 2013 |
| Non-Focal |  |  |  | Willis 2010 (thesis) |
| Non-Focal |  |  |  | Winer 2011 |
| Non-Focal |  |  |  | Wittchen 2018 |
| Non-Focal |  |  |  | Wu 2019 |
| Non-Focal |  |  |  | Xi 2011 |
| Non-Focal |  |  |  | Xia 2017 |
| Non-Focal |  |  |  | Xiang 2012 |
| **Focal** |  |  |  | Xie 2019 |
| Non-Focal | Xu 2021 | Xu 2021 |  |  |
| Non-Focal |  |  |  | Yan 2014 |
| Non-Focal |  |  |  | Yan 2020 |
| Non-Focal |  |  |  | Yang 2013 |
| Non-Focal |  |  |  | Yao 2016 |
| **Focal** |  |  |  | Yu 2015 |
| Non-Focal |  |  |  | Yu 2018 |
| Non-Focal |  |  |  | Yuan 2006 |
| Non-Focal |  |  |  | Zamanian-Daryoush 2020 |
| Non-Focal |  |  |  | Zhao 2018 |
| Non-Focal |  |  |  | Zheng 2020 |
| Non-Focal |  |  |  | Zuccotti 2008 |

### Supplementary Elaborations

**Elaboration S1.** Explanation of XynX “productive ignorance” example.

XynX, a DUF34 homolog of *Geobacillus stearothermophilus* T-6, was identified and described by a couple researcher groups over the course of several papers between the years 1994 and 2016 [1–6]. Upon identification, because of its genomic location and position within an operon of biosynthetic interest, it was named with respect to neighboring genes. This naming and system-level characterization were completed almost entirely independent of the preceding literature of the protein family of which it was a recognizable member. Additionally, and although the genomic information of the organism had been purportedly annotated and respective data made publicly available, it was—upon review of Reed et al. *Biomolecules* 2021 [7]—discovered that the genome and related annotation information was not as accessible as had been claimed. It had been observed that the complete genome and any annotated, encoded proteins had been intermittently submitted and updated by the submitter over the course of 15 years (1993-2008). Although the EuropePMC Data subsection for the earliest publication [1] lists 51 links to individual UniProt entries and a single ENA link to 57 nucleotide records (accessed May 20, 2022; <https://europepmc.org/article/MED/8031084>): the original publisher does not provide links to this information (now hosted at the website for the parent journal, ASM; <https://doi.org/10.1128/aem.60.6.1889-1896.1994>); this paper does not provide any systematic accession identifiers for the data allegedly being reported (i.e., the genome); and only 56 of the 57 EuropePMC reported genes are actually listed in the ENA database under the queryable publication-associated ID, 8031084.

It is hypothesized that this EuropePMC data being reported as being associated with this publication [1] is mistakenly attributed to this particular record. At this time, it is suspected that the accessioned data referenced in many of these online publications has been retroactively linked to these other works from the Gat et al. 1994 record and, subsequently, the Shulami et al. 2011 publication in the Journal of Bacteriology [5] (<https://doi.org/10.1128/JB.00222-11>; first cited by Alalouf et al. 2011 in the Journal of Biological Chemistry (PMID: 21994937) [4], methods section) with which a GenBank record was submitted for the 77,747 bp (linear) of the *G. stearothermophilus* T-6 genome that was reported to contain “the xylan, xylose, arabinan, and arabinose utilization region” relevant to the study. The record, GenBank: DQ868502.2, is titled “Geobacillus stearothermophilus strain T-6 genomic sequence”, suggesting that the genome is completely sequenced; GenBank record is dated “26-JUL-2016”. This record replaced the earlier GenBank record, DQ868502.1, titled “Geobacillus stearothermophilus strain T-6 xylan utilization gene cluster, complete sequence; and NAD(P)H-dependent flavin oxidoreductase (orfD) gene, partial cds” dated 01-FEB-2007. As per Gat et al., 1994, the earliest genomic record (annotated with only a few CDSs, but largely unannotated) was submitted to GenBank, DDBJ, and EMBL (ENA) with the accession identifier Z29080. The first in-text description of *xynX* and its encoded protein of the same name was in Shulami et al., *Applied and Environmental Microbiology* 2007 [3] and described as a “xylan utilization gene”. In Alalouf et al., 2011 [4], XynX is described as “a putative regulatory gene with unknown function”. It’s putative cooperative regulation of *xynA* expression alongside XylR and CodY is described in more detail in Shulami et al., *Journal of Biological Chemistry* 2014 [5]. No description of the family to which XynX belongs is mentioned, no attempt to identify the associated protein family is made, despite assigning the protein a name and describing a functional association. Although these characterizations (and alias) were assigned with ignorance of the protein family, the functional association of XynX as a DUF34 member was not dissimilar from subsequent characterizations, for example, in *Bacillus subtillis* [8], and, therefore, cannot be discounted in its contributions to describing functional associations at the family level.

### Supplementary Figures

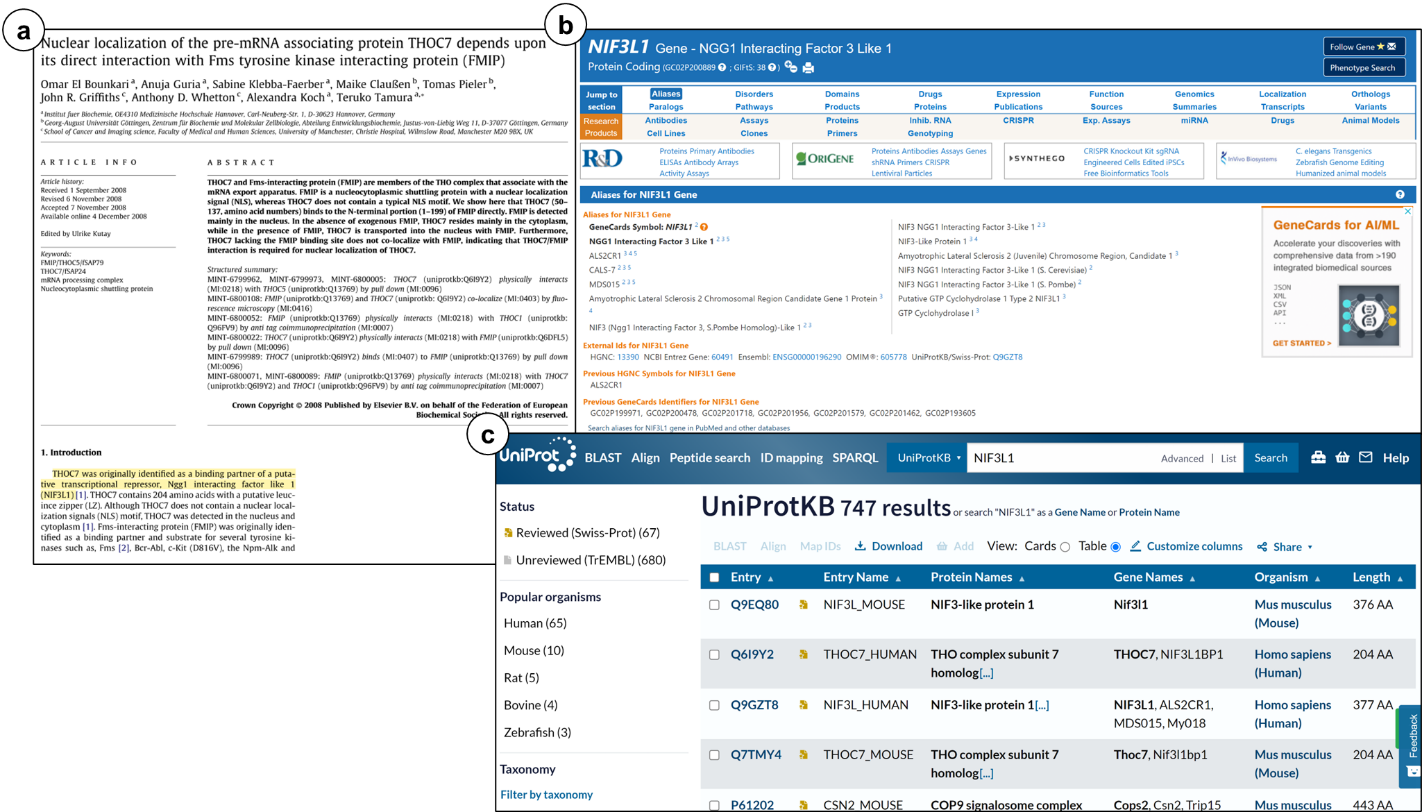
**Figure S1.** Seed Information Assessment: establish understanding of starting information in preparation for the capture of published data.

**Figure S1. Seed Information Assessment.** Assess what information you already have/know about your protein and its family. Do you have only text (e.g., aliases) (a-b) or (c) do you have access to tentatively representative member sequences? Both (a-c) and, if so, how many? Do you suspect that they are adequately representative of the family?

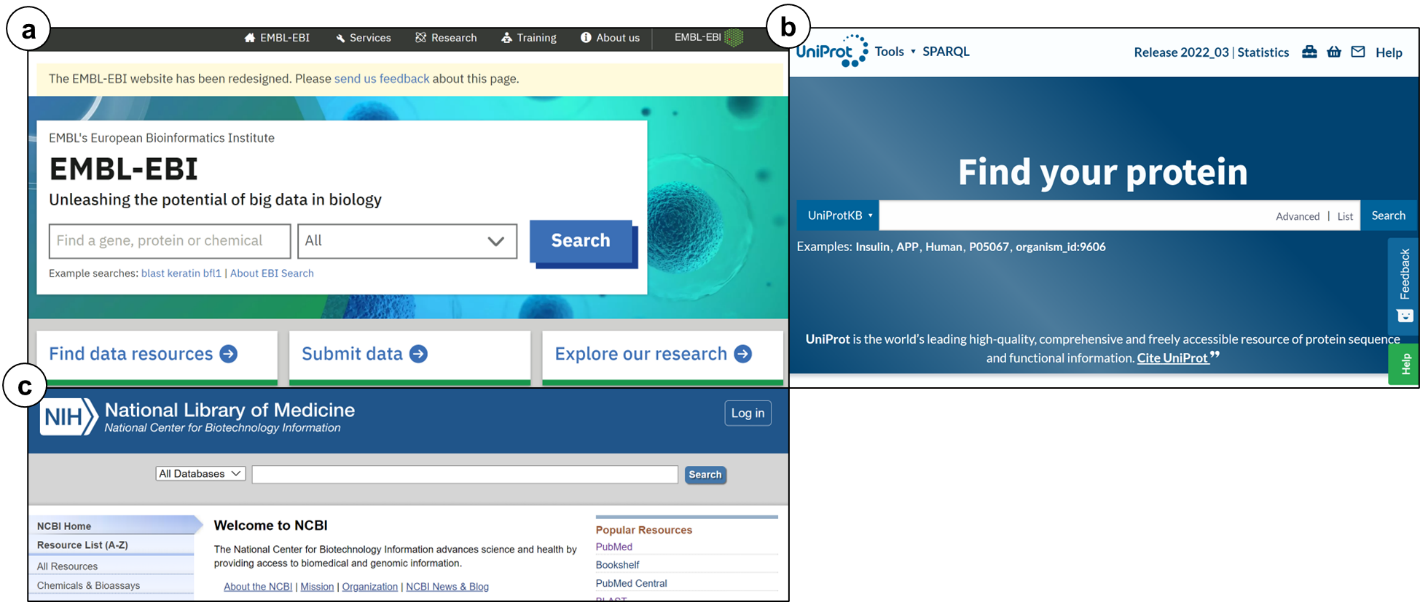
**Figure S2.** Retrieve sequences using known names/aliases.

**Figure S2. Retrieve Sequences.** Given a lack of member sequences but several known aliases, text searches for verifiable tentative representative sequences are recommended. A number of specialized search engines exist and are not limited to sequence databases (e.g., annotation or ortholog databases). Here, examples include (a) EMBL-EBI broad search, (b) UniProtKB, and (c) NIH National Library of Medicine all-database search.

**Figure S3a.** Family-level analysis examples from Annotree (left) and EFI Taxonomy (right).

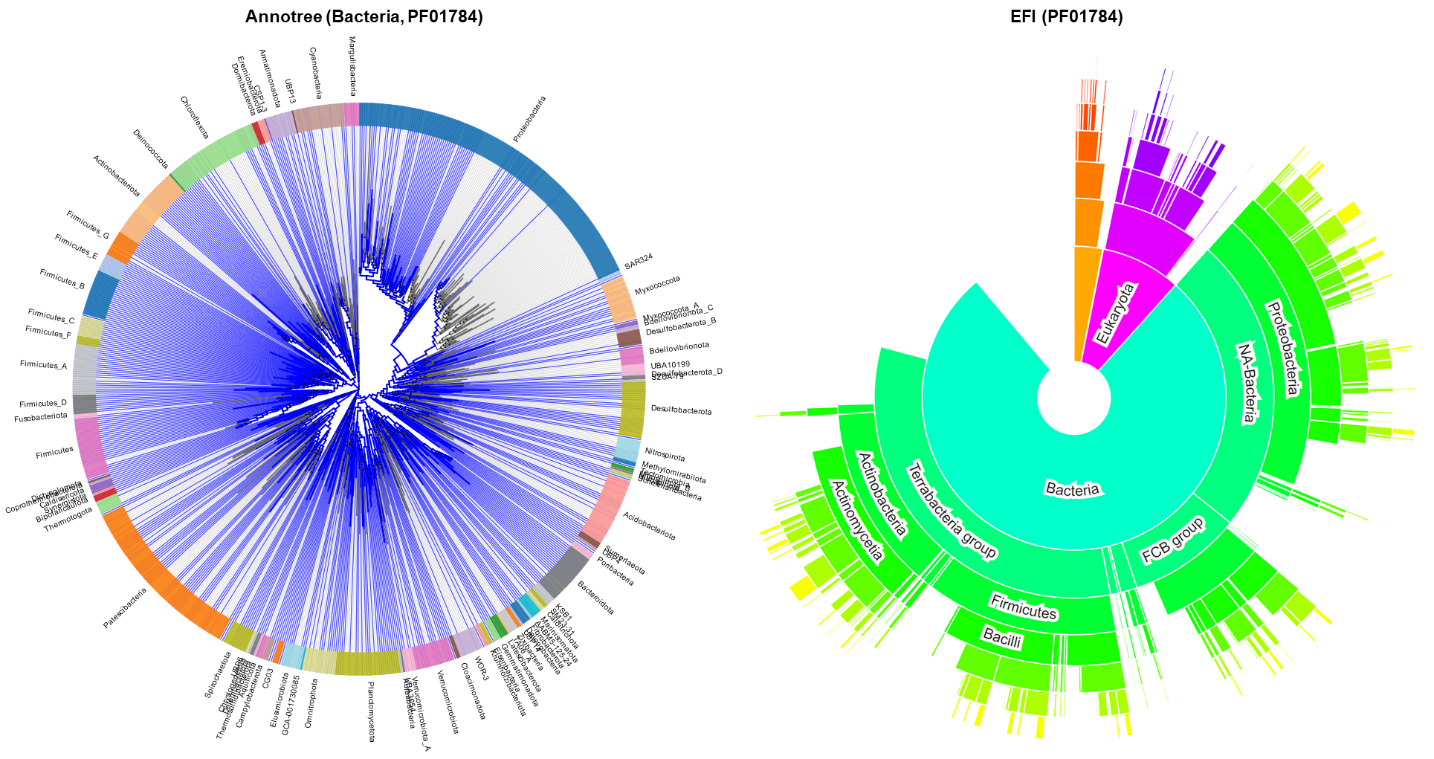

**Figure S3b.** Family-level analyses using various protein family classification databases (i.e., Pfam, CDD, OrthoMCL, InterPro, OrthoInspector, EggNOG, ENSEMBL tools), of which can be used to approximate architectural subgroups in concert with using representative sequences as inputs into the sequence-based search tools (e.g., i.e., PaperBLAST) for corpus retrieval.

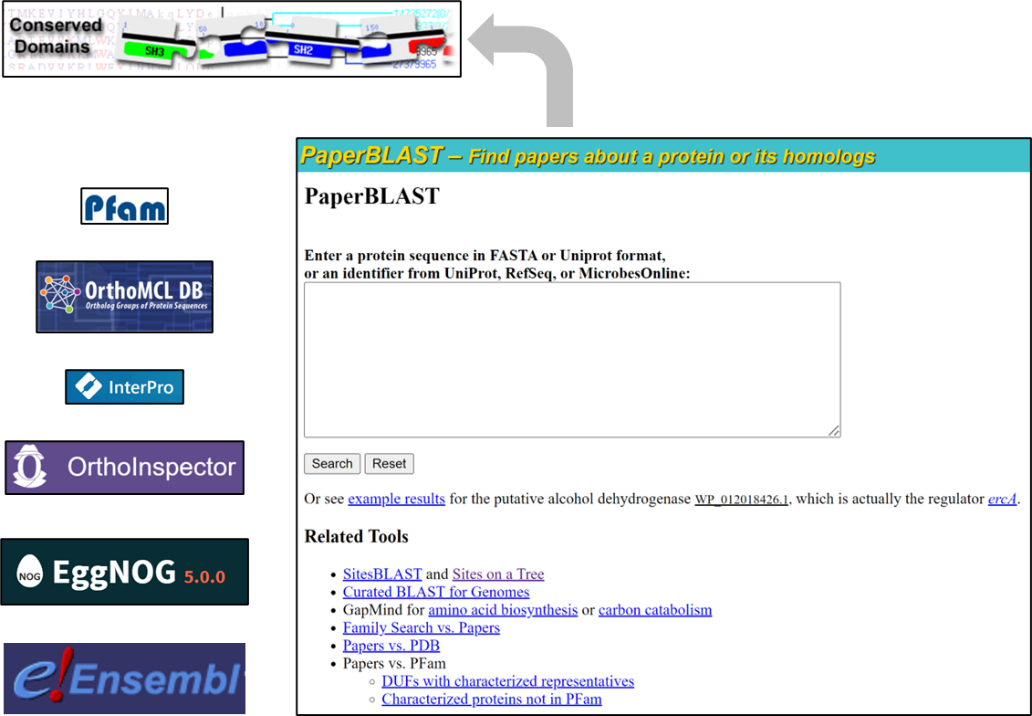

**Figure S4.** Example of over-propagation of root family attribution in EggNOG Database as a result of fusions. EggNOG aggregates data from other resources and often these collections accrue family members that are domain architectural variants and contain non-family sequence annotations. These annotations are not filtered out or flagged in any way and require the user to discern these erroneous aggregates on their own. (a) EggNOG root cluster of the DUF34 protein family. Added notation of screenshot: red bars underline the incorrect annotations aggregated from KEGG, one biologically incorrect based on published data, a second resulting from a fusion sequence of COG0327 and COG1579 [PMID: 34572495], and a third that is a product of a gene fusion of a eukaryotic DUF34 homolog and CIAO (COG2319). (b) Link-out to the first of three incorrect KO entries listed EggNOG listed KO, K22391, for the enzyme activity of GTP cyclohydrolase I; KO entry lists the corresponding COG, though in this case, no additional COGs are aggregated in KEGG Orthology. (c) Link-out to the second of three listed incorrect KOs of DUF34, that for an uncharacterized bacterial protein family, COG1579 (K07164), an annotation recognized as being associated with the DUF34 family by EggNOG—an association that is likely through the known fusion with the *Wollinella succinogenes* DUF34 homolog [PMID: 34572495] that may or may not have functionally diverged—while KO does not identify the source of this association. (d) Link-out to the third of three incorrect KOs listed in the entry card for LCOG0327 in EggNOG, CIAO (COG2319), also a product of fusion sequences within the DUF34 family. (e) COG0327 (child of root, LCOG0327) “Duplications profile” view.

**a.**

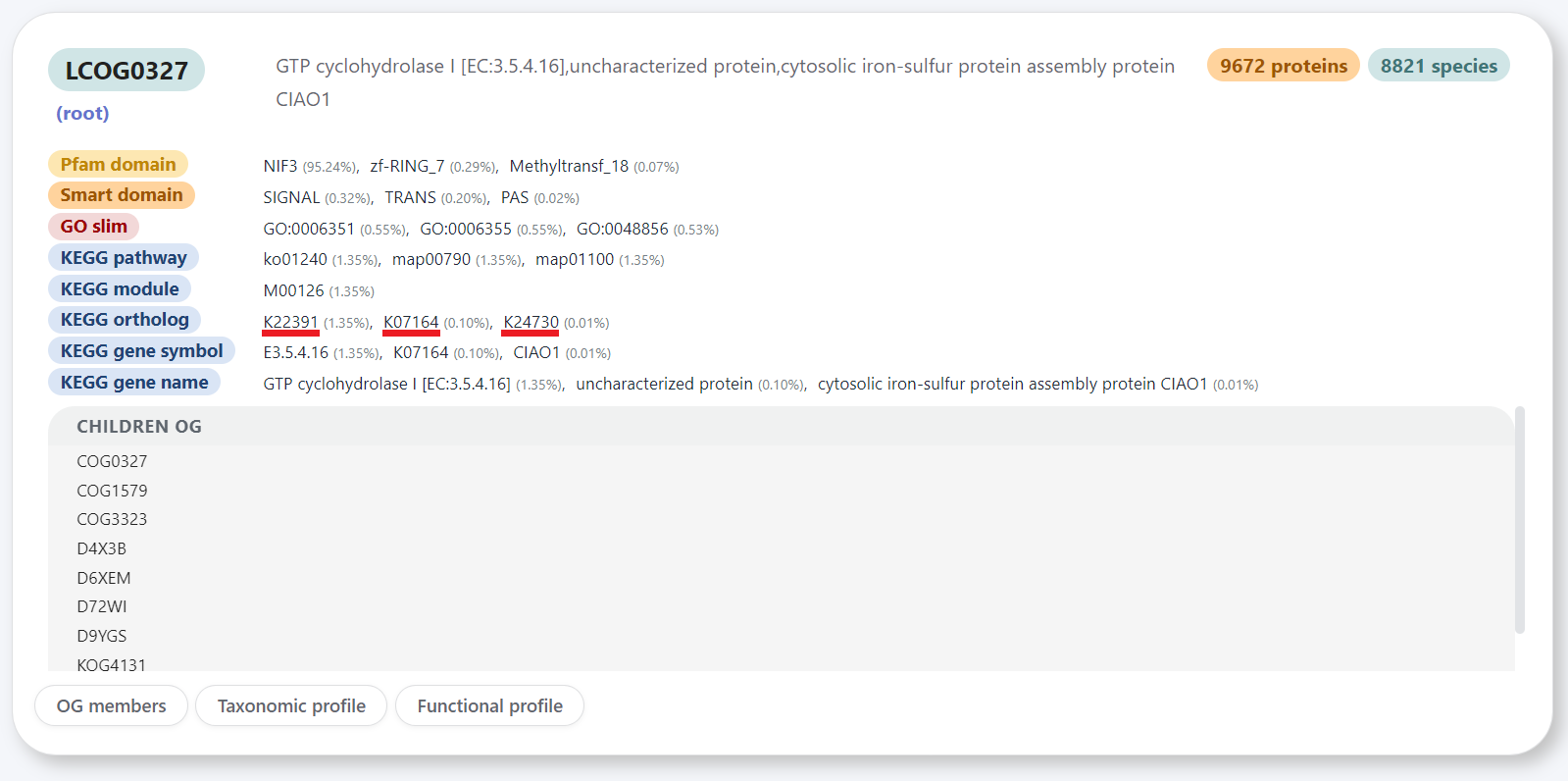

**b.**

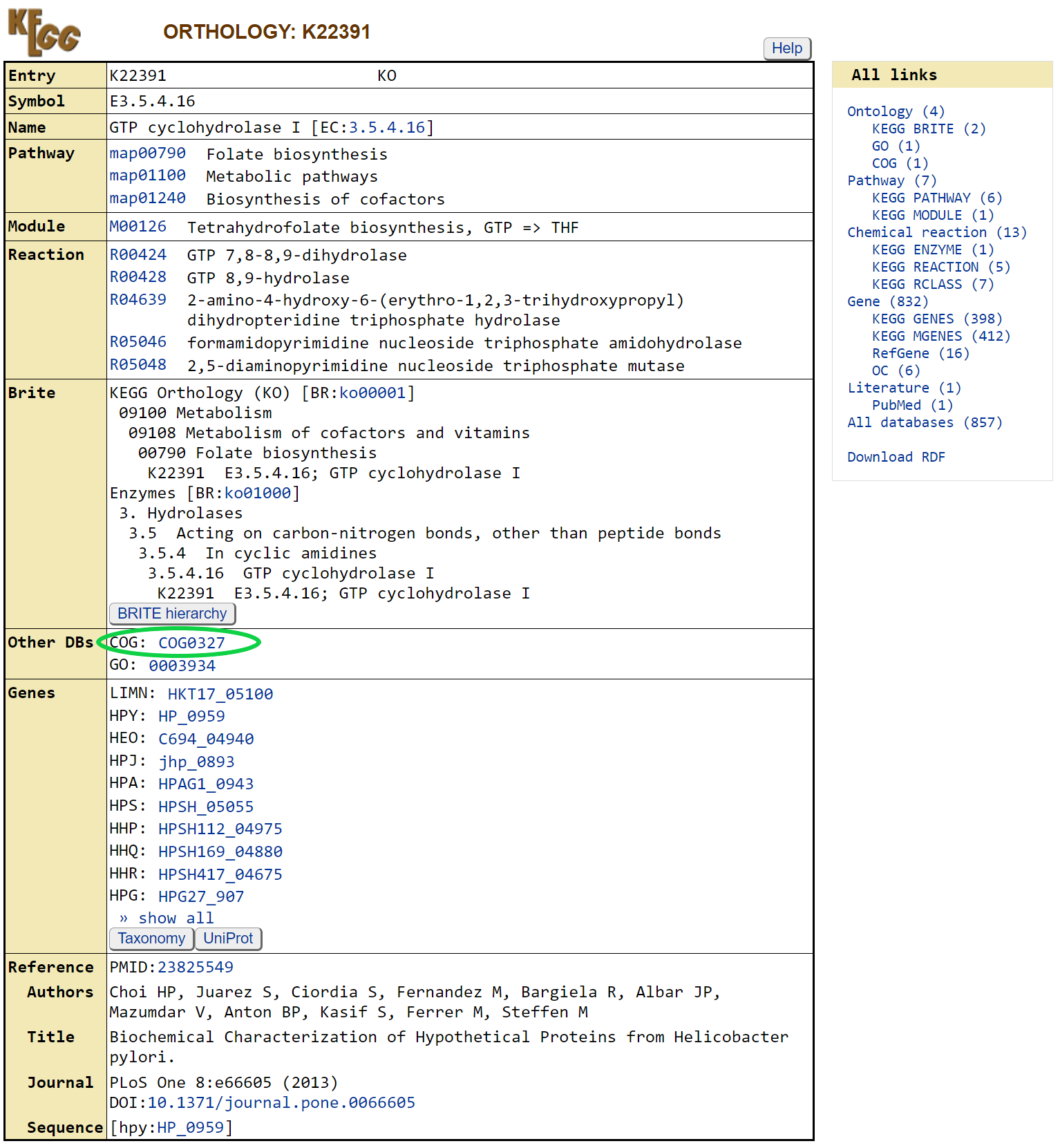

**c.**

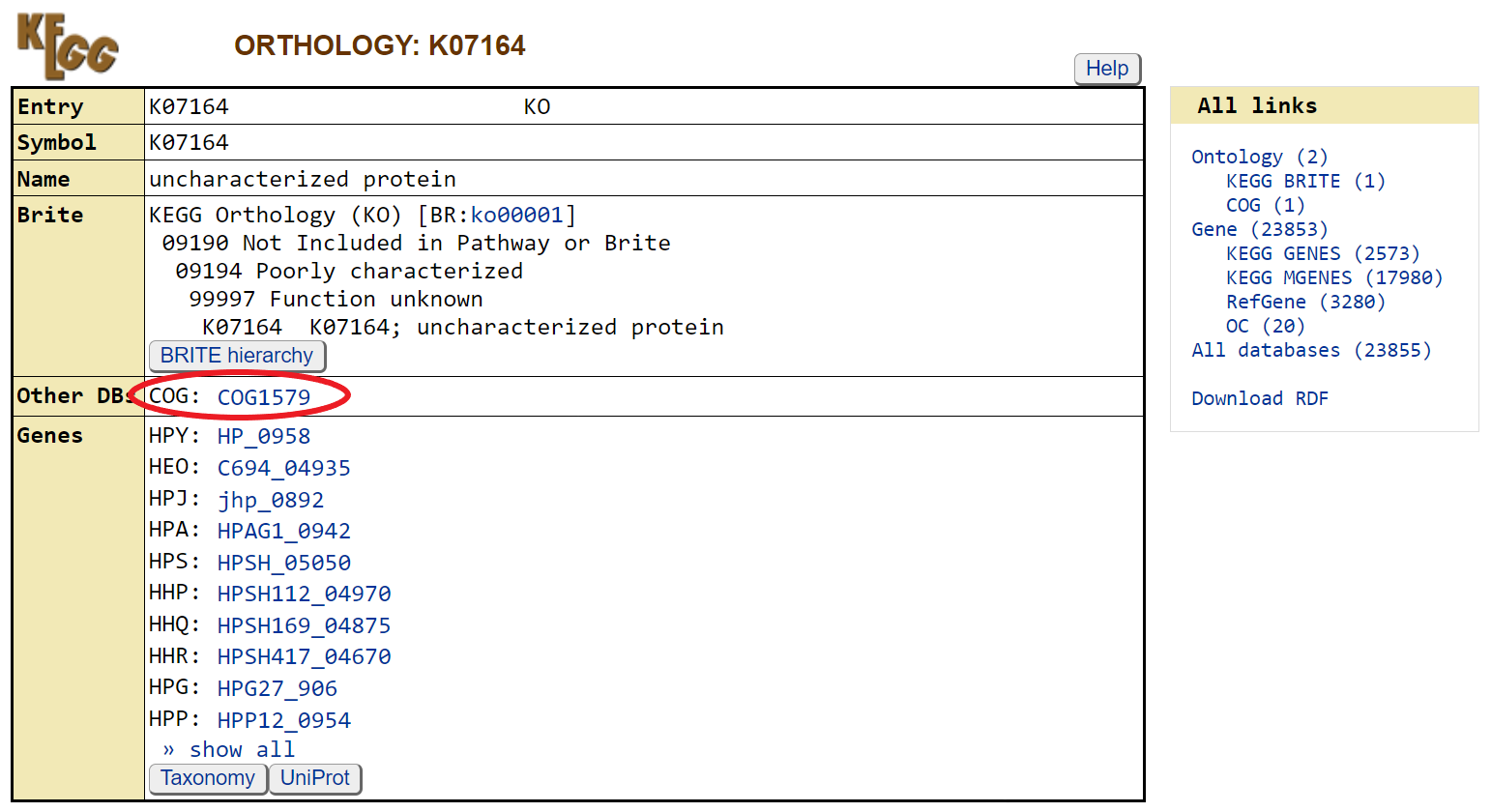

**d.**

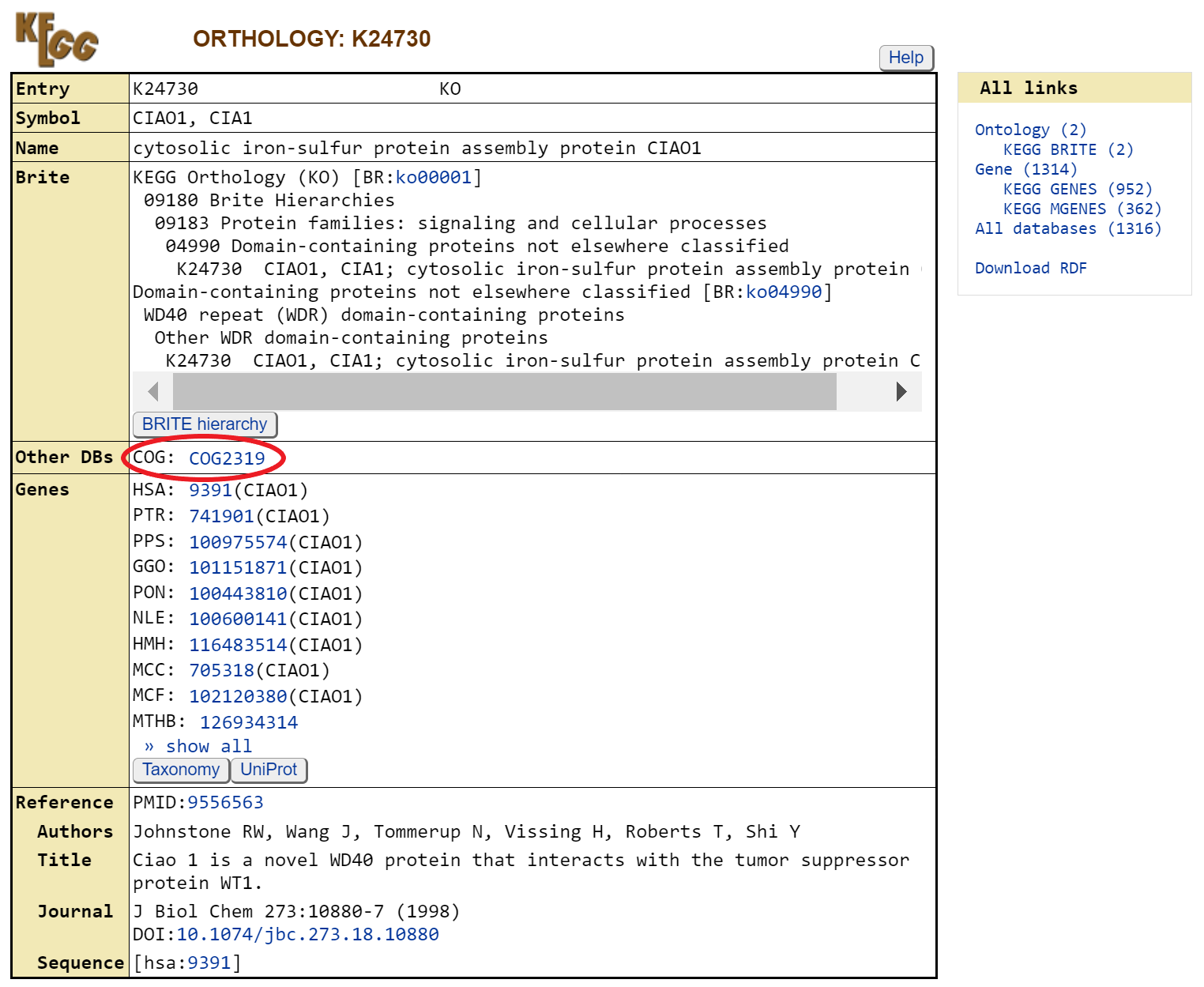

**e.**

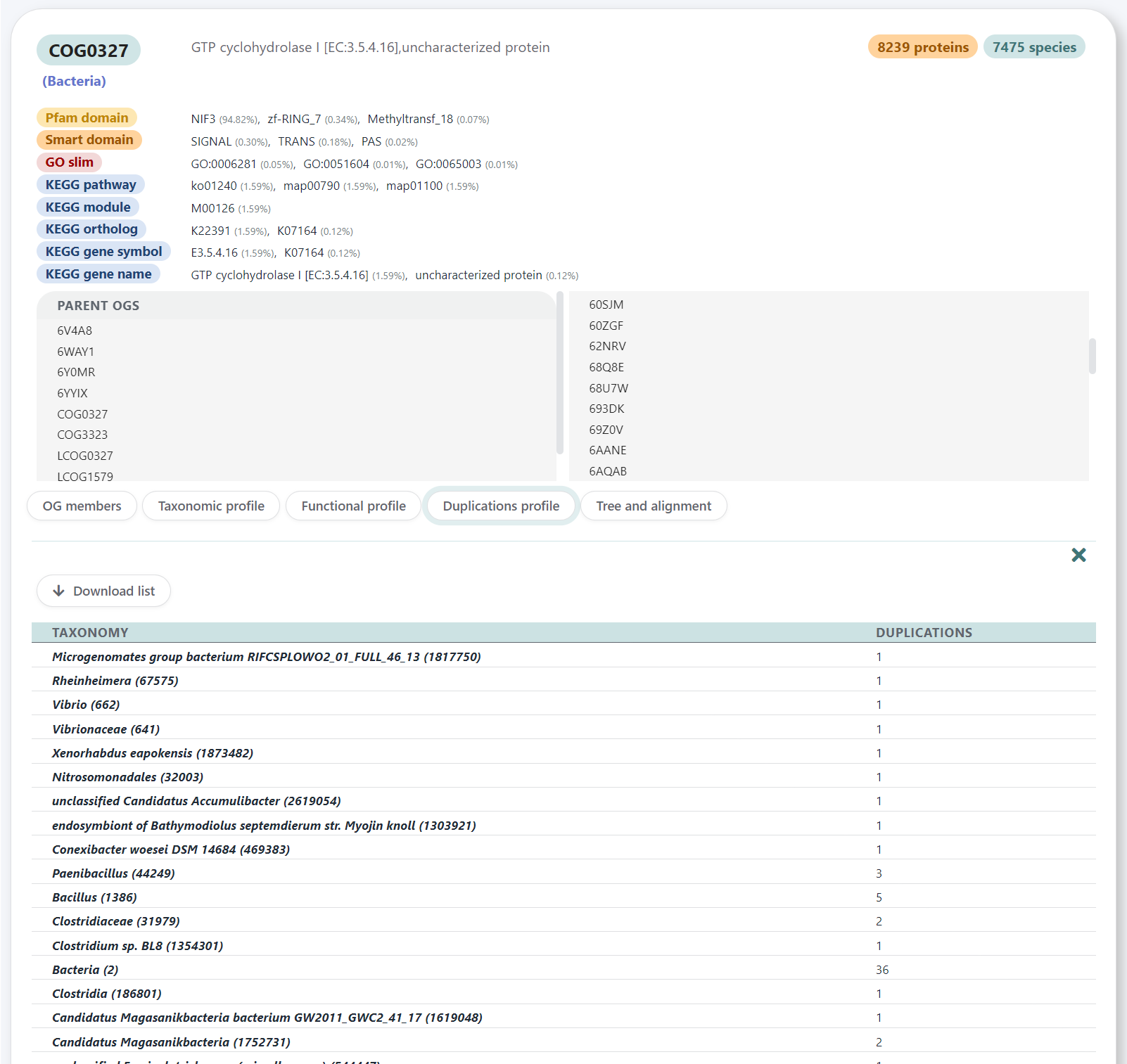

**Figure S5.** WorldWideScience.org-, ScienceResearch.com-generated diagrams of keywords. Visualized yields illustrate coincidental homonyms of NIF3 (DUF34).

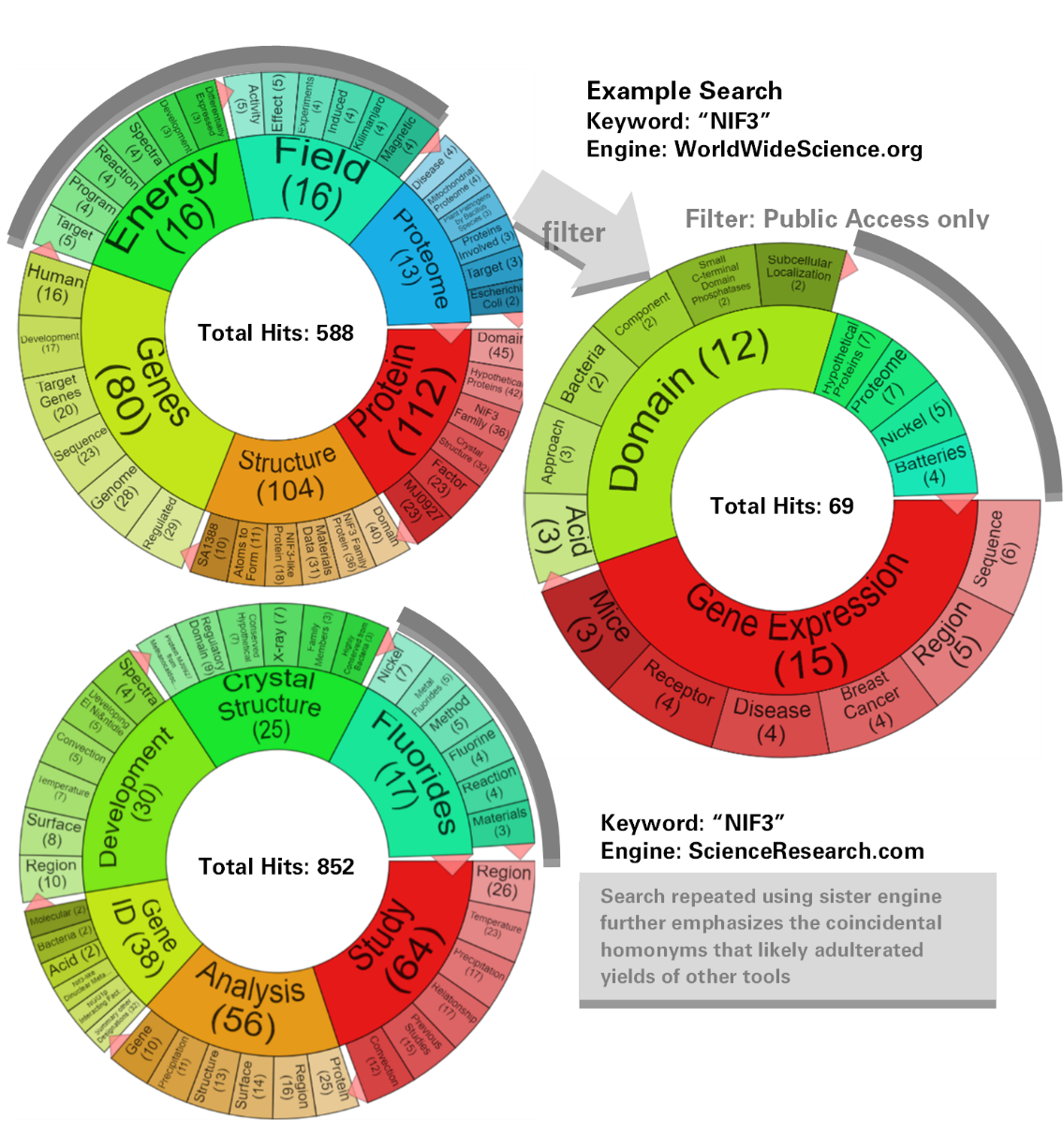

**Figure S6.** Coincidental homonyms are challenges for text-based search engines, even those as specialized as PubTator. Top screenshot shows PubTator top hits for “GTP cyclohydrolase 1 type 2” search, while the bottom screenshot shows the top hits for alias synonym “GTP cyclohydrolase I type 2”. These searches should hypothetically retrieve identical results; however, in addition to the false synonyms being retrieved due to the automatic generalization of the original search term to “GTP cyclohydrolase”, the engine simultaneously retrieves a correct result in only one of the two case uses (the bottom instance, “GTP cyclohydrolase I type 2”), suggesting that the tool is unable to discern singular “I” usage as the Roman numeral for 1, which is nomenclature practice frequently used in the naming of genes/proteins.

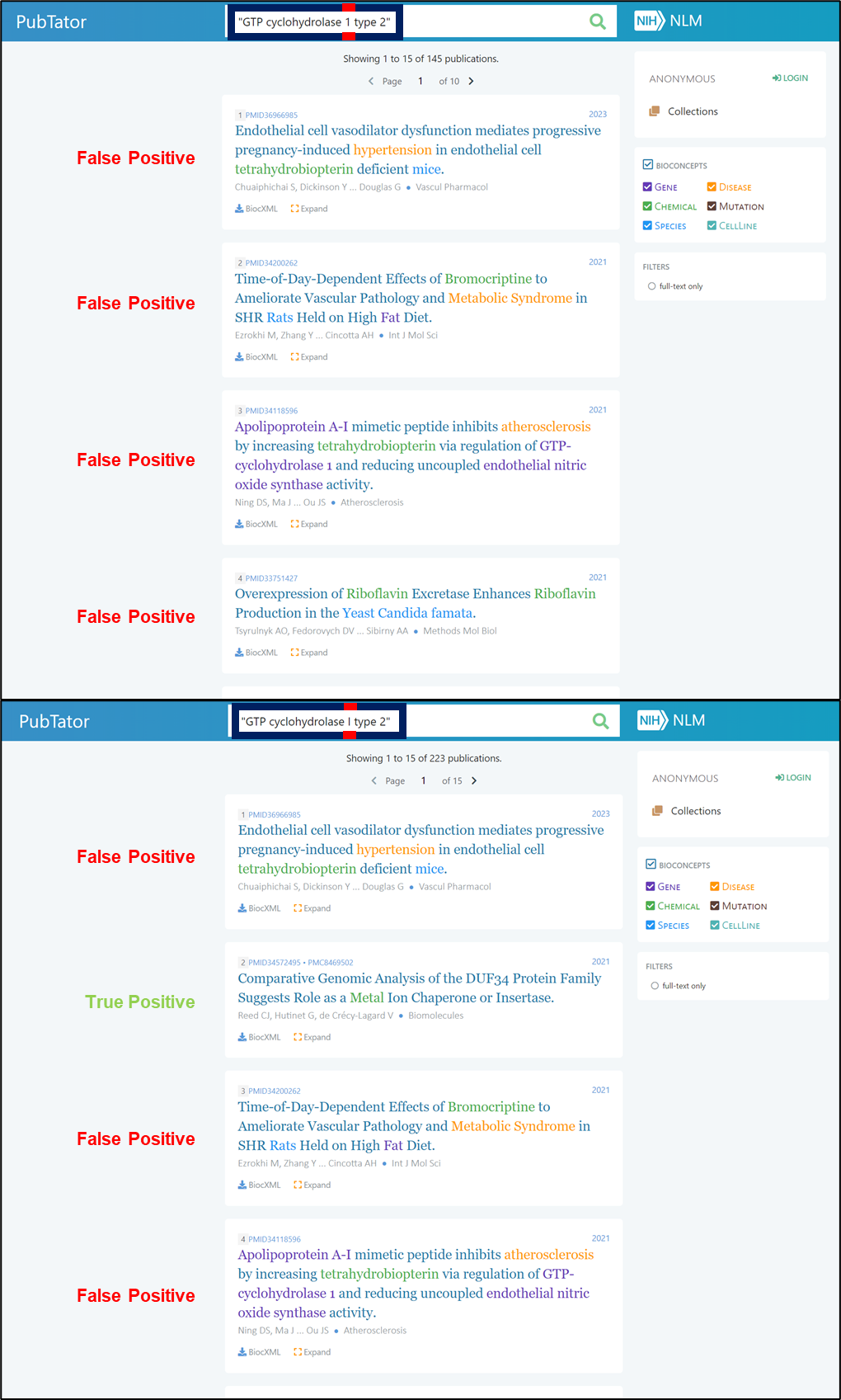

**Figure S7.** FlaGs (WebFlaGs) example outputs for NCBI BLASTp results for COG0327 (DUF34) family member of *E. coli*, YbgI (P0AFP6); list of RefSeq Accession IDs was used as the WebFlaGs input. a) Shows the PDF output showing the gene neighborhoods of the input sequences; cluster information is also available in text format accompanying this graphical output (not shown). b) Flanking genes tree graphical result option 1. c) Flanking genes tree graphical result option 2. d) MAFFT sequence alignment tree graphical result. The latter panels, b-d, are available as components constituting the exportable zipped folder available upon FlaGs output receipt.

**a.**

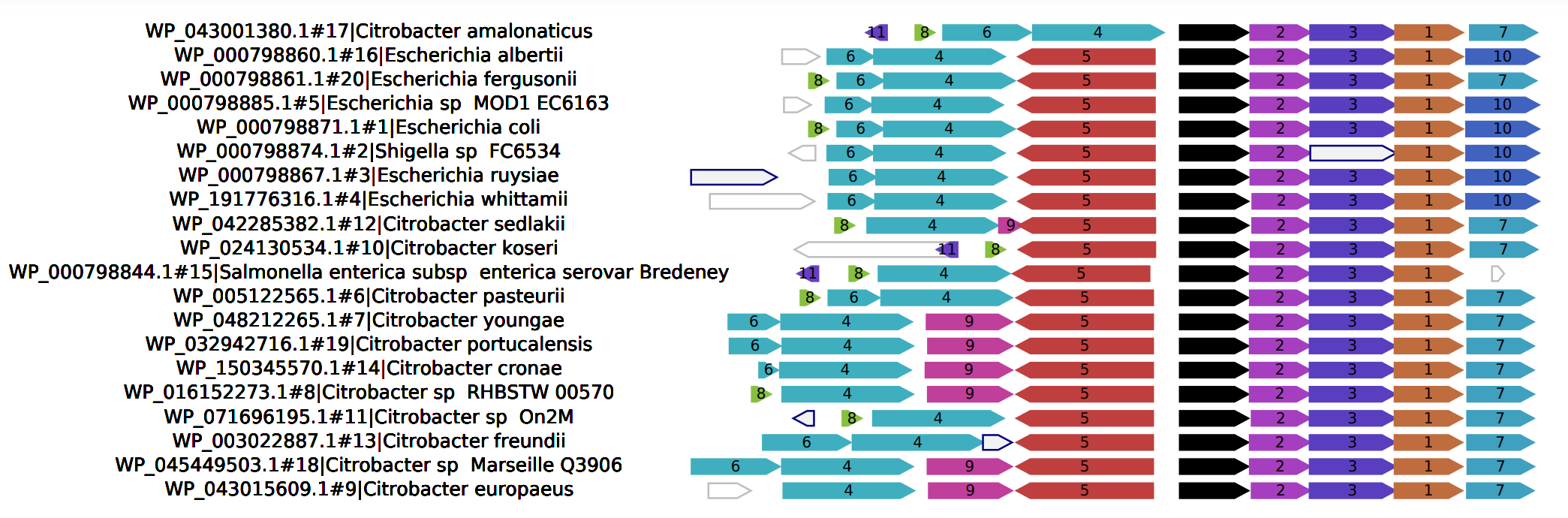

**b.**

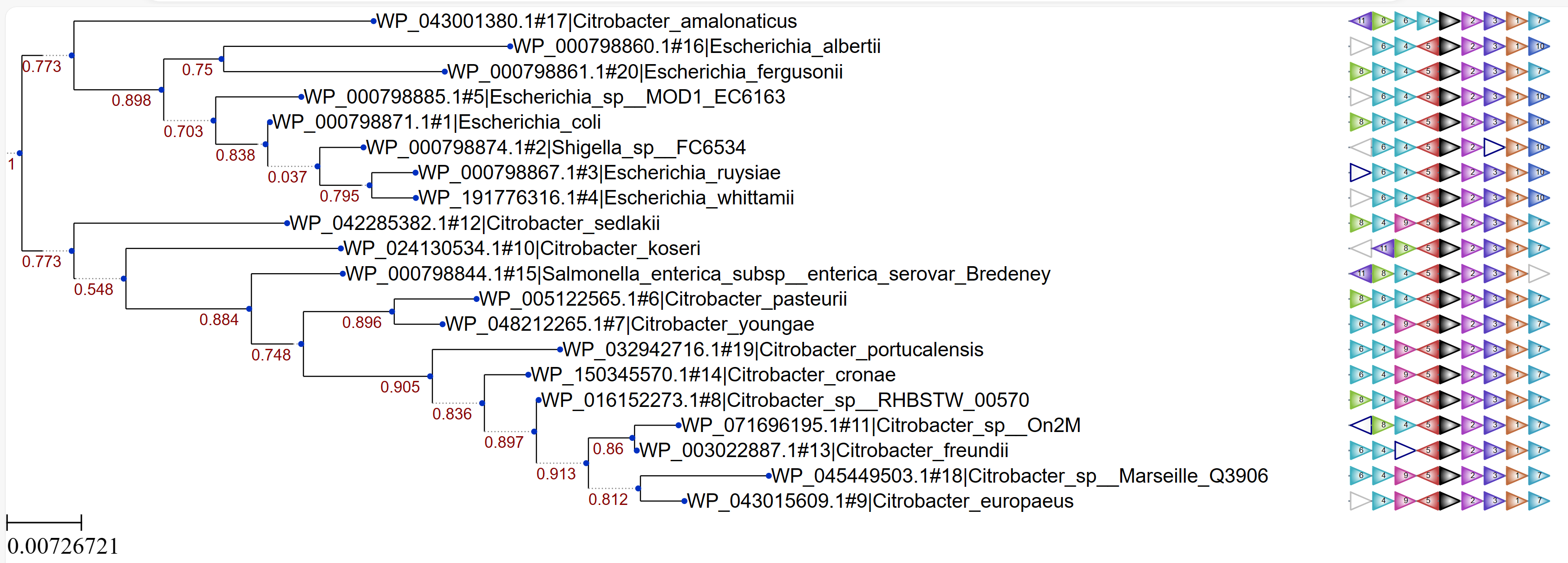

**c.**

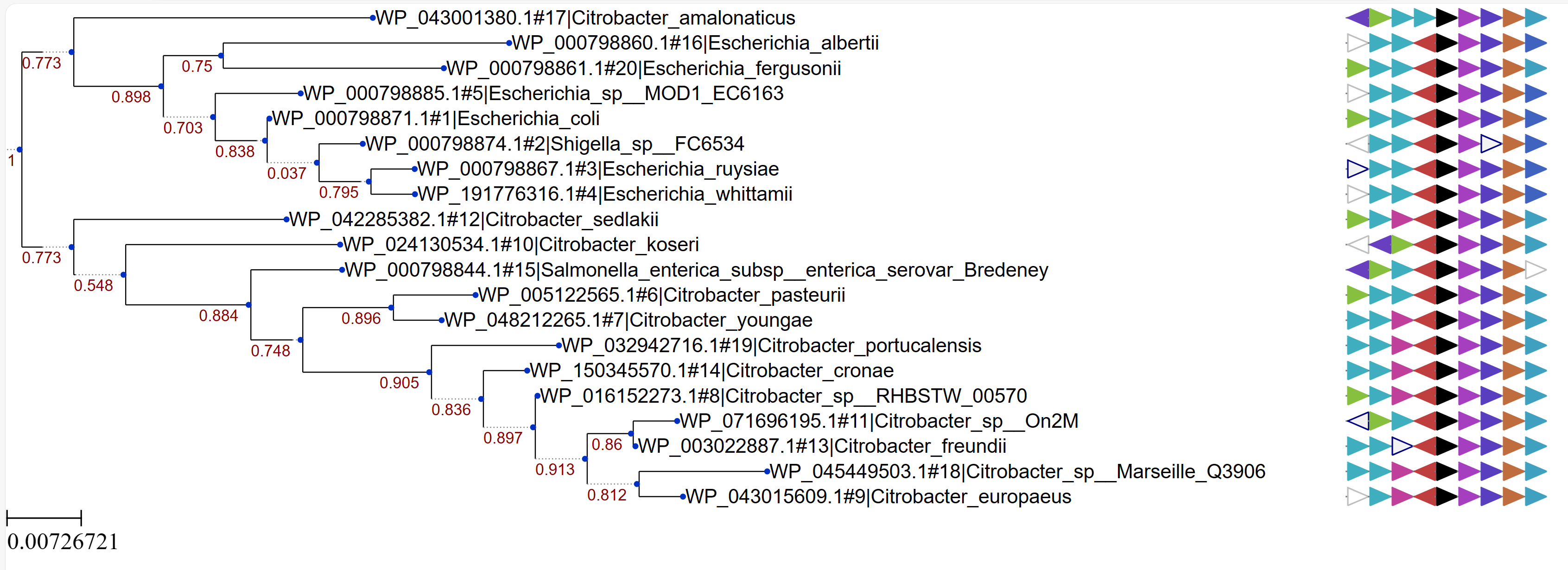

**d.**

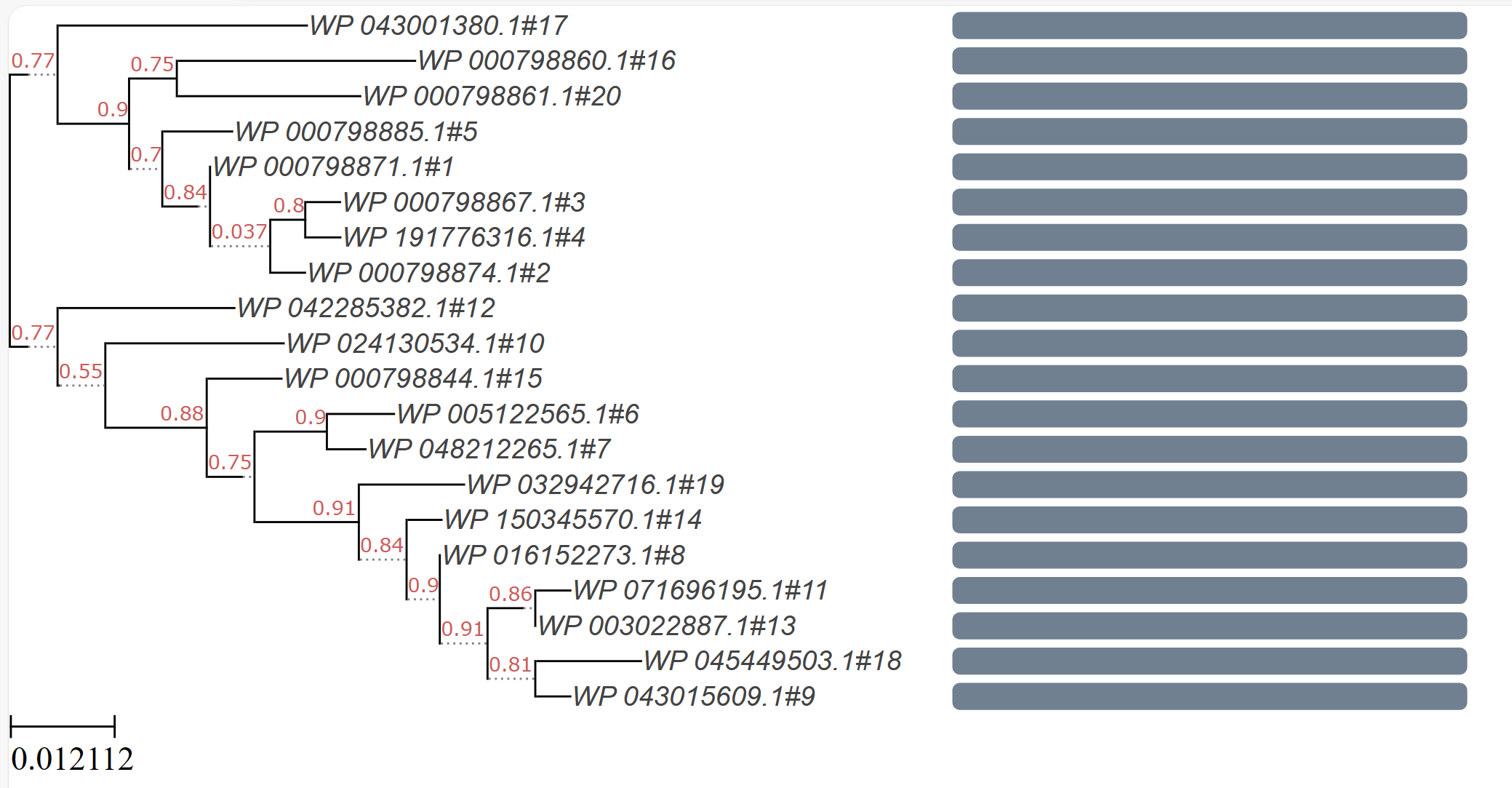

**Figure S8.** EFI-GNT

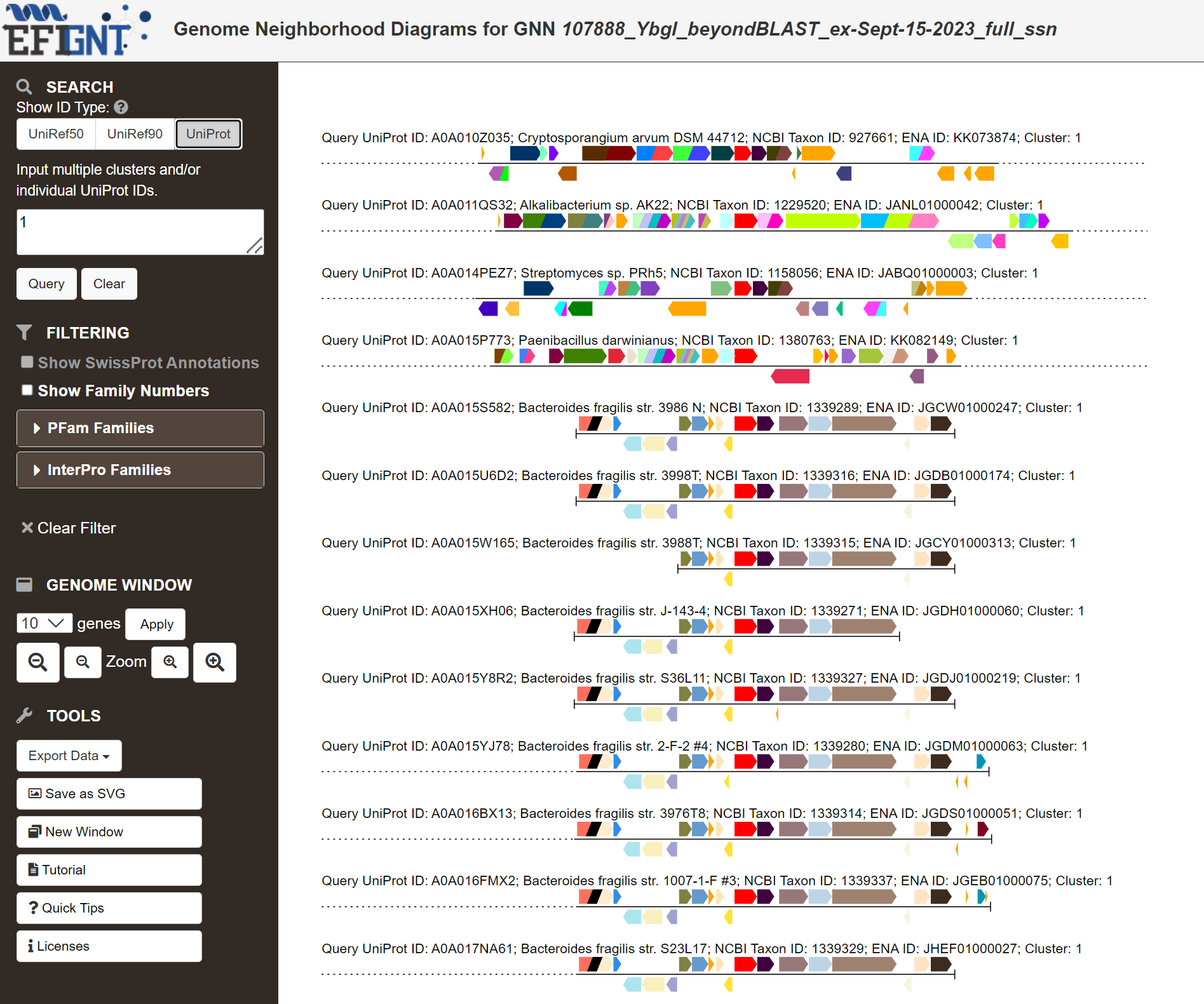

**Figure S9.** Annotree example: single family, PF02591, in bacteria.

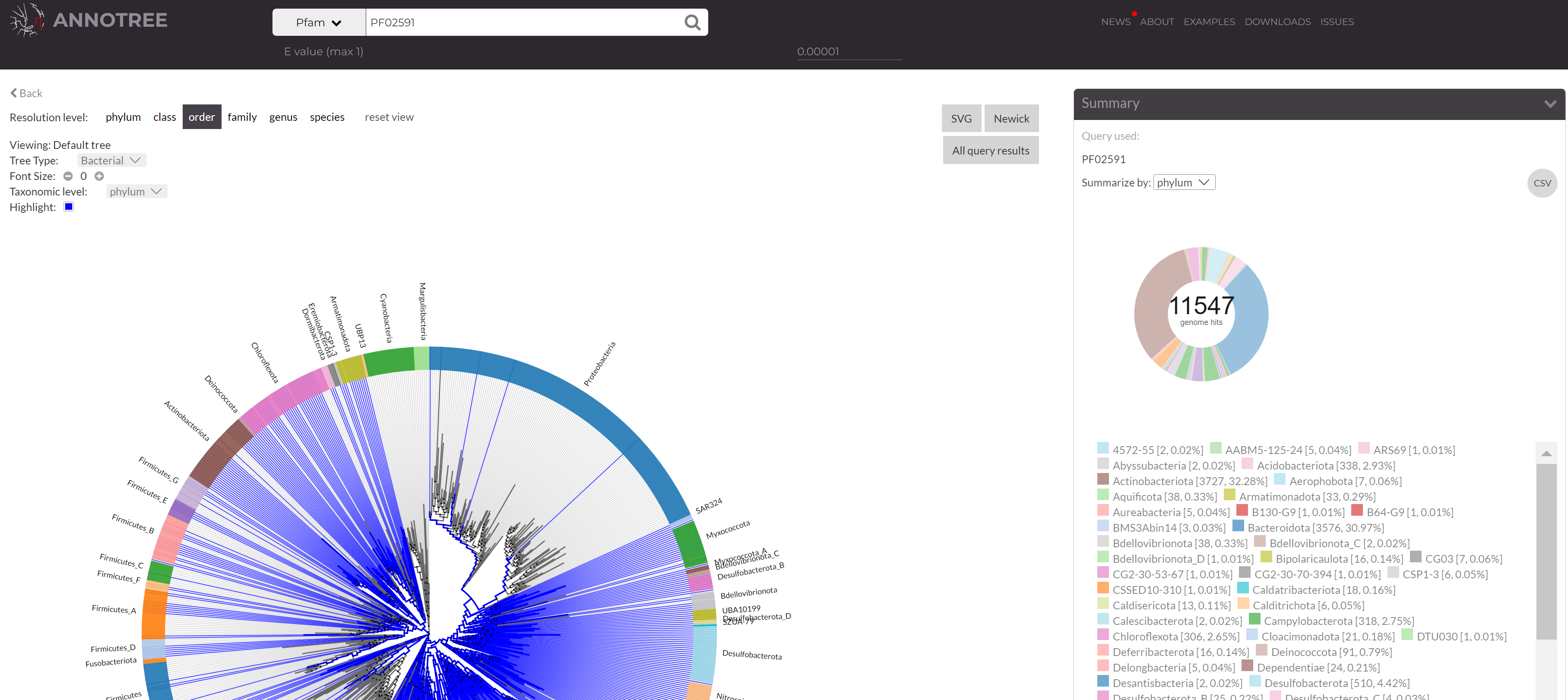

**Figure S10.** COGNAT COG-based gene neighborhood and physical clustering tool example output for COG0327 (DUF34 protein family). a) shows the raw output, while b) shows a portion of the pdf export made available via link-out download.

**a.**

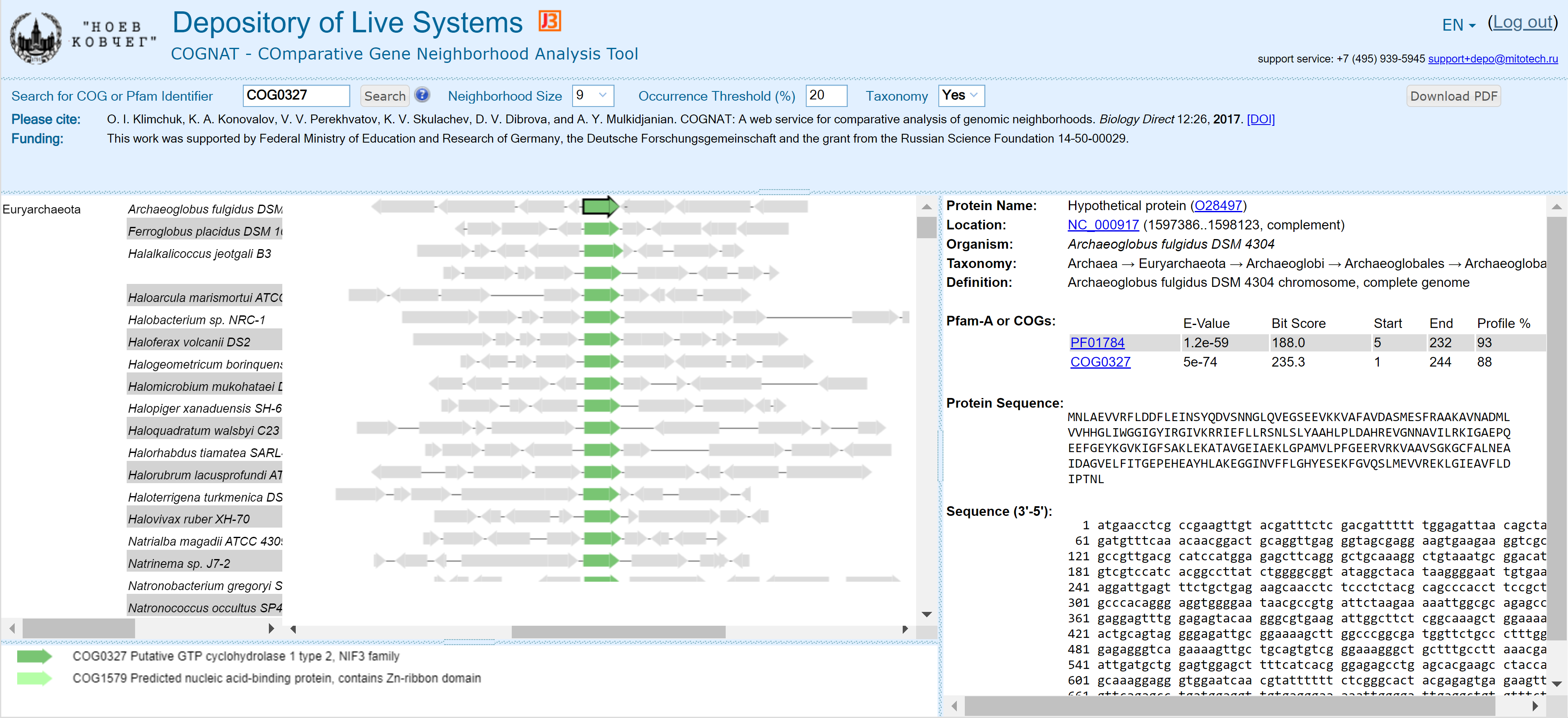

**b.**

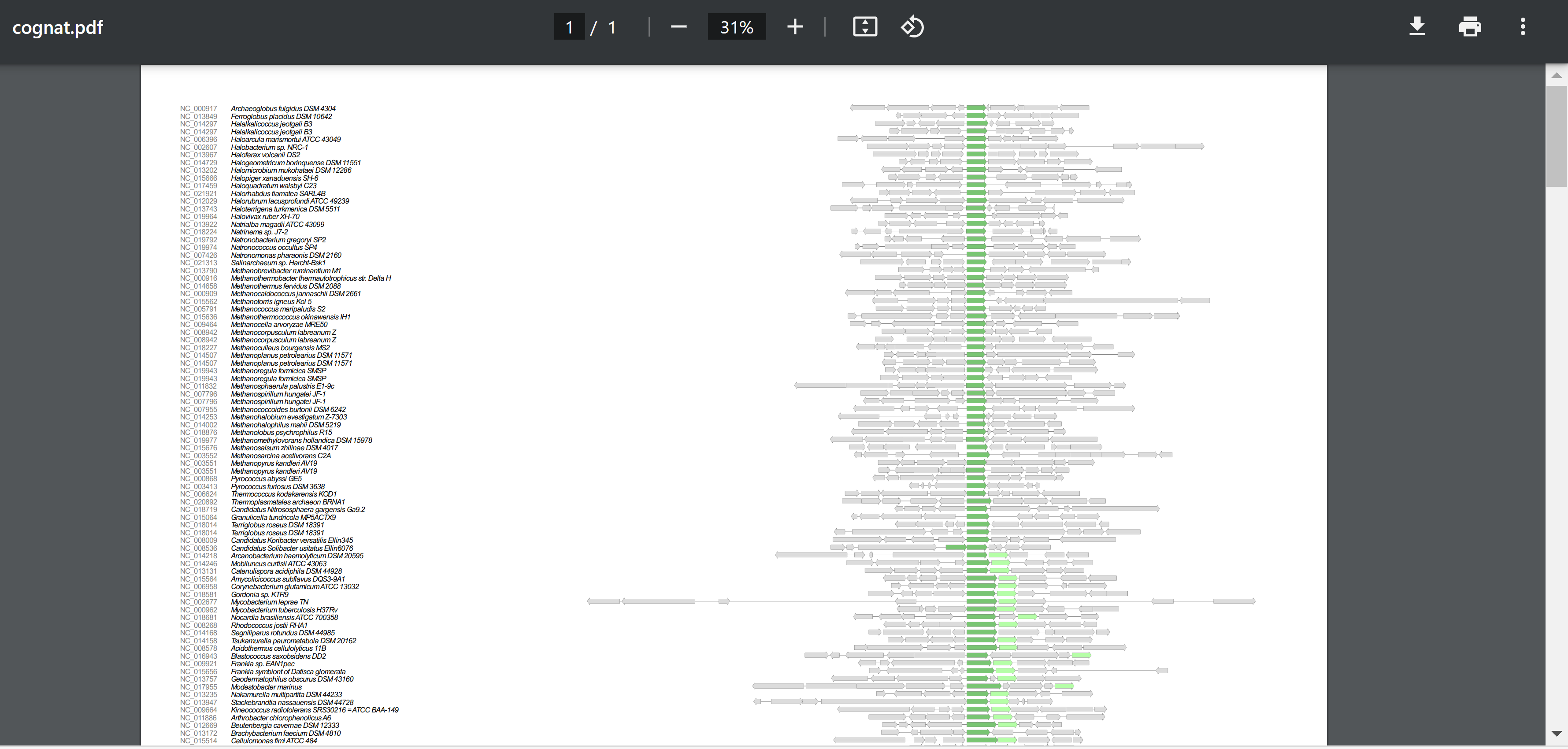

**Figure S11.** SubtiWiki (CoreWiki) beta feature of the database’s Genomic Neighborhood Comparison viewer of the DUF34 homolog of B. subtilis, YqfO (BSU_25170).

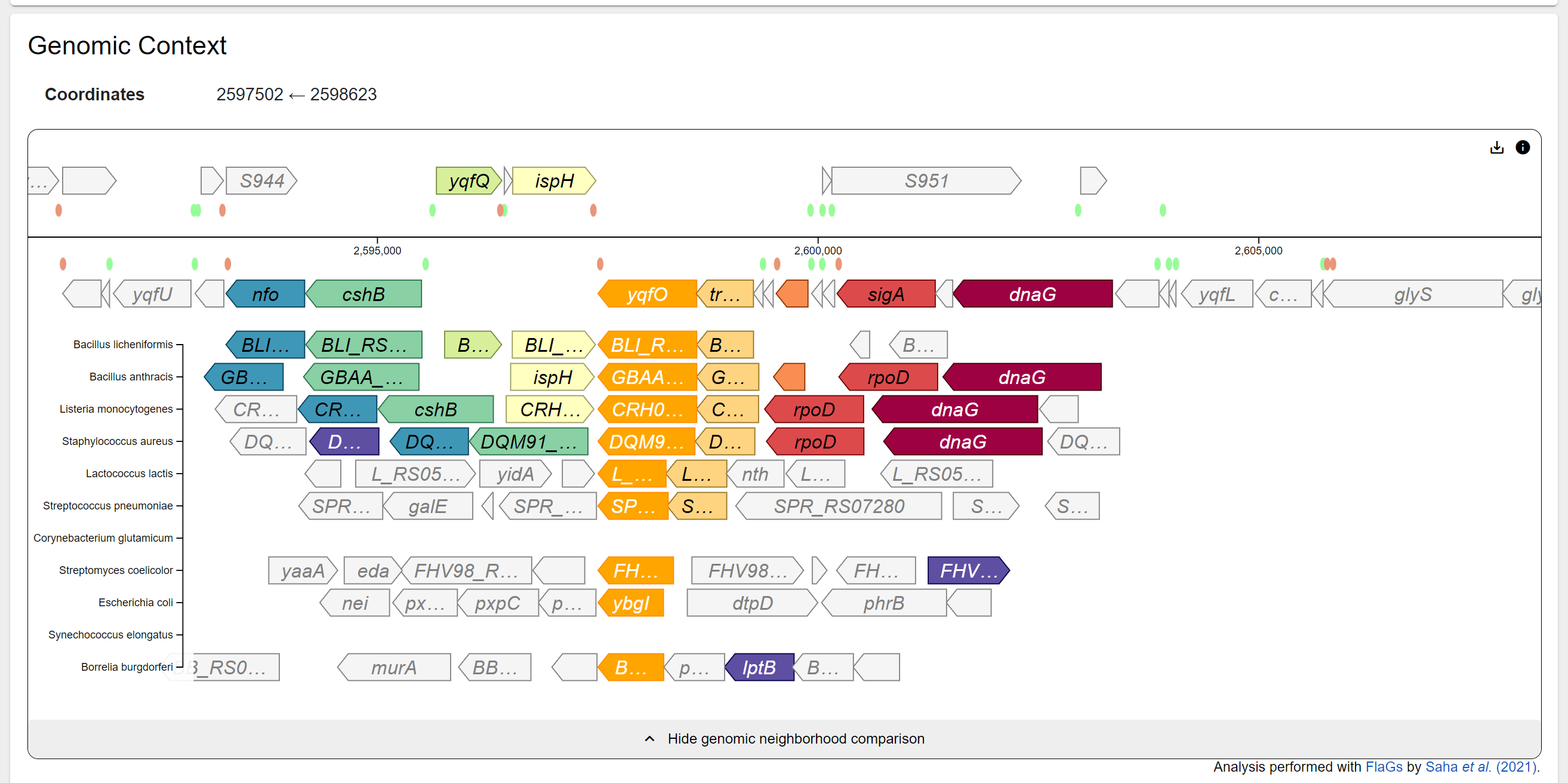

**Figure S12.** Example tree output of MicrobesOnline.

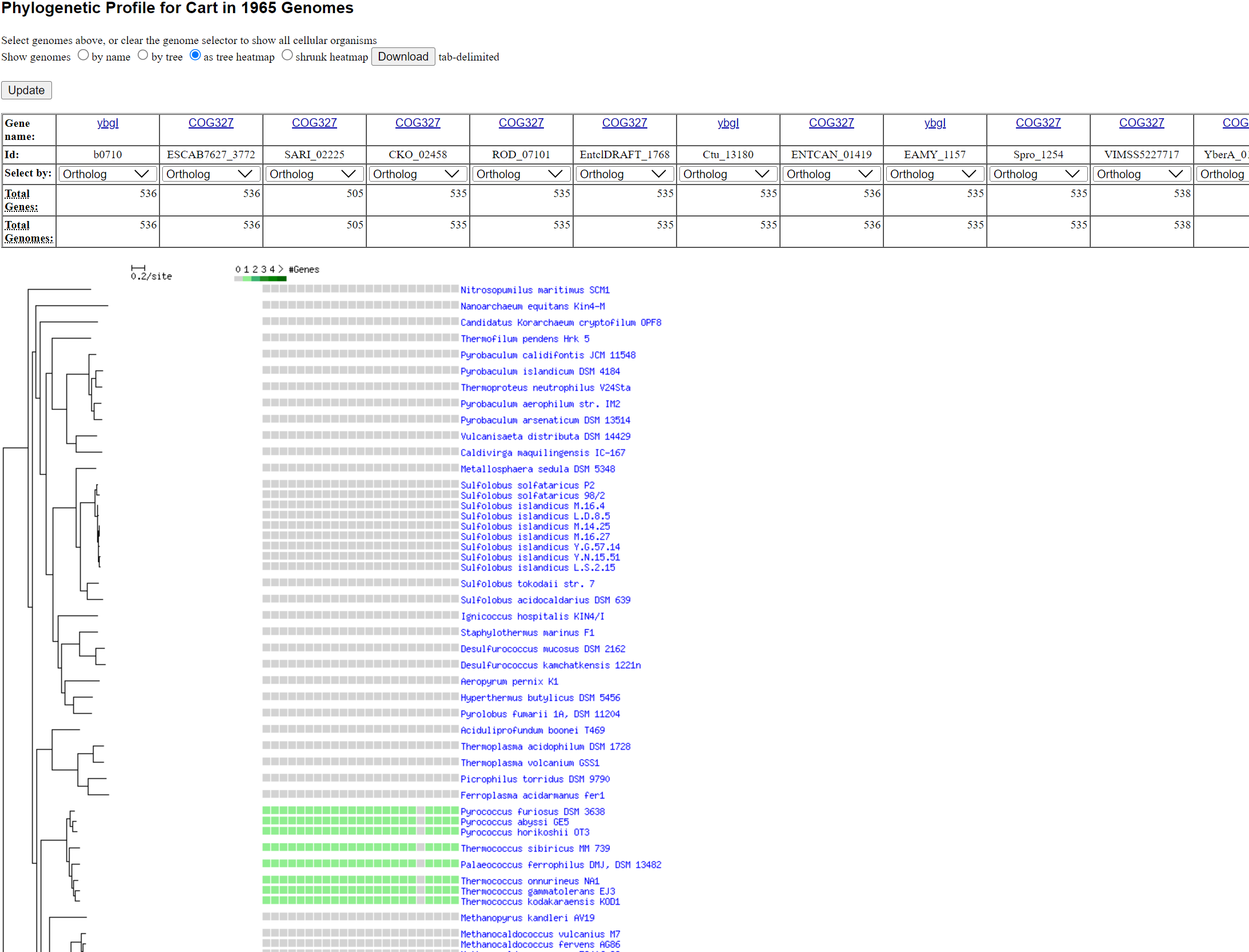

**Figure S13.** Screengrab demonstrating different “filters” for target family recognition, particularly used differentially with multiple targets (MicrobesOnline).
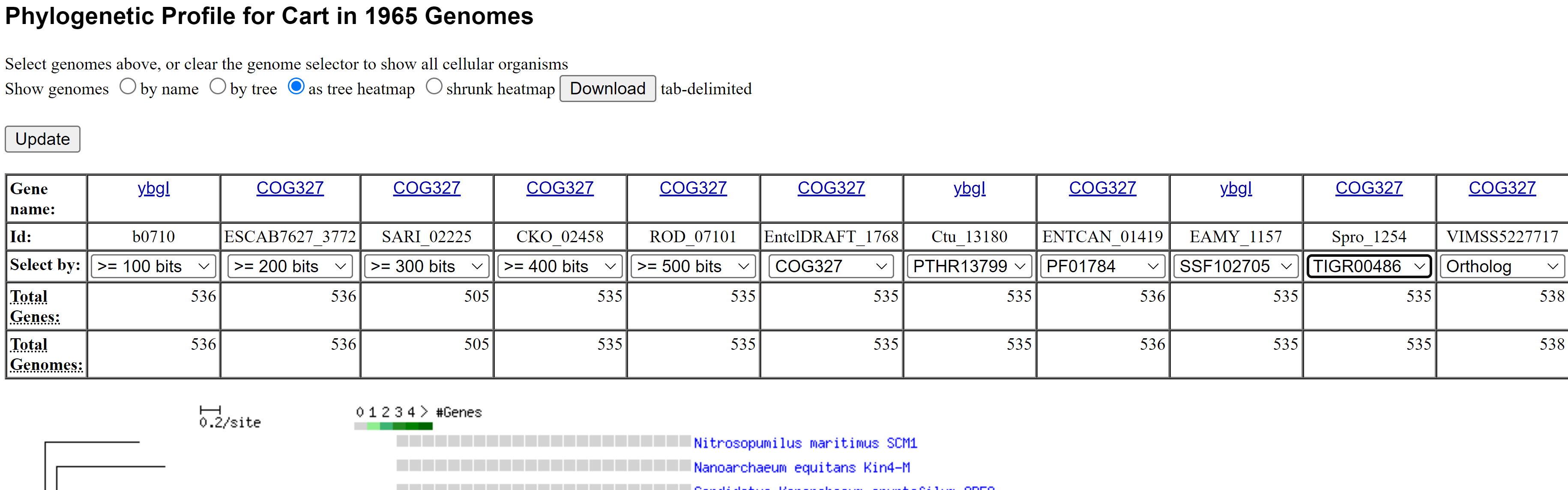

**Figure S14.** Screengrab demonstrating navigation to family-specific trees in MicrobesOnline via the “Gene Info” tab of each entry.

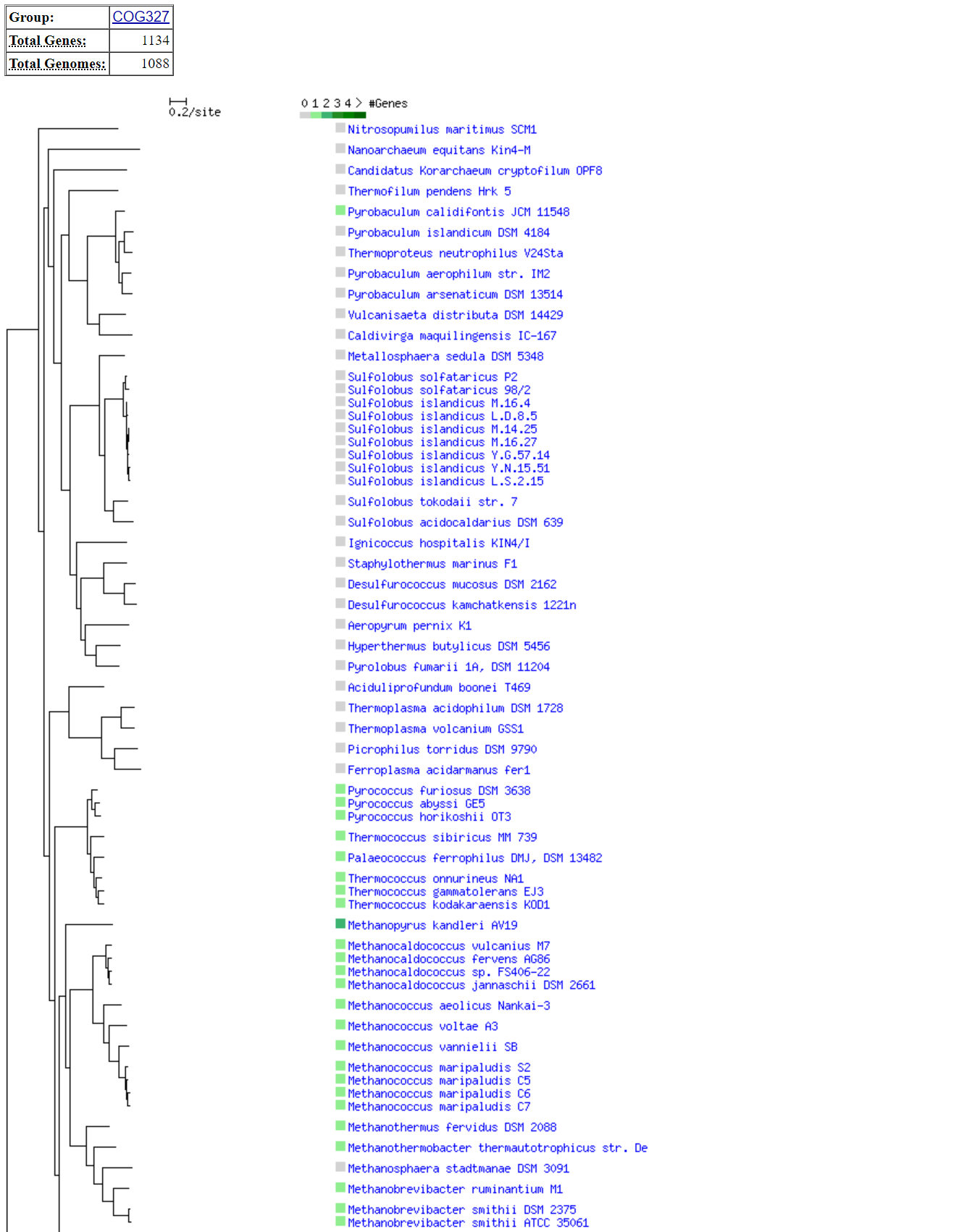

**Figure S15.** “Families” (COGs) in STRING search tool options.

**a.** Input, multiple protein names (release, 12.0)

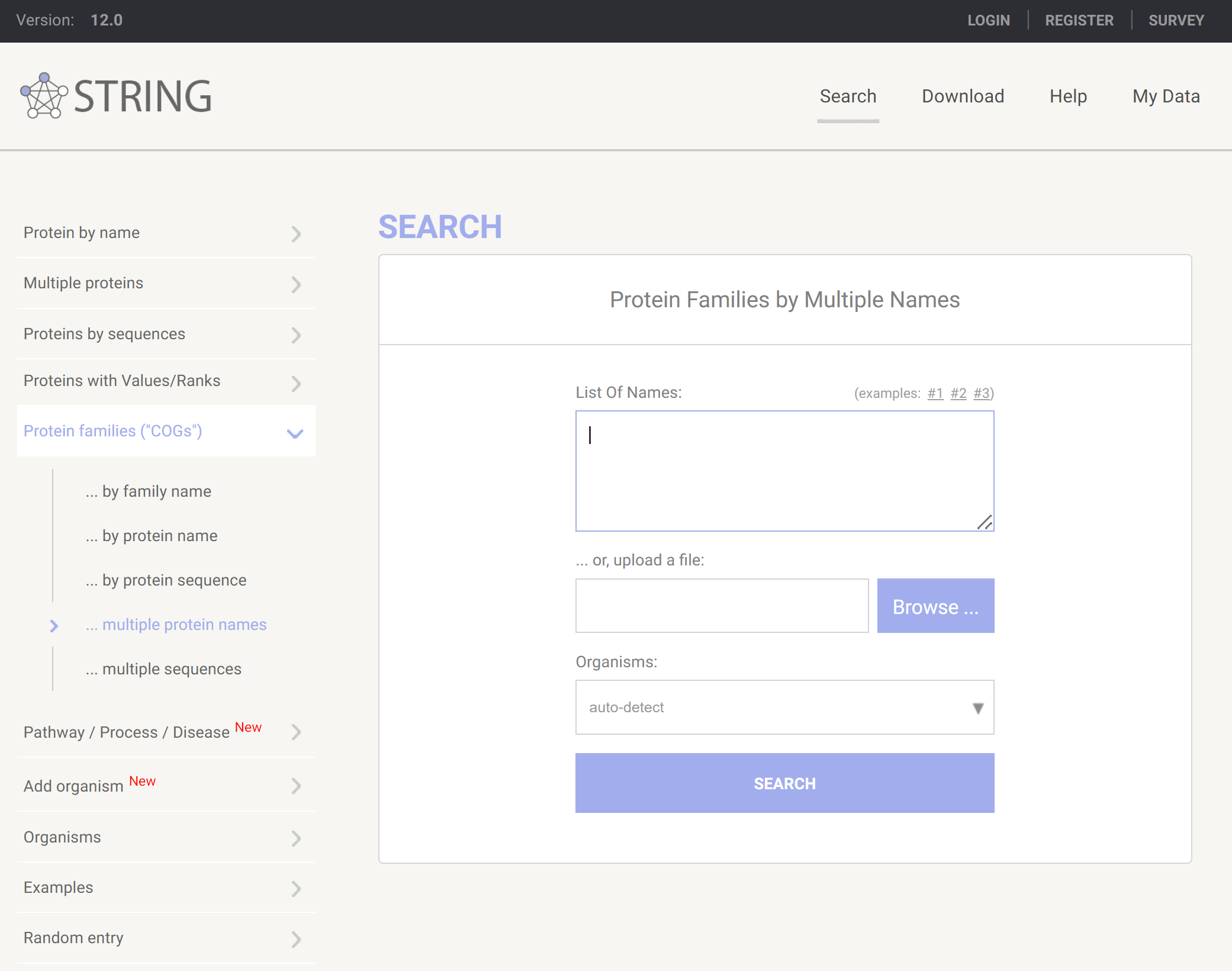

**b.** STRING network output for the physically-interacting proteins DUF34 family protein P9WFM1 (COG0327) and P9WLH3 (COG1579) of *Mycobacterium tuberculosis* (strain ATCC 25618 / H37Rv) via “multiple family names” type query; high-confidence (0.700) minimum required interaction score; no more than 50 interactors, max number of interactors to show for first shell; no more than 5 interactors, max number of interactions to show for second shell.

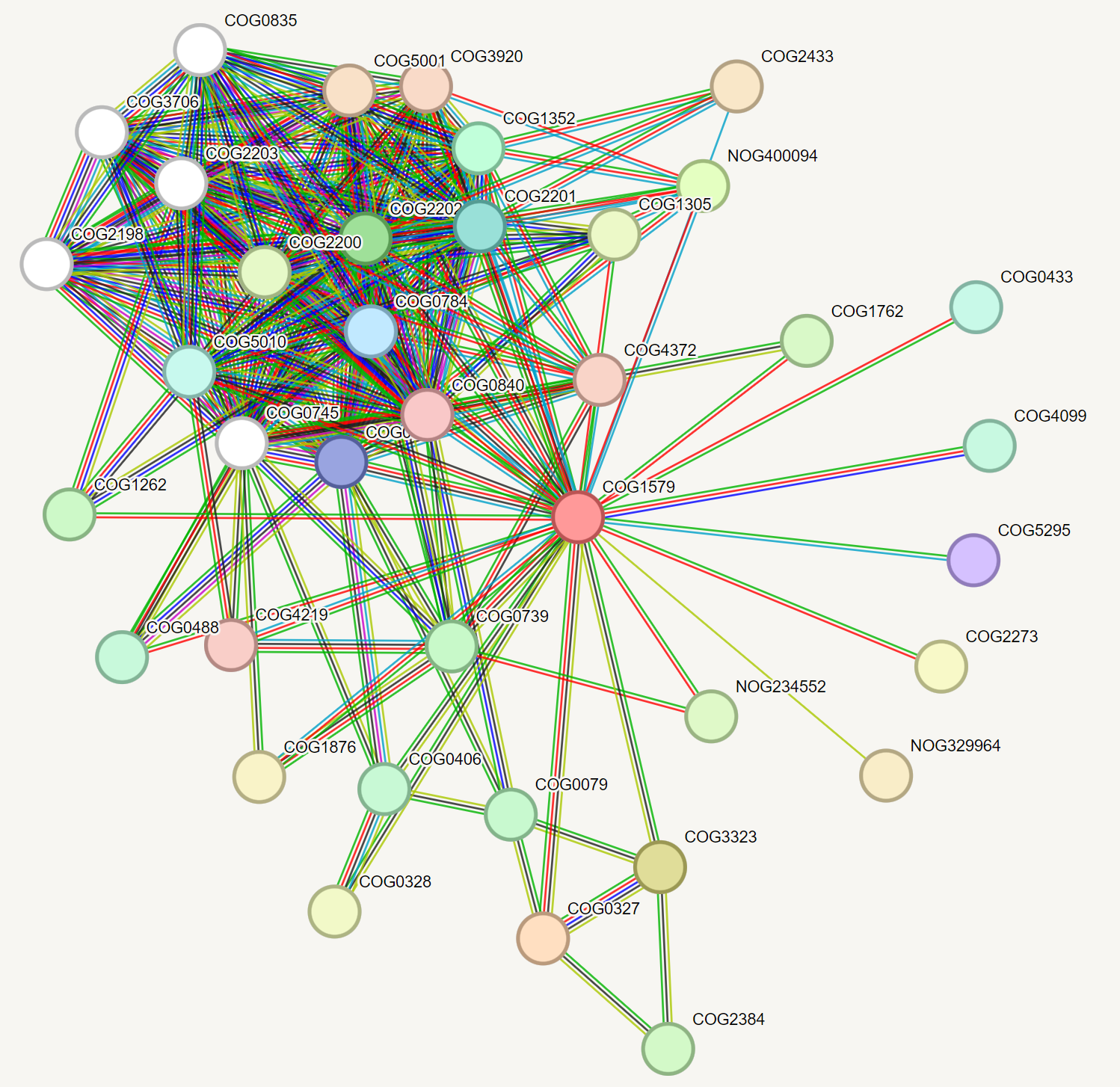

**c.** STRING output for the co-occurrence view for COG0327 (DUF34 family homolog, P9WFM1) and COG2072 (P9WLH3) (phylogeny collapses/expansions that are shown were generated by default).

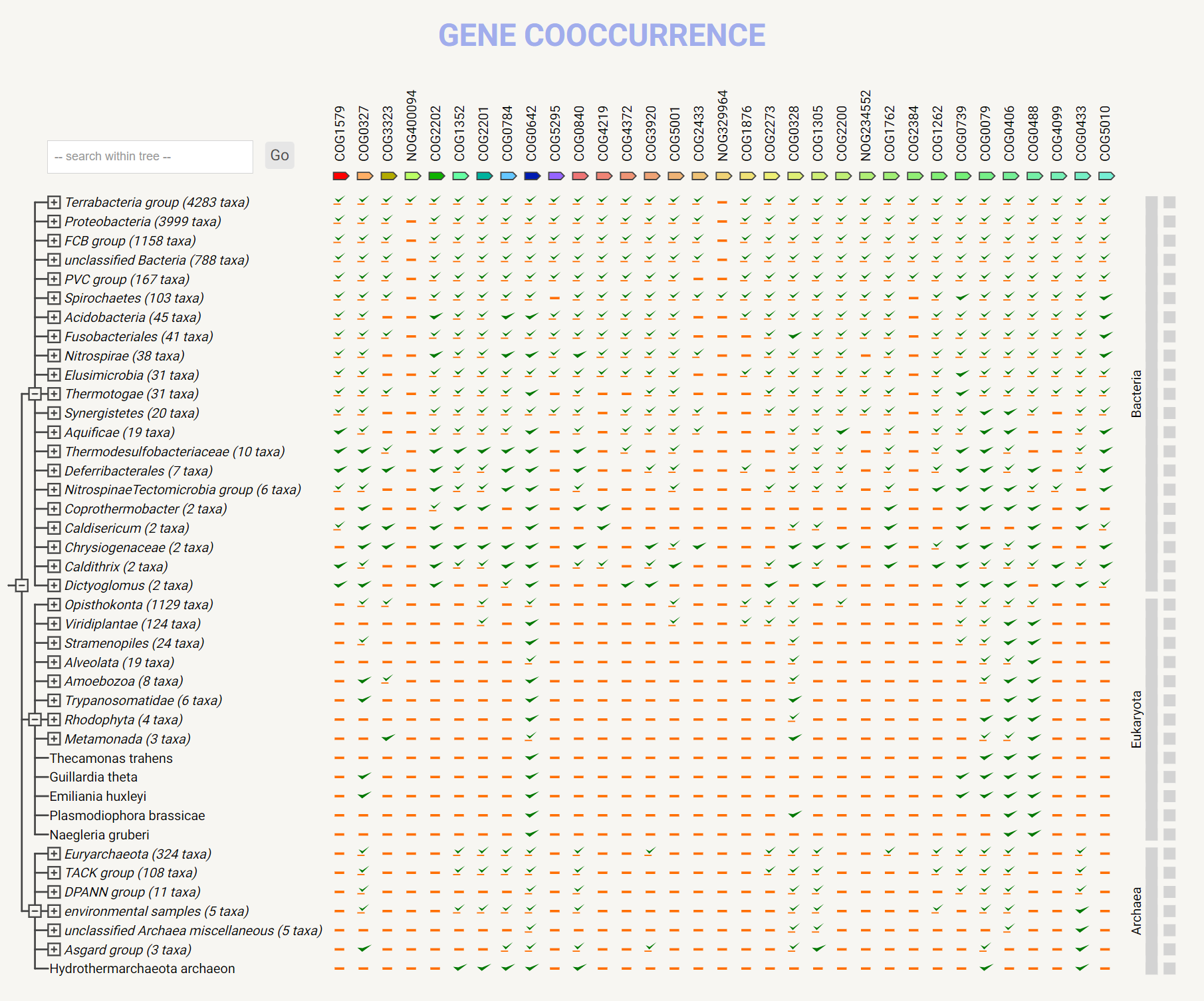

**Figure S16.** KO identifiers being supplied to KEGG orthology tool (multiple families, limited genome customization).

**a.** Output for TsaE (K06925, ubiquitous protein family) and COG1579 (K07164) in KEGG Orthology K number search.

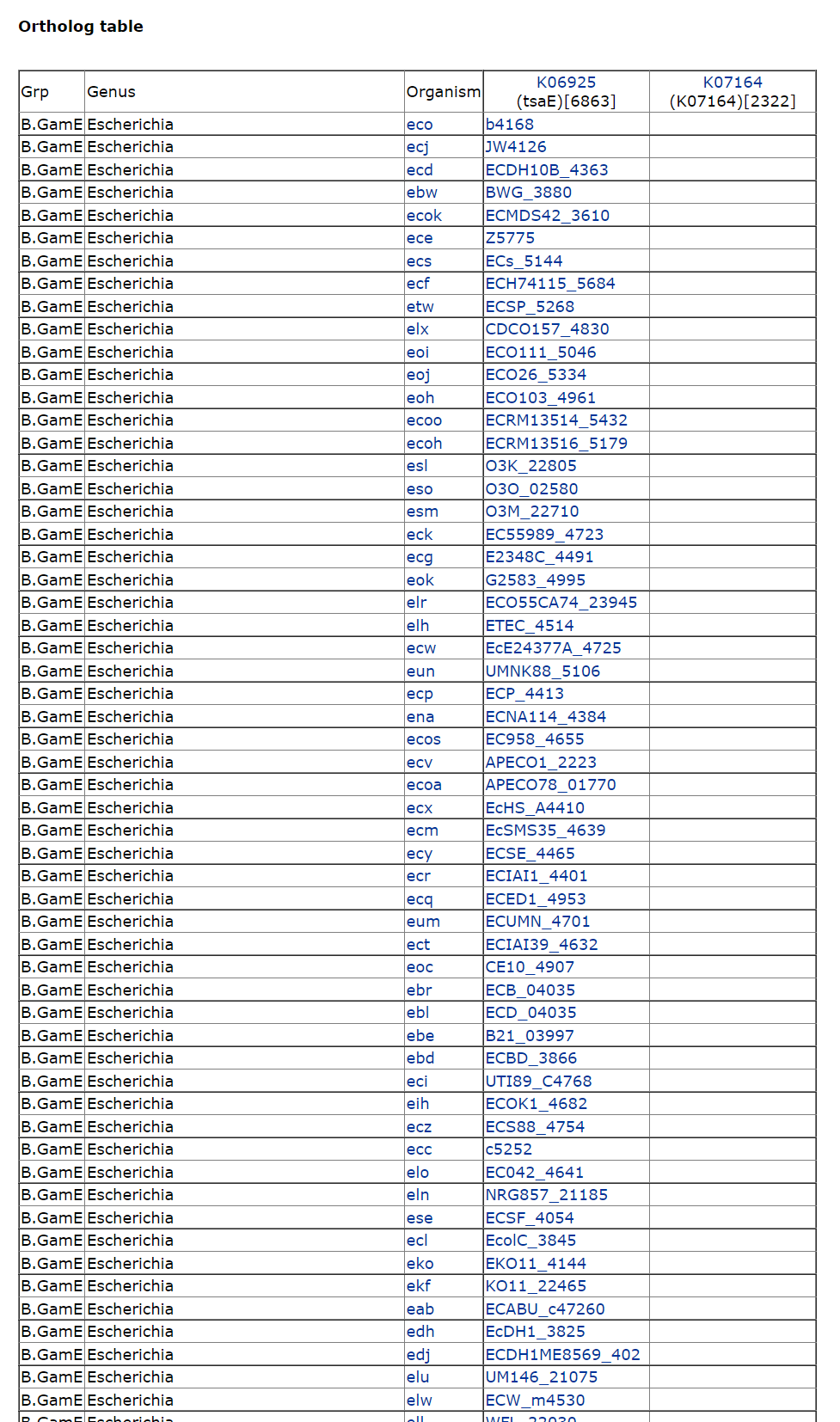

**b.** Input for KEGG Orthology K number search (ortholog table).

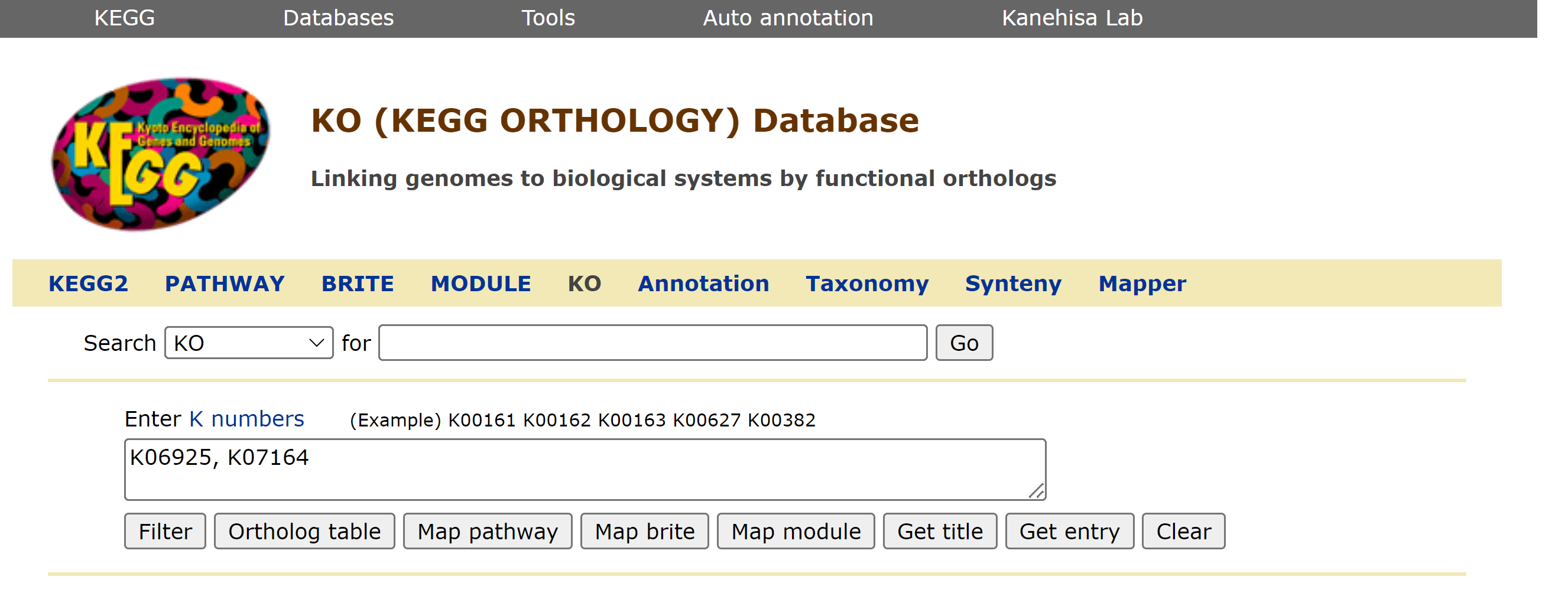

**Figure S17.** Example CAGECAT output using DUF34 and COG1579 homolog sequences: MSMEG_4307 (NCBI-ProteinID: ABK73705); MSMEG_4306 (NCBI-ProteinID: ABK70599). This figure was generated after using cblaster then clinker applications as part of a tandem suite of tools, namely the “compared to query” visualization.

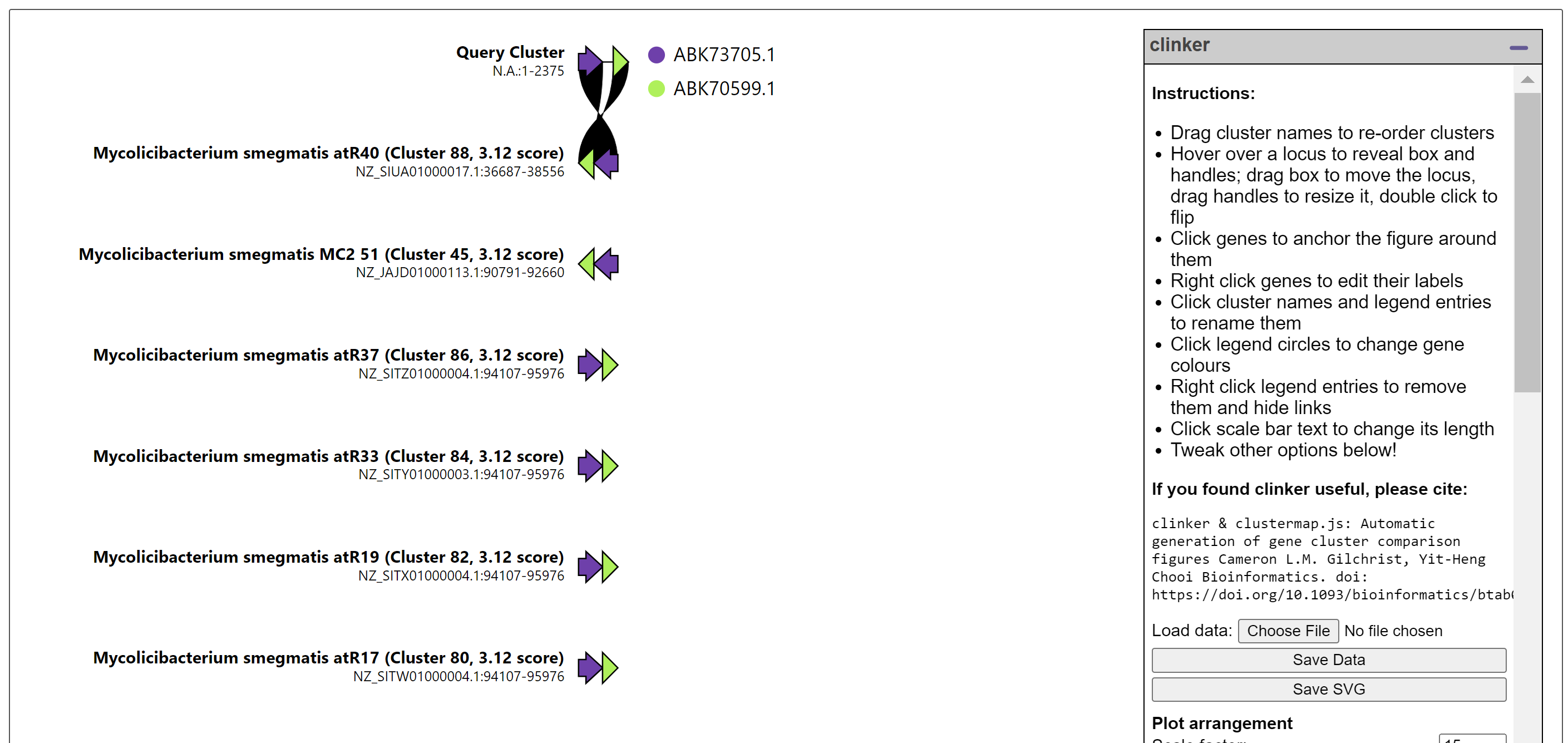

**Figure S18.** Example output for FunCoup using “YbgI” of *E. coli*. While other interaction/correlational data is present, operons are included in each subset.

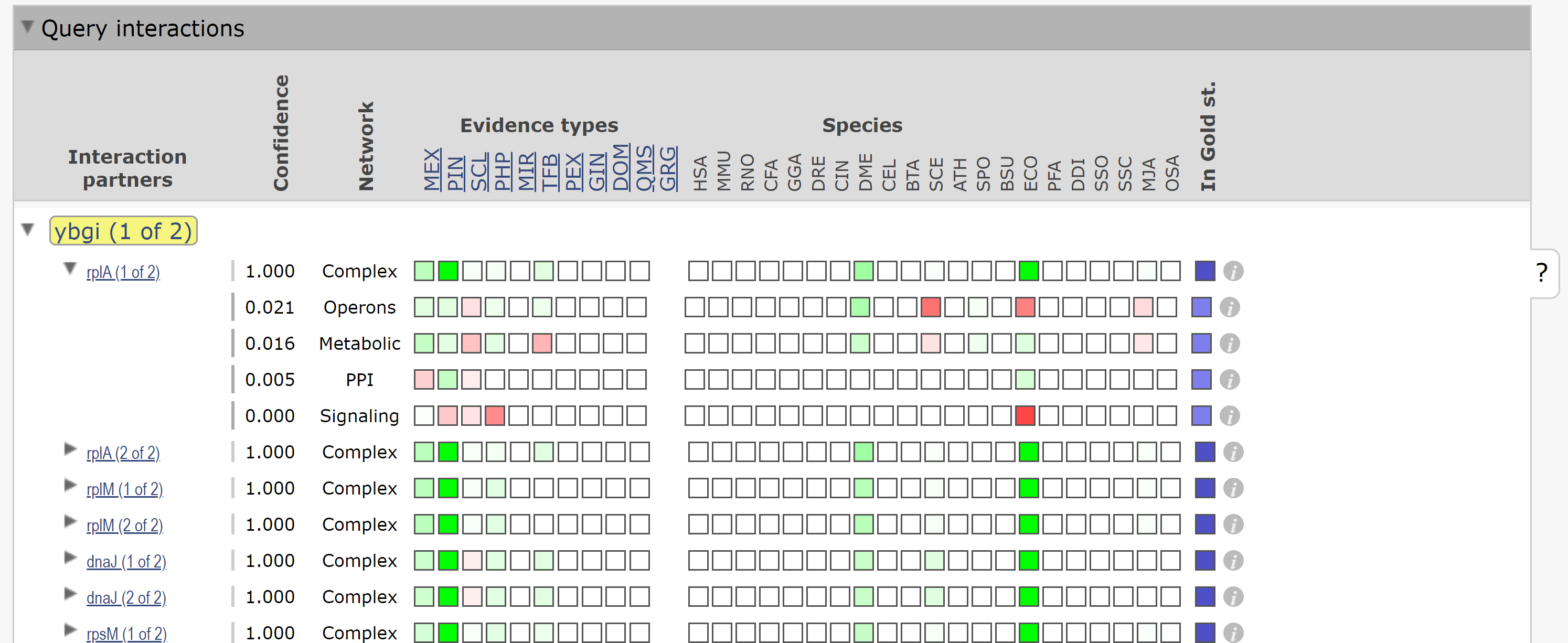

**Figure S19.** BV-BRC protein family sorter, general output example (a) with additional viewer examples: b) subsystems; c) pathways – KEGG Map; d) pathways – Heatmap; e) families.

**a. Initial Output Files (all)**

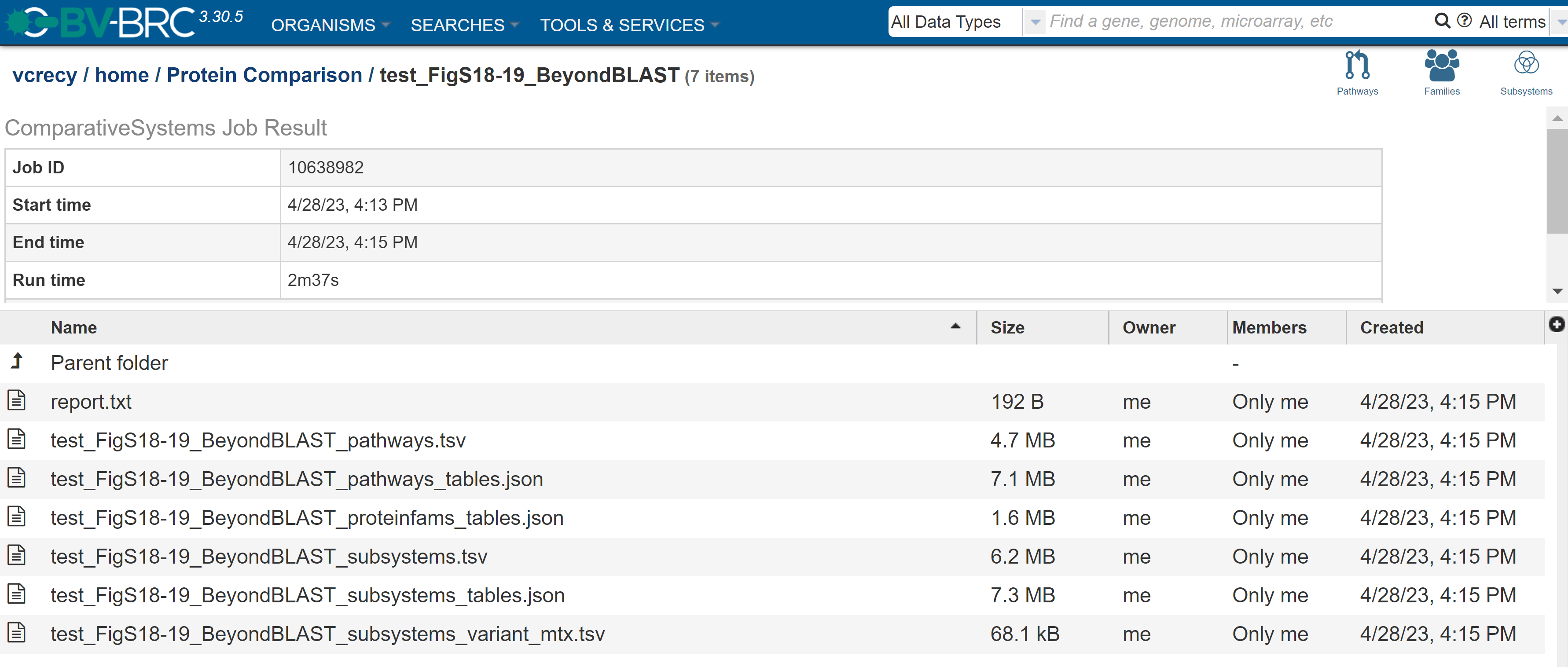

**b. Subsystems (overview).** Subsystems (via the “Subsystems” tab adjacent to the “Subsystems Overview” tab) can then be selected individually or together (>1) for either mapping for the former case or grouping for the latter.

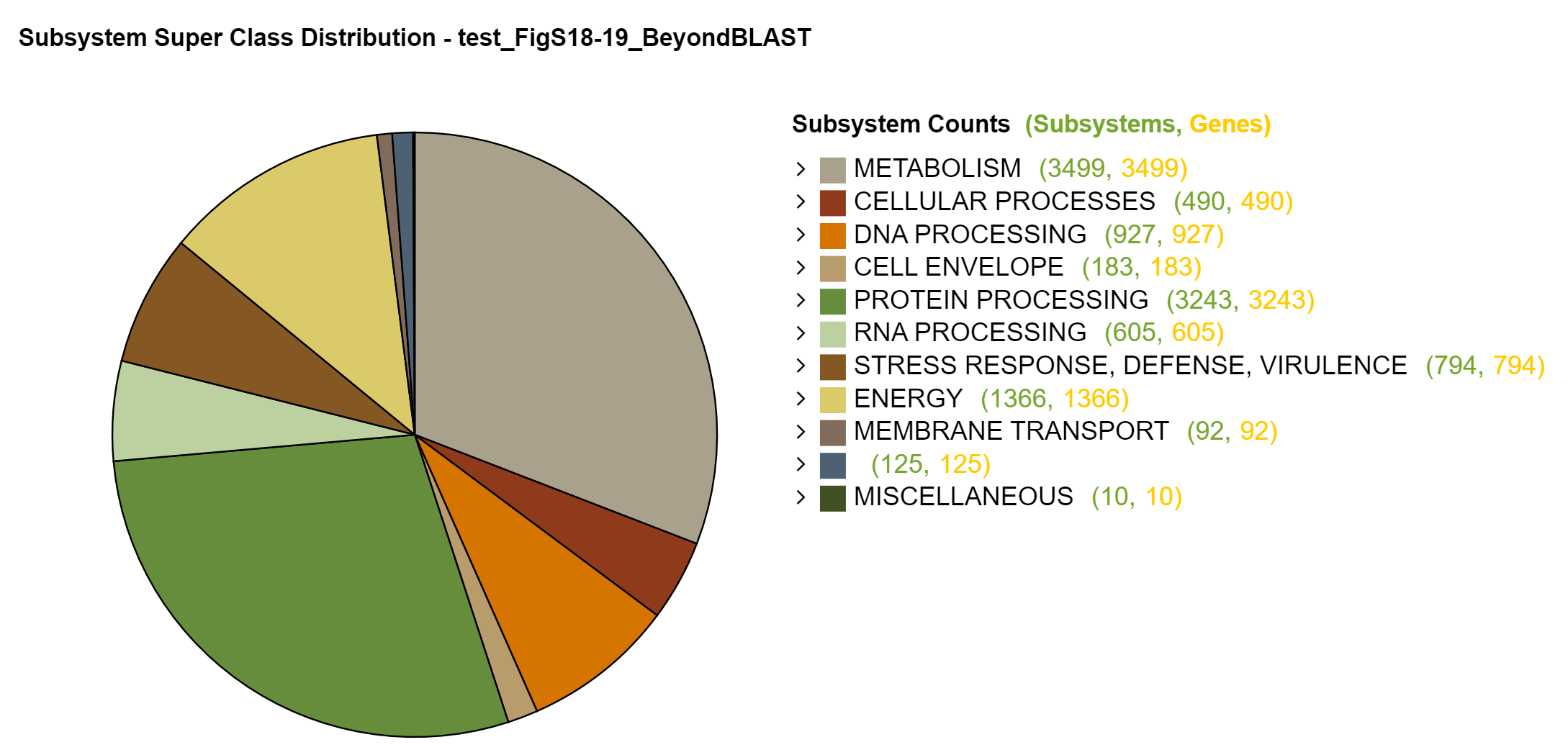

**c. Pathways – KEGG Map**

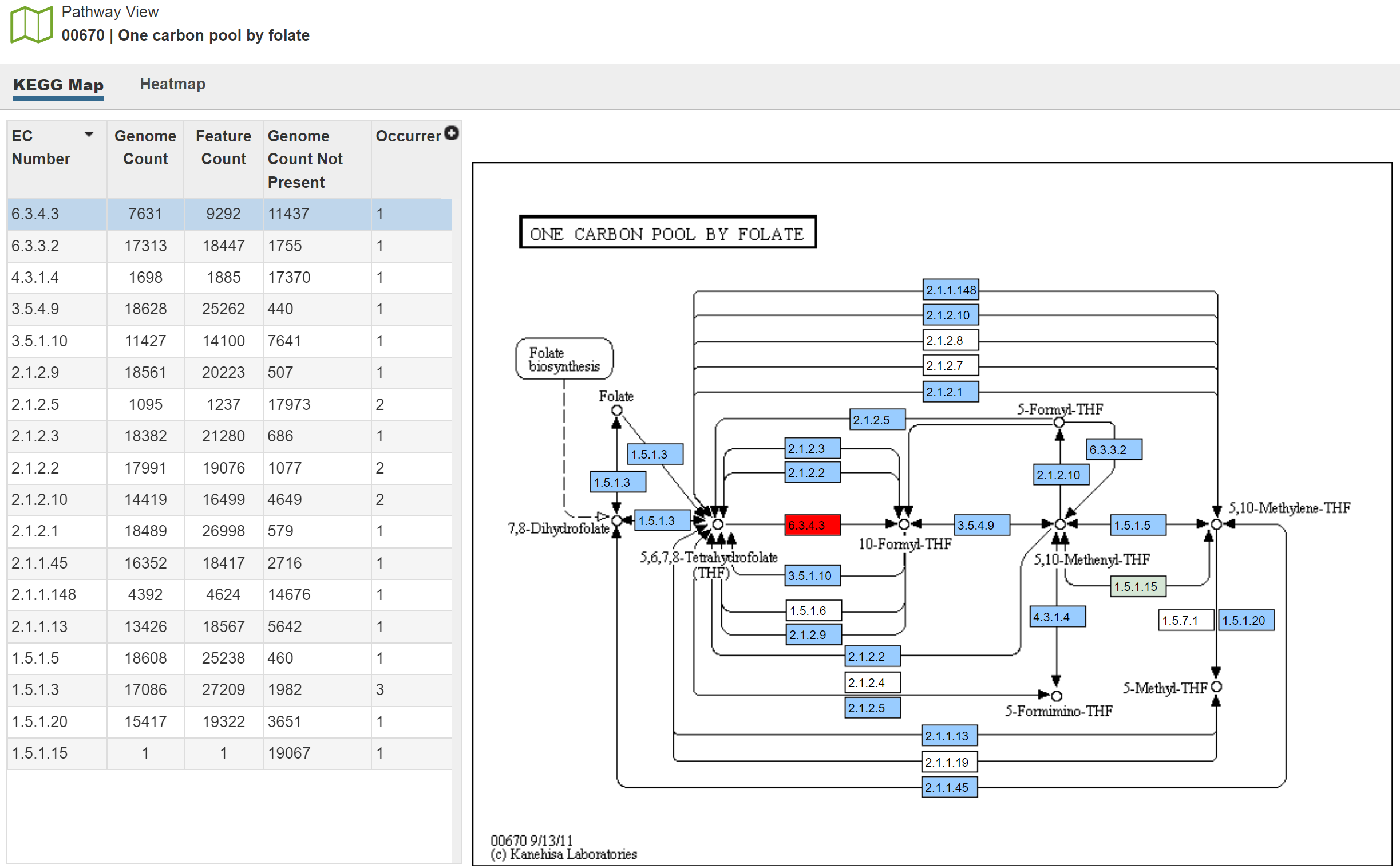

**d. Pathways – Heatmap**

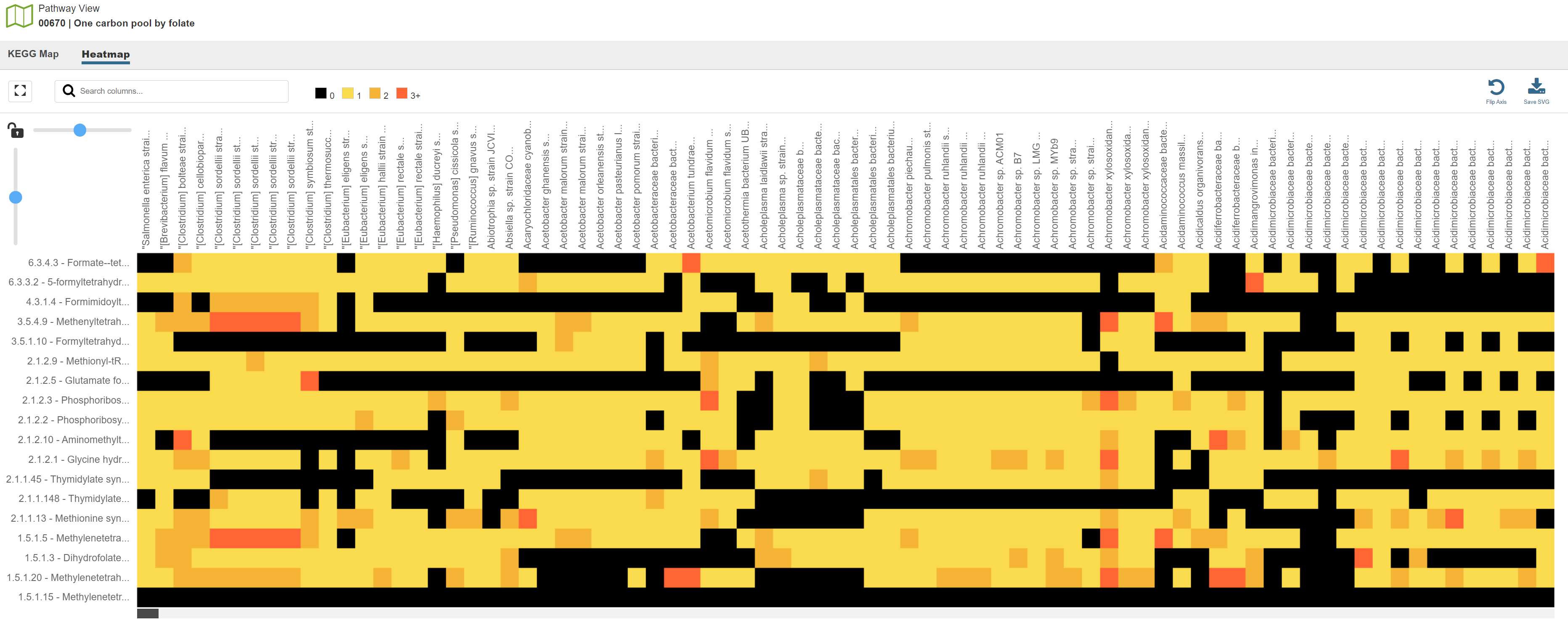

**e. Families (output)**

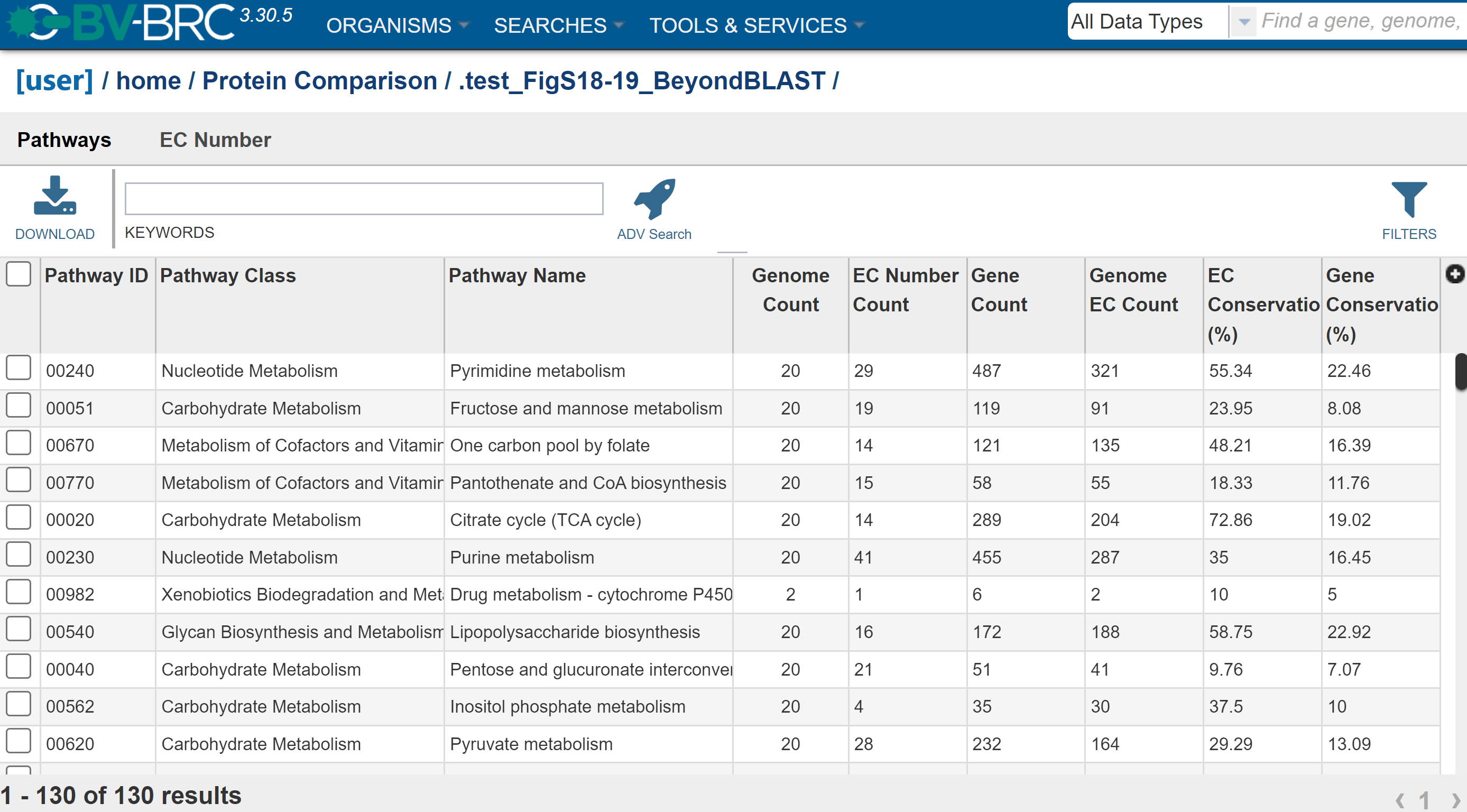

**Figure S20.** BV-BRC protein family sorter output, Families.

**a.** Heatmap view of Families output. Green boxes highlight the text entry box and the heatmap tab selection. Text box provides search ability that is dependent upon annotation quality in database.
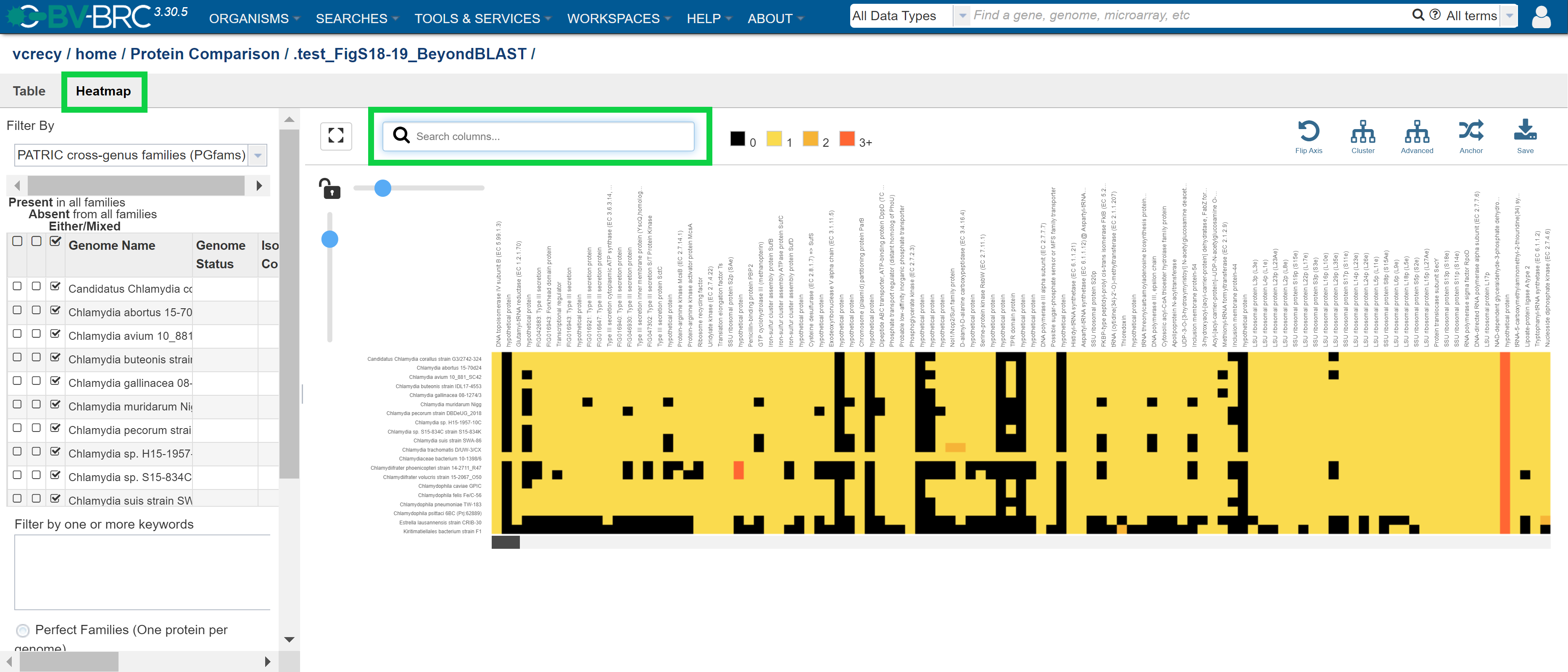

**b.** Table view of Families output. Purple boxes highlight the multiple-select checkboxes that a user may select for viewing in the accompanying heatmap or for export/grouping.

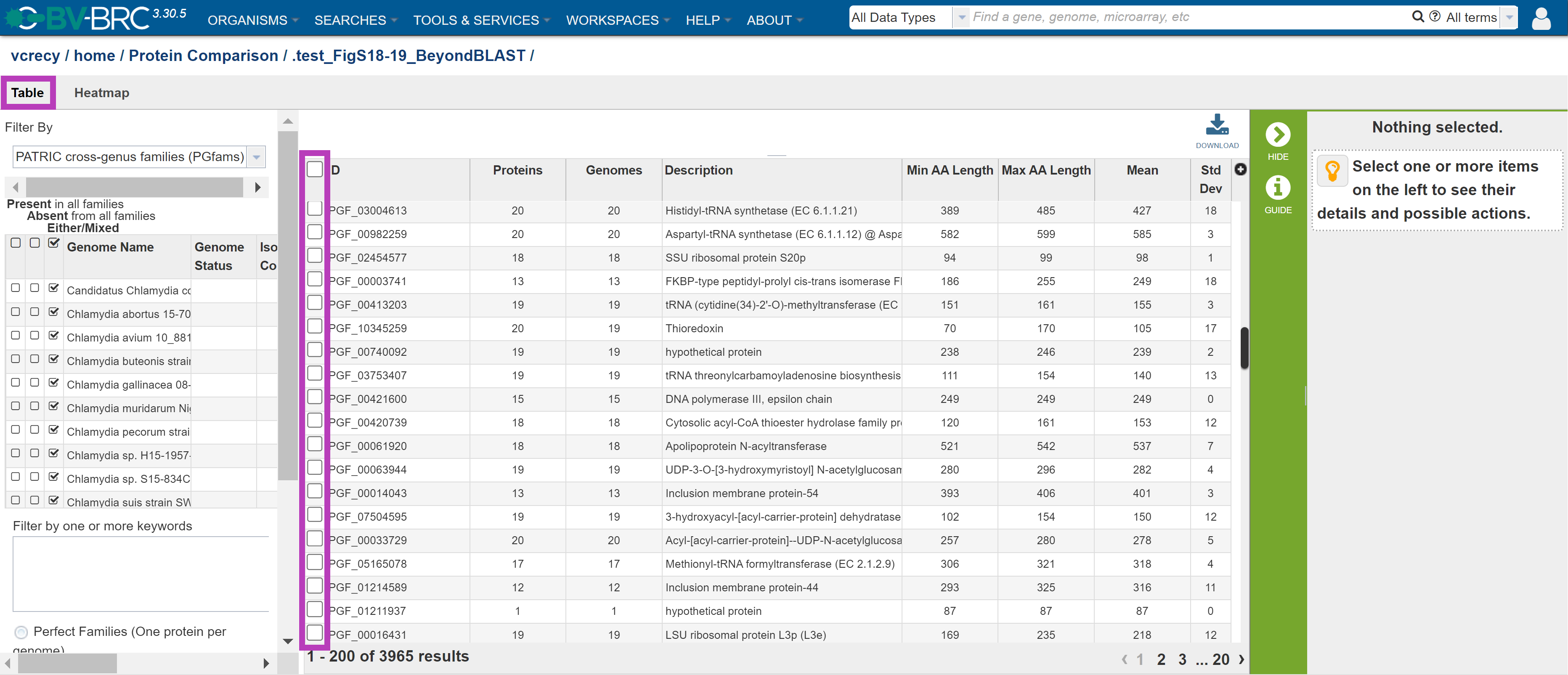

**Figure S21.** EggNOG Phylogenetic Profile tool example output using the site-provided example input data. At the time of writing, the tool seemed to be experiencing loading and/or query processing errors, failing to generate the intended product.

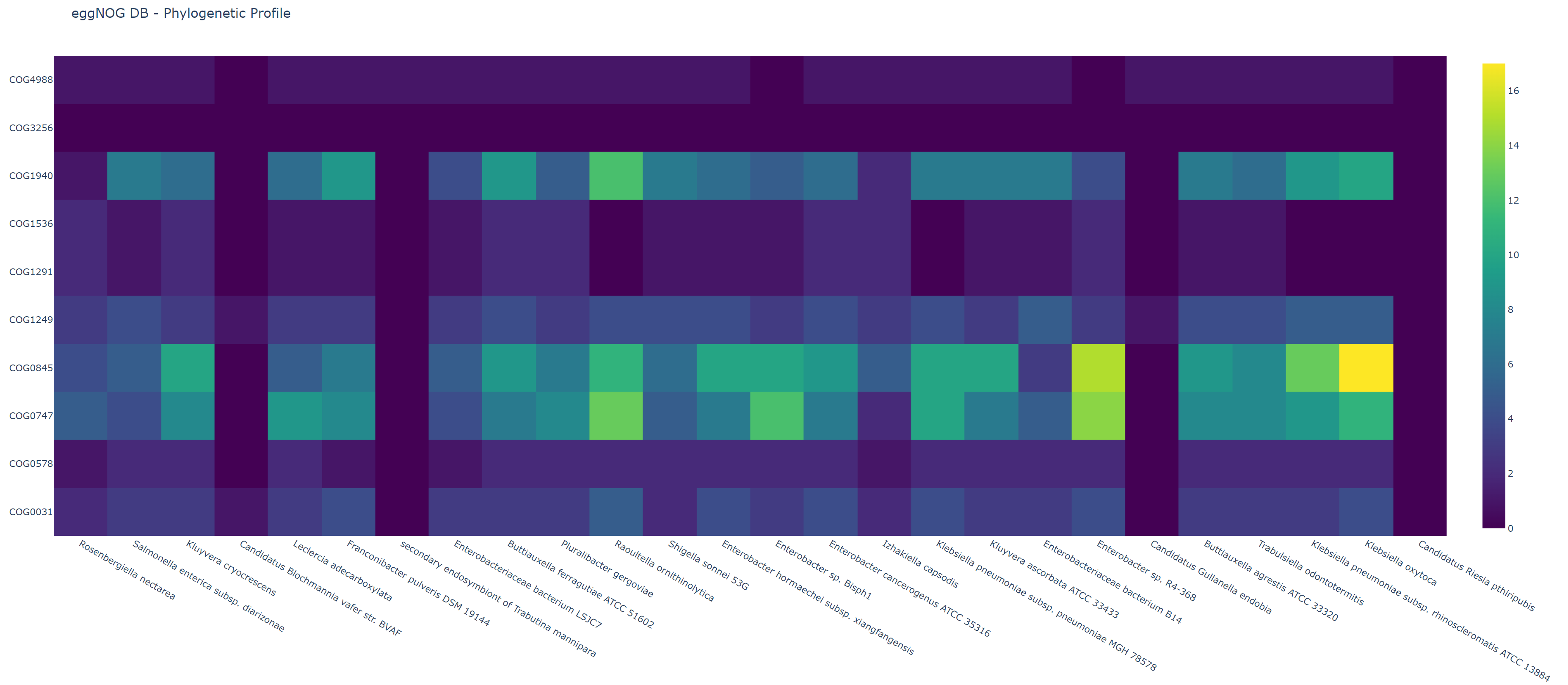
.

**Figure S22.** KBase workflow for generating a “gene tree” (as regarded by KBase) from several select genomes. Figure was directly adapted from a KBase 2020 phylogenomics online workshop diagram. Circles indicate data for input/outputs. Squares indicate applications within the KBase *“narrative”* (i.e., “narratives” in KBase are curated workflows of applications, as well as respective inputs and outputs [data objects], that can be created by users or, alternatively, made available to users by developers for implementation as example templates for common workflows).

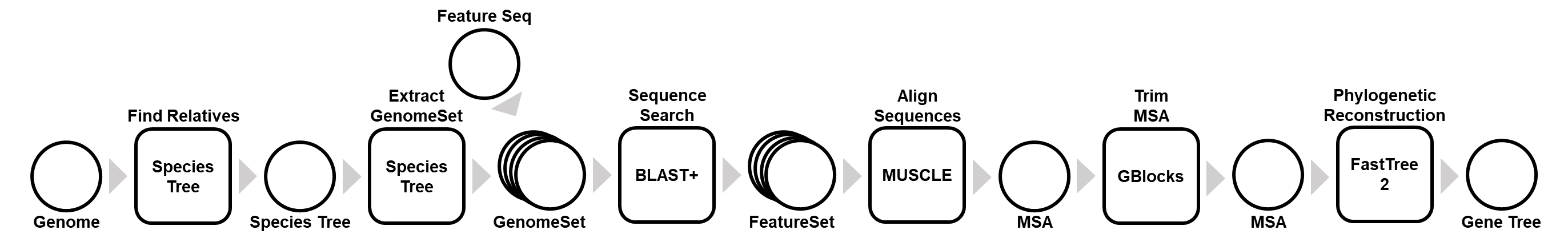

**Figure S23.** Genomic Context Visualizer (GeCoViz)

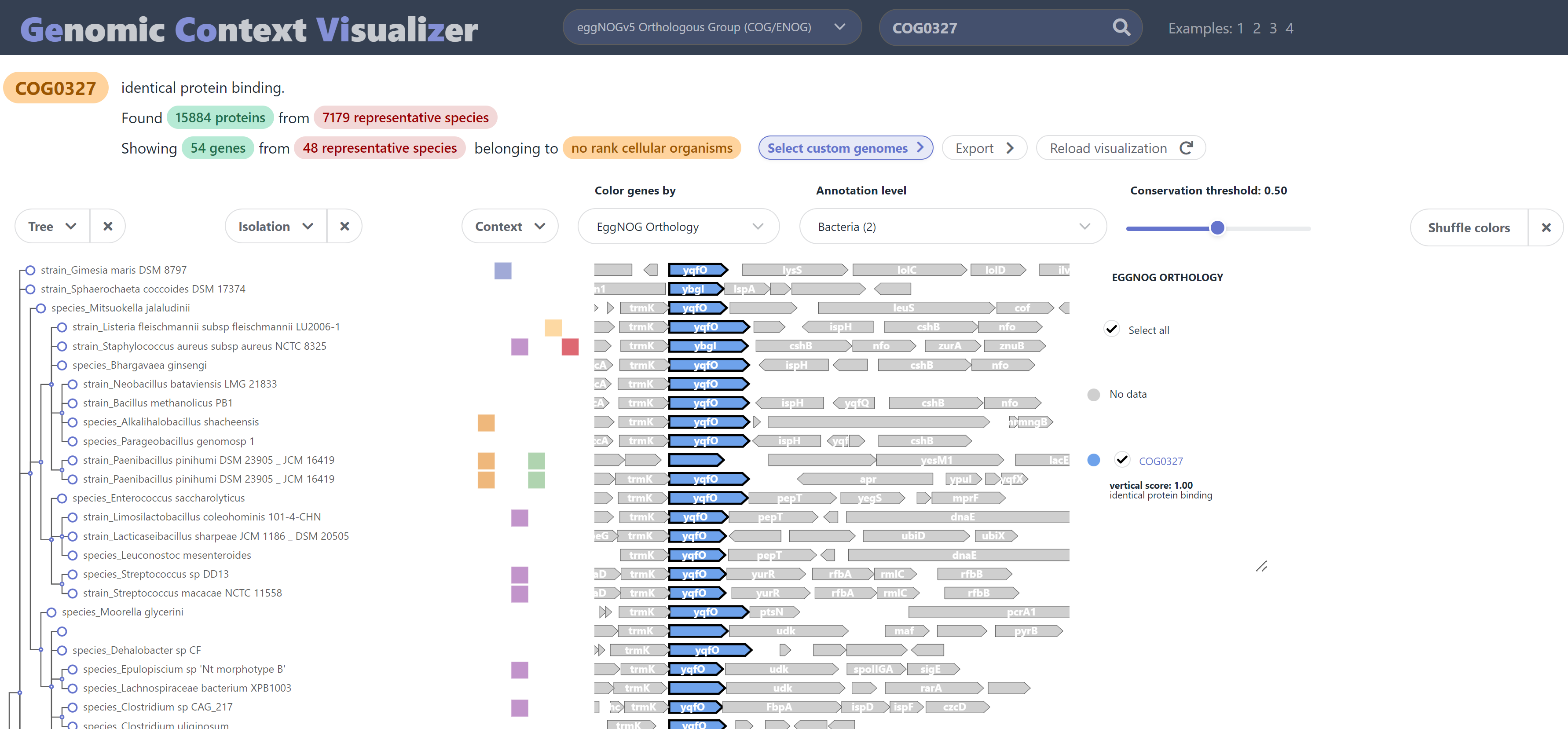

**Figure S24.** GizmoGene example output for the DUF34 homolog of *Mycobacterium tuberculosis* H37Rv_CG (BV-BRC feature ID: fig|1773.25616.peg.2443) using a custom set of BV-BRC genomes (62 Representative genomes, bacteria).

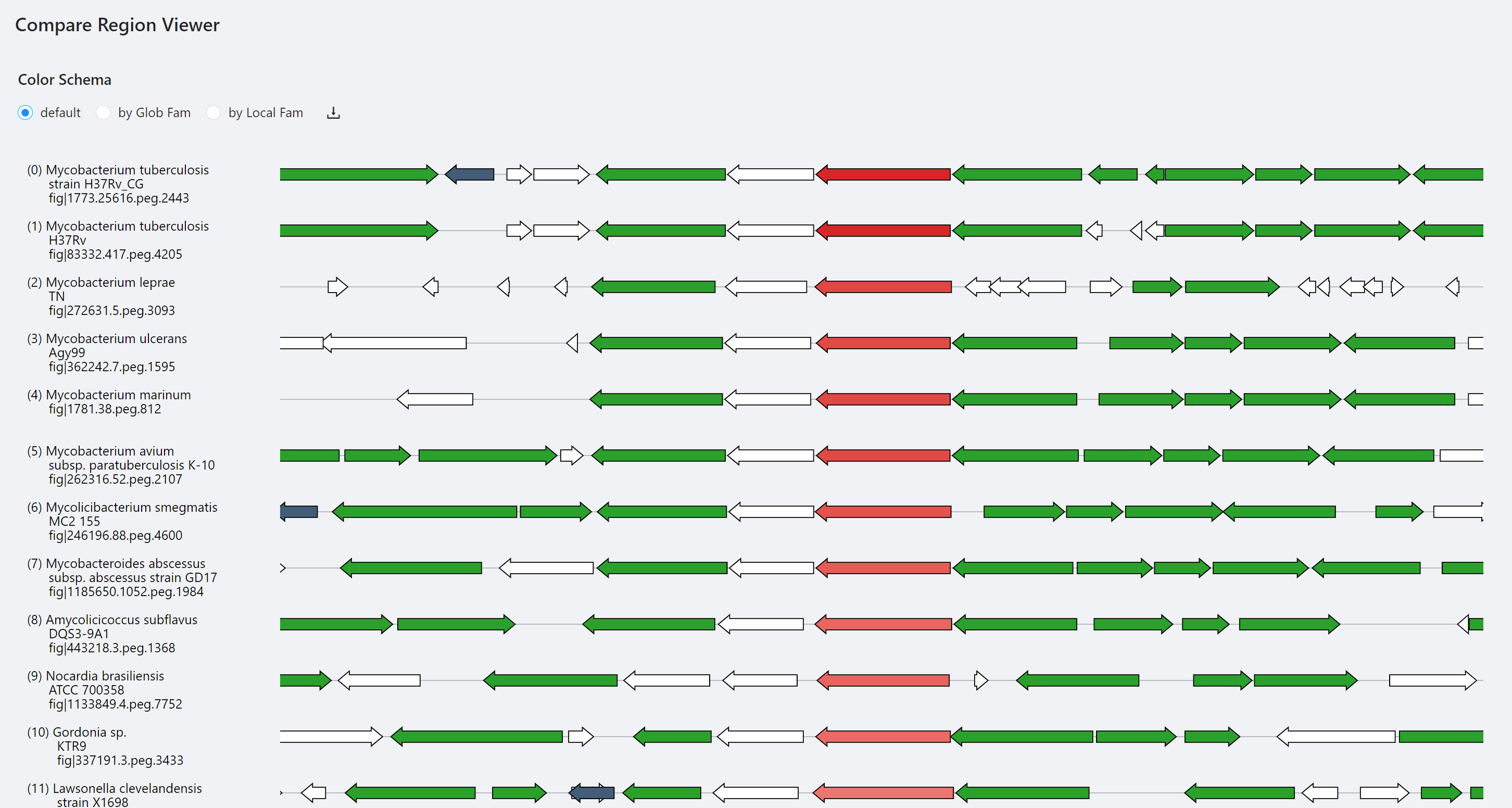
